# Resource Profile and User Guide of the Polygenic Index Repository

**DOI:** 10.1101/2021.05.08.443158

**Authors:** Joel Becker, Casper A.P. Burik, Grant Goldman, Nancy Wang, Hariharan Jayashankar, Michael Bennett, Daniel W. Belsky, Richard Karlsson Linnér, Rafael Ahlskog, Aaron Kleinman, David A. Hinds, 23andMe Research Group, Avshalom Caspi, David L. Corcoran, Terrie E. Moffitt, Richie Poulton, Karen Sugden, Benjamin S. Williams, Kathleen Mullan Harris, Andrew Steptoe, Olesya Ajnakina, Lili Milani, Tõnu Esko, William G. Iacono, Matt McGue, Patrik K.E. Magnusson, Travis T. Mallard, K. Paige Harden, Elliot M. Tucker-Drob, Pamela Herd, Jeremy Freese, Alexander Young, Jonathan P. Beauchamp, Philipp Koellinger, Sven Oskarsson, Magnus Johannesson, Peter M. Visscher, Michelle N. Meyer, David Laibson, David Cesarini, Daniel J. Benjamin, Patrick Turley, Aysu Okbay

## Abstract

Polygenic indexes (PGIs) are DNA-based predictors. Their value for research in many scientific disciplines is rapidly growing. As a resource for researchers, we used a consistent methodology to construct PGIs for 47 phenotypes in 11 datasets. To maximize the PGIs’ prediction accuracies, we constructed them using genome-wide association studies—some of which are novel—from multiple data sources, including 23andMe and UK Biobank. We present a theoretical framework to help interpret analyses involving PGIs. A key insight is that a PGI can be understood as an unbiased but noisy measure of a latent variable we call the “additive SNP factor.” Regressions in which the true regressor is the additive SNP factor but the PGI is used as its proxy therefore suffer from errors-in-variables bias. We derive an estimator that corrects for the bias, illustrate the correction, and make a Python tool for implementing it publicly available.

## Main

The ability to predict complex outcomes from genotype data alone is rapidly increasing. The main catalyst behind the increases is the success of genome-wide association studies ^1^ (GWAS). GWAS estimate the relationship between a trait, called a “phenotype,” and each of millions of genetic variants. The “summary statistics” (coefficients and standard errors) from GWAS can be used to construct a DNA-based predictor of the phenotype, calculated essentially as a coefficient-weighted sum of allele counts ^2,3^. There are a variety of terms used for such DNA-based predictors. In this paper, we will refer to them as “polygenic indexes” (see Box).

#### Box. Note on Terminology

In this paper, we use the term “polygenic index” instead of the commonly used terms “polygenic score” and “polygenic risk score.” Most of us prefer the term polygenic index because we are persuaded by the argument that it is less likely to give the impression of a value judgment where one is not intended. The term polygenic index was first proposed by Martha Minow at a meeting of the Trustees of the Russell Sage Foundation.

As GWAS sample sizes have grown, coefficients are estimated more precisely, enabling the construction of more predictive PGIs. One example is the PGI for educational attainment. The original PGI was constructed from a GWAS of ~100,000 individuals and predicted ~2% of the variance in years of schooling across individuals ^4^. The third and most recent PGI for educational attainment (EA) predicts ~12% of the variance ^5^. Qualitatively similar patterns have been observed in PGIs for other complex-trait phenotypes ^1,6^, including height, fertility, personality traits, and risk of many common diseases.

PGIs became mainstream in human genetics remarkably quickly. While predictive genetic indexes have a long history in plant and animal genetics ^7^, the idea of using GWAS summary statistics to generate a PGI for humans was first proposed in 2007 ^2^. The first study to empirically construct and validate a PGI was a GWAS of bipolar disorder and schizophrenia published in 2009 ^3^. Soon thereafter, command of methods used to construct PGIs became a standard part of the skill repertoire of analysts specializing in genome-wide data.

Today, PGIs are profoundly impacting research across the disciplinary spectrum. In medicine, much of the discussion revolves around their potential use as tools for identifying individuals who could benefit from enhanced screening and preventive therapies ^8^. Though much uncertainty remains about their ultimate clinical utility ^9^, one recent study of polygenic risk for five common diseases concluded that the science is sufficiently far along to contemplate incorporating polygenic prediction into clinical care ^10^. Researchers working at the intersection of the social and natural sciences have articulated visions of how PGIs could be productively leveraged in a number of ways to advance knowledge about important questions ^11–13^. Already, the various iterations of the EA PGI have been used, among other things, to trace out pathways for genetic influences that develop with age ^14^ and through school ^15^, study assortative mating ^16,17^, trace recent migration patterns ^18,19^, and improve analyses of the relationship between education and earnings ^20^. As PGIs become more predictive and available for more phenotypes, potential applications will multiply, and novel areas of research are likely to open up.

To depict the rapid growth in research using PGIs, Figure 1 shows the percentage of PGI-related papers presented at the annual meetings of the Behavior Genetics Association. The percentage increased from zero in 2009 to 20% in 2019. The figure also shows how the percentages of papers classified as candidate-gene studies and twin/family/adoption studies—two other commonly used approaches—have evolved over time. The declining fraction of candidate-gene studies in the figure is consistent with the hypothesis of a paradigm shift, with candidate-gene-based approaches gradually being displaced by PGI-based approaches ^13^. This shift occurred, at least in part, because PGIs are not subject to some well-known methodological limitations of candidate-gene studies ^21–23^.

**Figure 1:**
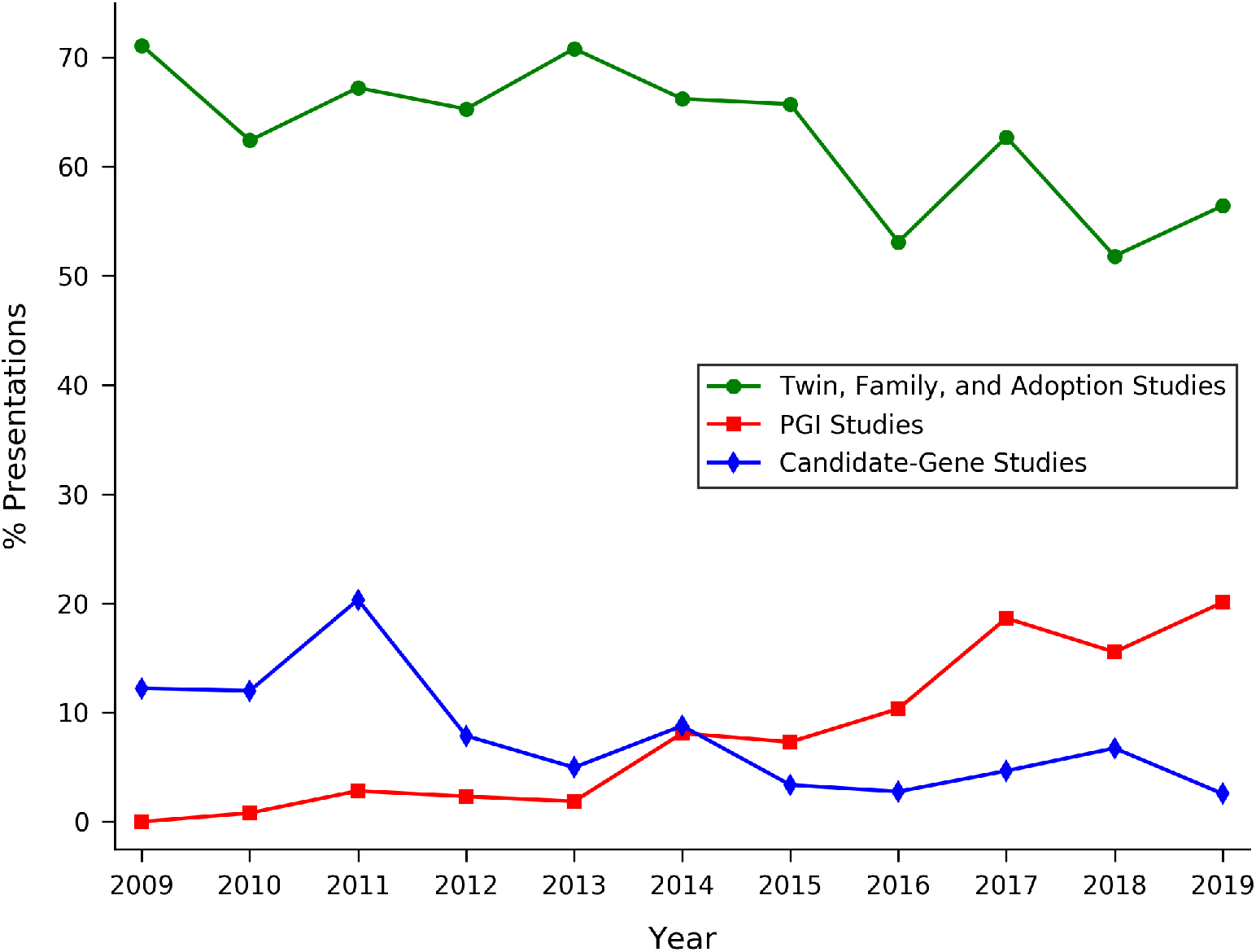
Type of study in presentations at Behavior Genetics Association Annual Meetings. Notes: For a description of the data underlying this figure, see Methods. Out of 1,993 presentations in total (over the 2009-2019 period), the percentages that are in exactly 0, 1, 2, or 3 categories are 26.76%, 67.56%, 5.5%, and 0.2%, respectively.

In this paper, we hope to promote productive behaviour-genetic research using PGIs in three ways. First and most centrally, we make a broad array of PGIs available via a Polygenic Index Repository, covering a number of datasets that may be useful to social scientists. By constructing the PGIs ourselves and making them available as variables downloadable from the data providers, our resource eliminates a number of roadblocks for researchers who would like to use PGIs in their research, as we detail below. The Repository contains PGIs for 47 phenotypes. To maximize prediction accuracy of the PGIs, we meta-analysed summary statistics from multiple sources, including several novel large-scale GWASs conducted in UK Biobank and the personal genomics company 23andMe. 23andMe shared summary statistics from 37 separate association analyses, 9 of which have not been reported previously. Therefore, almost all PGIs in our initial release perform at least as well as currently available PGIs in terms of prediction accuracy. We will update the Repository regularly with additional PGIs and datasets.

Second, we present a theoretical framework for interpreting associations with a PGI. Using this framework, we show that a PGI can be understood as an unbiased but noisy measure of what we call the “additive SNP factor,” which is the best linear predictor of the phenotype from the measured genetic variants. Because the PGI is a noisy measure, regressions that use the PGI as an explanatory variable suffer from errors-in-variables bias. Since different papers use different versions of a PGI, the magnitude of this bias varies. We hope that the theoretical framework helps establish a common language for discussions about the interpretation of PGIs and their effect sizes.

Third, we propose an approach that improves the interpretability and comparability of research results based on PGIs: to use in place of ordinary least squares (OLS) regression, we derive an estimator that corrects for the errors-in-variables bias. (We are aware of four papers to date that have implemented a measurement-error correction along the lines we propose here ^24–27^. Our approach is most similar to that of ref. ^26^, who develops a nearly identical framework using a psychometrics modeling approach but focuses on the univariate case.) The estimator produces coefficients in units of the standardized additive SNP factor, which has a more meaningful interpretation than units of some particular PGI. We illustrate by applying the estimator to multivariate and gene-by-environment regressions from a recently published paper ^20^. We make a Python command-line tool publicly available for implementing the estimator.

## Results

### The Polygenic Index Repository

The Polygenic Index Repository is a resource that addresses several practical obstacles that researchers interested in using PGIs must often confront. These include:

1. Constructing PGIs from individual genotype data can be a time-consuming process, even for researchers trained to work with large datasets.
2. Since the prediction accuracy of a PGI is increasing in the sample size of the underlying GWAS, it is generally desirable to generate PGI weights from GWAS summary statistics based on the largest available samples. However, privacy and IRB restrictions often create administrative hurdles that limit access to summary statistics and force researchers to trade off the benefit of summary statistics from a larger sample against the costs of overcoming the hurdles. In practice, researchers often end up constructing PGIs using only publicly available summary statistics.
3. Publicly available GWAS summary statistics are sometimes based on a discovery sample that includes the target cohort (or close relatives of cohort members) in which the researcher wishes to produce the PGI. Such sample overlap causes overfitting, which can lead to highly misleading results ^9^. (Sometimes, when GWAS consortia provide summary statistics upon request from a GWAS that is restricted so as to exclude the cohort, this barrier is surmounted at low cost.)
4. Because different researchers construct PGIs from GWAS summary statistics using different methodologies, it is hard to compare and interpret results from different studies.

We overcome #1 by constructing the PGIs ourselves and releasing them to the data providers, who in turn will make them available to researchers. This simultaneously addresses #2 because we use all the data available to us that may not be easily available to other researchers or to the data providers, including genome-wide summary statistics from 23andMe. Using these genome-wide summary statistics from 23andMe is what primarily distinguishes our Repository from existing efforts by data providers to construct PGIs and make them available, such as the effort by the Health and Retirement Study (https://hrs.isr.umich.edu/data-products/genetic-data/products#pgs). It also distinguishes our Repository from efforts to make publicly available PGI weights directly available for download ^28^. To deal with #3, for each phenotype and each dataset, we construct a PGI from GWAS summary statistics that excludes that dataset. We overcome #4 by using a uniform methodology across the phenotypes.

Figure 2 depicts the algorithm that determined which PGIs we constructed. In a preliminary step, we obtained GWAS summary statistics for a comprehensive list of 53 candidate phenotypes (see Supplementary Tables 1 and 2, meta-analyzed the summary statistics for each candidate phenotype, and calculated the expected *R*^2^ from an out-of-sample regression of each candidate phenotype on a PGI derived from its GWAS summary statistics (see Methods for details). If the expected explanatory power exceeded *R*^2^ = 0.01, then we used the meta-analysis output to construct a PGI for the phenotype. We call these the “single-trait PGIs.” For each candidate phenotype, we also identified a list of supplementary phenotypes: any other phenotype whose pairwise genetic correlation with the candidate exceeds 0.6 in absolute value. For each candidate with at least one supplementary phenotype, we then calculated the out-of-sample expected *R*^2^ of a PGI derived from a joint analysis of the candidate and supplementary phenotype summary statistics. If the expected *R*^2^ exceeded 0.01, then we used the joint-analysis output to construct a “multi-trait PGI” for the phenotype. When both single-trait and multi-trait PGIs are available, the multi-trait PGI generally has greater predictive power, but the single-trait PGI may be better suited for some applications (see Supplementary Methods).

**Figure 2:**
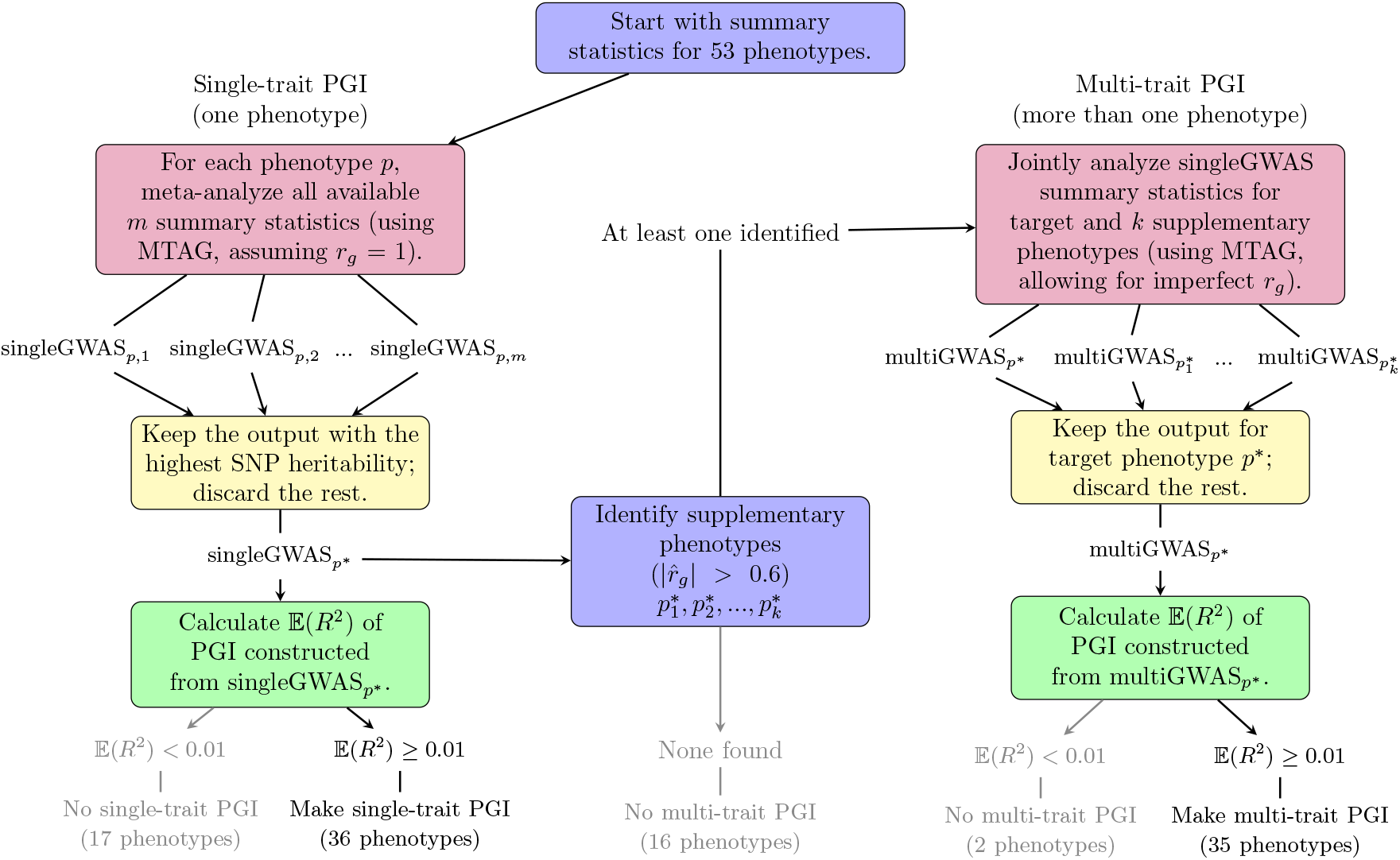
Algorithm determining which single-trait and multi-trait PGIs were generated for the Repository. Notes: See Table 1 for the 36 single-trait PGIs and 35 multi-trait PGIs included in the Repository.

**Table 1.** Repository phenotypes and GWAS sample sizes.

| Phenotype |  | GWAS Sample Size |  |  |  |  | PGIs Released |  | Suppl. Phenotypes |
| --- | --- | --- | --- | --- | --- | --- | --- | --- | --- |
|  |  | Total | 23andMe | UKB | Other | Public N | Single | Multi |  |
| Anthropometric |  |  |  |  |  |  |  |  |  |
| 1 | Body Mass Index (BMI) | 760,630 | - | 438,476 | 322,154 | 795,640 | X |  |  |
| 2 | Height | 698,334 | - | 445,054 | 253,280 | 709,706 | X |  |  |
| Cognition and Education |  |  |  |  |  |  |  |  |  |
| 3 | Childhood Reading | 172,503 | 172,503 | - | - | - | X |  |  |
| 4 | Cognitive Performance | 260,354 | - | 225,056 | 35,298 | 269,867 | X | X | 5, 6, 7 |
| 5 | Educational Attainment | 1,047,538 | 365,536 | - | 682,002 | 766,345 | X | X | 4, 6, 8, 33, 45 |
| 6 | Highest Math | 430,439 | 430,439 | - | - | - | X | X | 4, 5, 7, 8, 33 |
| 7 | Self-Rated Math Ability | 564,692 | 564,692 | - | - | - | X | X | 4, 6 |
| Fertility and Sexual Development |  |  |  |  |  |  |  |  |  |
| 8 | Age First Birth | 407,884 | 9,370 | 156733* | 241,781 | 241,781 | X | X | 5, 6, 11, 12, 19, 22 |
| 9 | Age First Menses (Women) | 329,345 | 76,831 | - | 252,514 | 252,514 | X | X | 10 |
| 10 | Age Voice Deepened (Men) | 55,871 | 55,871 | - | - | - |  | X | 9 |
| 11 | Number Ever Born (Men) | 260,991 | - | 168,056* | 92,935 | 165,492 |  | X | 8, 12 |
| 12 | Number Ever Born (Women) | 399,803 | - | 188,208* | 211,595 | 211,595 | X | X | 8, 11 |
| Health and Health Behaviors |  |  |  |  |  |  |  |  |  |
| 13 | Alcohol Misuse | 151,067 | 19,407 | 131,660 | - | - | X | X | 24 |
| 14 | Allergy - Cat | 46,646 | 46,646 | - | - | - |  | X | 15, 16, 17, 18, 26 |
| 15 | Allergy - Dust | 46,646 | 46,646 | - | - | - |  | X | 14, 16, 17, 18, 26 |
| 16 | Allergy - Pollen | 46,646 | 46,646 | - | - | - |  | X | 14, 15, 17, 19, 26 |
| 17 | Asthma | 445,965 | - | 445,965 | - | 361,141 | X | X | 14, 15, 16, 18, 26 |
| 18 | Asthma/Eczema/Rhinitis | 685,716 | 135,538 | 307,609* | 242,569 | 242,569 | X | X | 14, 15, 16, 17, 26 |
| 19 | Attention Deficit<br>Hyperactivity Disorder<br>(ADHD) | 117,754 | 62,380 | - | 55,374 | 55,374 | X | X | 8, 22 |
| 20 | Cannabis Use | 202,180 | 22,771 | 144,112 | 35,297 | 117,911 | X |  |  |
| 21 | Cigarettes per Day | 340,140 | 76,186 | - | 263,954 | 263,954 | X |  |  |
| 22 | Chronic Obstructive<br>Pulmonary Disease (COPD) | 445,965 | - | 445,965 | - | 91,787 |  | X | 8, 19, 30 |
| 23 | Depressive Symptoms | 942,579 | 307,354 | 404,984 | 230,241 | 500,199 | X | X | 30, 40, 43, 47 |
| 24 | Drinks per Week | 941,287 | 403,938 | - | 537,349 | 537,349 | X | X | 13 |
| 25 | Ever Smoker | 1,255,948 | 623,146 | - | 632,802 | 632,802 | X |  |  |
| 26 | Hayfever | 445,963 |  | 445,963 | - | 360,527 | X | X | 14, 15, 16, 17, 18,<br>Eczema† |
| 27 | Migraine | 693,993 | 283,985 | 410,008 | - | 361,194 | X |  |  |
| 28 | Nearsightedness | 367,906 | 191,843 | 176,063 | - | 360,677 | X |  |  |
| 29 | Physical Activity | 357,039 | 265,934 | - | 91,105 | 91,105 | X |  |  |
| 30 | Self-Rated Health | 1,203,099 | 758,713 | 444,386 | - | 359,681 | X | X | 22, 23, 37 |
|  |  |  |  | - |  |  |  |  |  |
|  | Personality and Well-Being |  |  | - |  |  |  |  |  |
| 31 | Adventurousness | 557,923 | 557,923 | - | - | - | X | X | 46 |
| 32 | Cognitive Empathy | 46,861 | 46,861 | - | - | - |  | X | Agreeableness† |
| 33 | Delay Discounting | 23,217 | 23,217 | - | - | - |  | X | 5, 6 |
| 34 | Extraversion | 122,255 | 59,225 | - | 63,030 | 63,030 | X | X | 35 |
| 35 | Left Out of Social Activity | 507,804 | 507,804 | - | - | - | X | X | 34, 38, 40, 47 |
| 36 | Life Satisfaction: Family | 168,313 | - | 168,313 | - | 118,818 | X | X | 38, 39, 47 |
| 37 | Life Satisfaction: Finance | 169,051 | - | 169,051 | - | 119,394 |  | X | 30, 40, 47 |
| 38 | Life Satisfaction: Friends | 168,001 | - | 168,001 | - | 118,649 | X | X | 35, 36, 39, 47 |
| 39 | Life Satisfaction: Work | 115,038 | - | 115,038 | - | 82,190 |  | X | 36, 38, 47 |
| 40 | Loneliness | 439,525 | - | 439,525 | - | 355,583 |  | X | 23, 35, 37, 43, 47 |
| 41 | Morning Person | 493,043 | 91,967 | 401,076 | - | 449,734 | X |  |  |
| 42 | Narcissism | 452,535 | 452,535 | - | - | - | X |  |  |
| 43 | Neuroticism | 484,560 | 59,206 | 361,688 | 63,666 | 380,060 | X | X | 23, 40, 47 |
| 44 | Openness | 76,551 | 59,176 | - | 17,375 | 17,375 | X |  |  |
| 45 | Religious Attendance | 444,842 | - | 444,842 | - | 360,063 | X | X | 5 |
| 46 | Risk | 1,427,867 | 969,309 | - | 458,558 | 466,571 | X | X | 31 |
| 47 | Subjective Well-Being | 1,022,510 | 728,752 | 169,219 | 124,539 | 204,978 | X | X | 23, 35, 36, 37, 38, 39,<br>40, 43 |

Notes: \*For Age First Birth, Number Ever Born (Men), Number Ever Born (Women) and Asthma/Eczema/Rhinitis, the publicly available summary statistics include the first release of UKB. Therefore, there is sample overlap between our UKB GWAS and publicly available summary statistics. For these phenotypes, in the UKB column, we report the UKB sample size excluding samples from the publicly available GWAS. †For Eczema and Agreeableness, both the single- and multi-trait PGIs had an expected predictive power less than 0.01, so they were used only as supplementary phenotypes for other phenotypes. Therefore, they are not included in the table and are not represented by a number. The GWAS sample for Eczema consists of only UKB, with $N = 440,177$ . The GWAS sample for Agreeableness consists only of 23andMe, with $N = 59,176$ .

For each of the 47 phenotypes for which we constructed a single-trait and/or multi-trait PGI, Table 1 lists the total sample size included in the GWAS summary statistics (Total *N*), followed by the sample-size contributions from three separate sources. For comparison, we also report the sample size of the largest GWAS whose summary statistics are in the public domain (Public *N*). With three exceptions, Total *N* exceeds Public *N*. Two exceptions are height and BMI, where our UKB sample inclusion filters lead to a slightly smaller sample size than the Public *N*. The remaining exception is cognitive performance, where the sample size of our GWAS is smaller due to overlap between the discovery sample in the largest GWAS with publicly available summary statistics and some of our Repository cohorts. For the remaining phenotypes, the gains in sample size relative to the public *N* are often substantial, and driven by our inclusion of summary statistics from large-scale GWASs conducted in 23andMe, UKB, or both. Table 1 also shows the 36 and 35 phenotypes for which we created single-trait and multi-trait PGIs, respectively. We created PGIs for these phenotypes in 11 Repository cohorts that shared their individual-level genetic data with us (regardless of whether the phenotype itself is measured in the cohort). Table 2 lists the datasets and some of their basic characteristics. Each data provider will make these PGIs available to researchers through their own data access procedures (see Supplementary Note).

**Table 2.** Datasets participating in the Repository.

| Dataset | *N* | Country | Population- or Family-based |
| --- | --- | --- | --- |
| Dunedin Multidisciplinary Health and Development Study | 887 | New Zealand | Population |
| English Longitudinal Study of Ageing (ELSA) | 7,310 | UK | Population |
| Environmental Risk (E-Risk) Longitudinal Twin Study | 2,316 | UK | Family |
| Estonian Genome Center, University of Tartu (EGCUT) | 51,719 | Estonia | Population |
| Health and Retirement Study (HRS) | 12,090 | USA | Population |
| Minnesota Center for Twin and Family Research (MCTFR) | 7,654 | USA | Family |
| National Longitudinal Study of Adolescent to Adult Health (Add Health) | 5,689 | USA | Family |
| Swedish Twin Registry (STR) | 38,072 | Sweden | Family |
| Texas Twin Project | 556 | USA | Family |
| UK Biobank (UKB) | 445,985 | UK | Population |
| Wisconsin Longitudinal Study (WLS) | 8,949 | USA | Family |

Notes: The “*N*” column gives the number of participants in each dataset for whom the PGIs in Table 1 are supplied in the initial release of the Repository (i.e., those who passed the subject-level exclusion filters described in Methods). “Population- or Family-based” refers to how individuals were recruited to the dataset.

The UK Biobank is among the 11 cohorts included in the Polygenic Index Repository. Because of its large sample size (see Table 2), the UK Biobank contributes substantially to the available sample for the GWAS for many phenotypes. We therefore did not want to exclude the entire UK Biobank from the GWASs used to create the PGIs. Instead, we split the UK Biobank sample into three equal-sized partitions. We ran three 1/3-sample GWASs for each phenotype. To create the PGI for each partition, we included results from the other two partitions in the meta-analysis. Consequently, researchers can conduct analyses of a PGI in any one of the partitions and obtain unbiased results. However, we caution researchers against conducting analyses in two or three of the partitions and meta-analyzing across partitions; because the other partitions are used to create the PGI, the results obtained across different partitions (although individually unbiased) will be correlated. Meta-analysis standard errors will therefore be anticonservative, and this bias can be substantial (see Methods). Therefore, to maximize the usefulness of our PGIs for research involving related individuals or brain-scan data, we assigned to the same partition all pairs of individuals that are related up to second degree (and some pairs of third degree), as well as all individuals with brain-scan data.

For validating the predictive power of the PGIs, we used five cohorts for which we had access to individual-level genetic and phenotypic data: the Health and Retirement Study, a representative sample of Americans over the age of 50; the Wisconsin Longitudinal Study, a sample of individuals who graduated from high school in Wisconsin in 1957; the Dunedin Multidisciplinary Health and Development Study, a sample of residents of Dunedin, New Zealand, born in 1972-1973; the Environmental Risk (E-Risk) (our third partition). The top panel of Figure 3 shows the observed *R*^2^ and 95% confidence intervals for Longitudinal Twin Study, a birth cohort of twins born in England and Wales in 1994-1995; and the UKB the single-trait PGIs in one or more validation cohorts, depending on which had a measure of the phenotype. Height, BMI, and educational attainment are shown separately because the *y*-axis scale is different. The bottom panel of Figure 3 shows the difference between the *R*^2^ of the single-trait Repository PGI and that of a PGI we constructed using the largest non-overlapping GWAS whose summary statistics are in the public domain. The Repository PGIs are almost always at least as predictive as the PGIs based on publicly available GWAS results. For the corresponding results for the multi-trait PGIs, which generally have higher *R*^2^’s than the single-trait PGIs, see Supplementary Figure 1.

**Figure 3:**
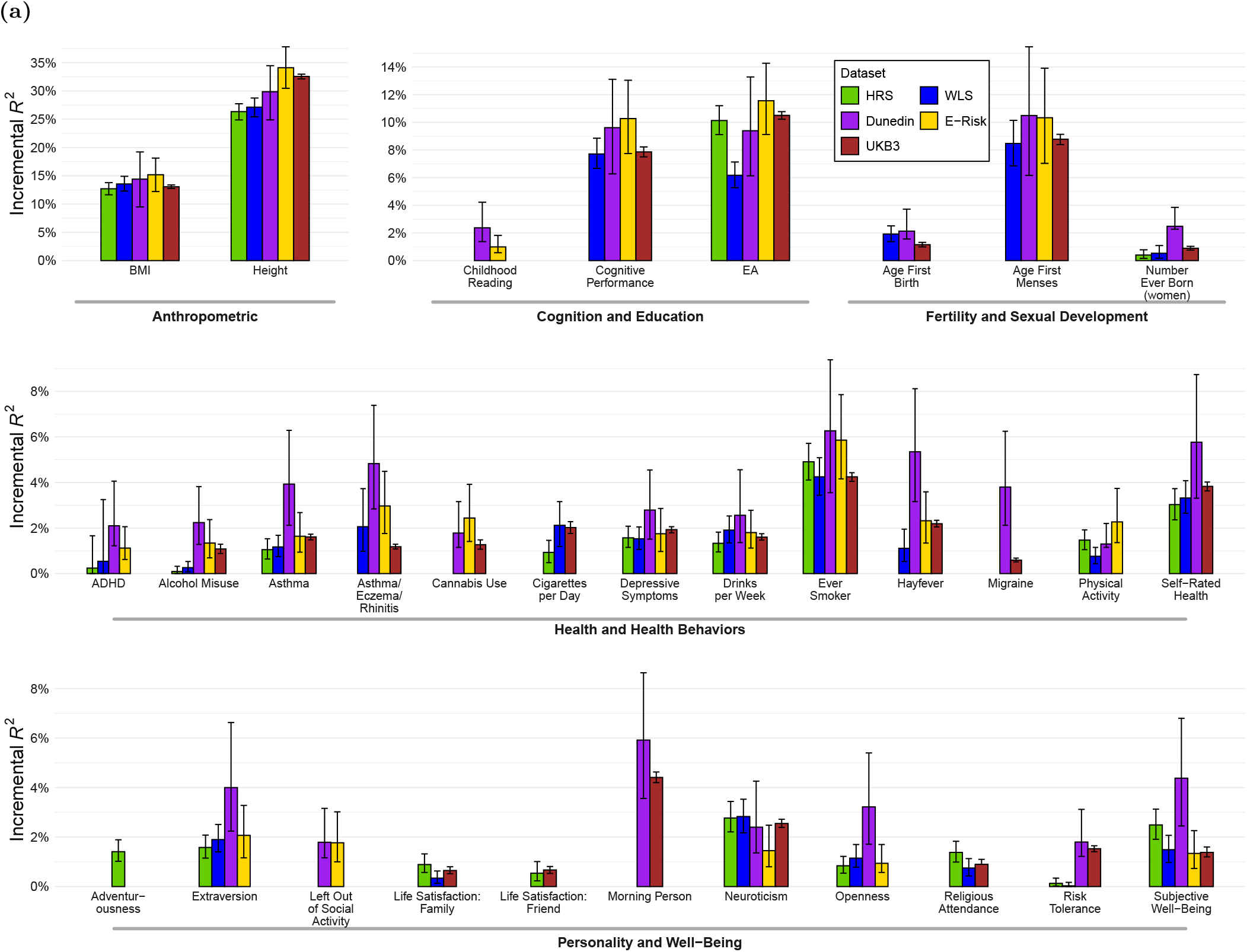
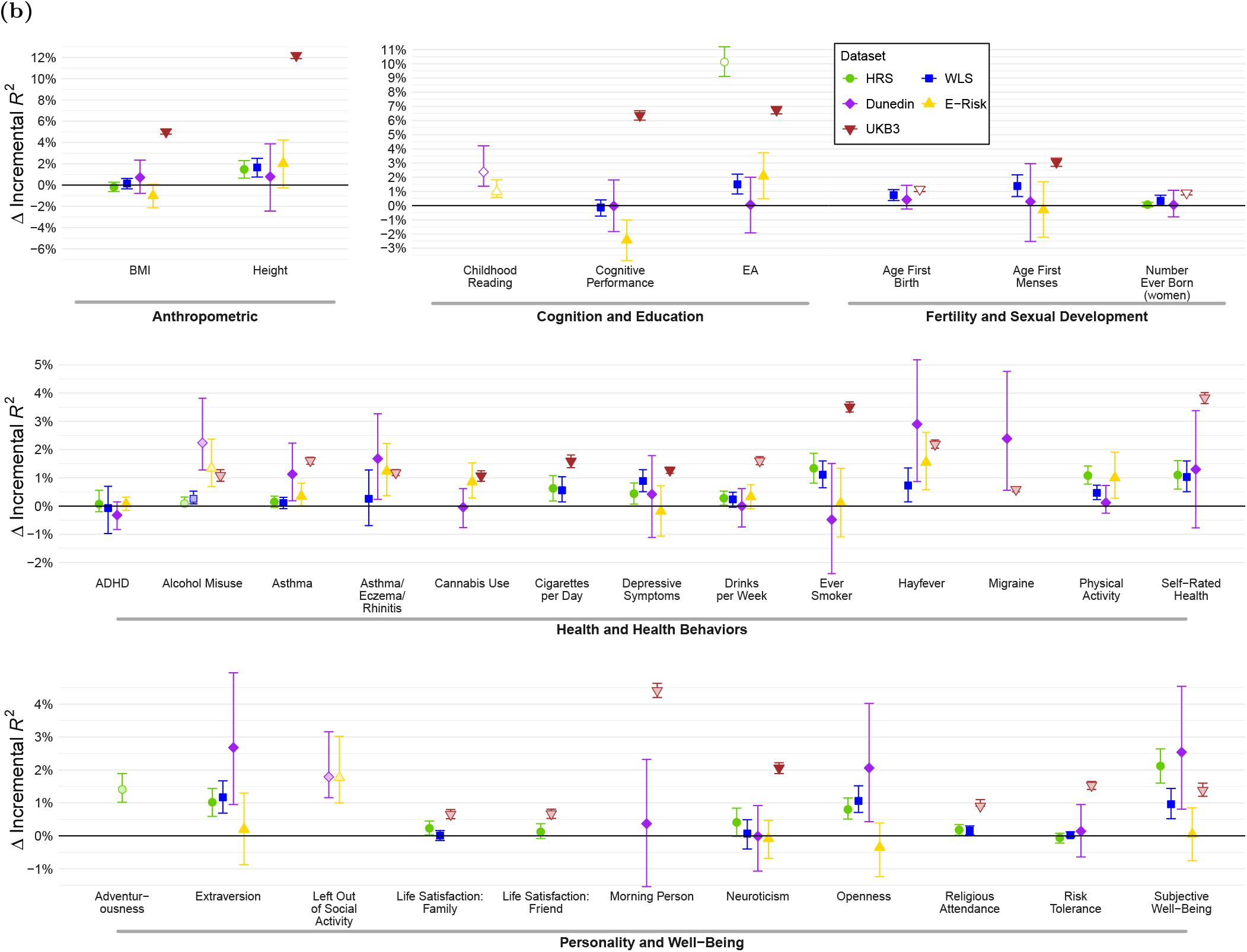
Predictive power of Repository single-trait PGIs. Notes: Error bars show 95% confidence intervals from bootstrapping with 1,000 repetitions. Panel (A): Incremental *R*^2^ from adding Repository’s single-trait PGI to a regression of the phenotype on 10 principal components of the genetic relatedness matrix for HRS, WLS, Dunedin and ERisk, and on 20 principal components and 106 genotyping batch dummies for UKB. Prior to the regression, phenotypes are residualized on a second-degree polynomial for age or birth year, sex, and their interactions (see Supplementary Tables 5 and 12). For the sample sizes of the GWAS that the PGIs are based on, see Supplementary Table 478. Panel (B): Difference in incremental *R*^2^ between Repository single-trait PGI and PGI constructed from publicly available summary statistics using our Repository pipeline. (Note that the latter do not include PGI directly available from cohortdatasets, such as the ones accessible from the HRS website.) If no publicly available summary statistics are available for a phenotype, then the difference in incremental *R*^2^ is equal to the incremental *R*^2^ of the single-trait PGI and is represented by an open circle. “Cigarettes per Day” in Dunedin was omitted from the Figure because the confidence interval (−5.99% to 0.94%) around the point estimate (−2.38%) required extending the y-axis substantially, making the figure hard to read. For the GWAS sample sizes of the PGIs based on publicly available summary statistics, see Supplementary Table 13.

We have written a User Guide (reproduced in the Supplementary Methods) that will be distributed by participating cohorts along with the Repository PGIs. It discusses interpretational issues, including those relevant for whether researchers should use the single-trait or multi-trait PGIs when both are available.

### Theoretical Framework for Polygenic Indexes

To help interpret PGIs, we lay out a theoretical framework. Denote individual *i*’s phenotype value by 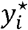 Denote individual *i*’s allele count at genetic variant *j* by 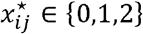. Without loss of generality, we use a mean-centred transformation of the phenotype and allele counts, such that 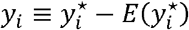 and 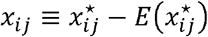 for each SNP *j*. We denote the vector of mean-centered allele counts at *j* genetic variants by ***x***_*i*_ = (*x*_*i*1_, *x*_*i*2_, … , *x*_*ij*_)′ As a benchmark, consider the standardized best linear predictor of the phenotype based on the allele counts:

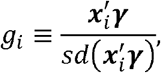

where

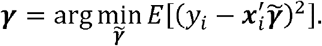

That is, the optimal weight vector **γ** is the vector of coefficients from the population regression of *y*_*i*_ on ***x***_*i*_ This population regression may also include control variables; we omit them here to avoid cluttering notation, but in the Supplementary Methods we extend the framework to include them and explain why they do not affect the results in this paper. In the User Guide (also in the Supplementary Methods), we explain how control variables do matter for the interpretation of a PGI.

When the set of genetic variants in ***x***_*i*_ is all variants in the genome, *g*_*i*_ is referred to as the “standardized additive genetic factor.” The variance in the phenotype explained by *g*_*i*_ is called the “(narrow-sense) heritability,” often the object of interest in twin, family, and adoption studies that draw inferences without access to molecular genetic data.

In studies with molecular genetic data—our focus here—the set of genetic variants in ***x***_*i*_ is restricted to those measured or imputed from the single-nucleotide polymorphisms (SNPs) assayed by standard genotyping platforms (and which pass quality-control filters). In that case, the variance in the phenotype explained by *g*_*i*_ is called the “SNP heritability”^29^, which we denote 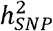. We will refer *g*_*i*_ as the standardized “additive SNP factor.”

Since the population regression cannot be run, the vector ***γ*** is unknown, so *g*_*i*_ cannot be constructed empirically. What can be constructed empirically is a “polygenic index (PGI),”, 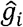 which is a standardized, weighted sum of allele counts using some other weight vector 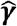 calculated from GWAS summary statistics:

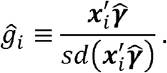

In general, 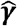 will not be equal to ***γ*** because 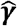 is calculated from GWAS summary statistics that are estimated in a finite sample. The key observation for our framework is that when 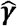 is calculated using standard methods (that include all the SNPs ***x***_*i*_), such as LDpred ^30^ and PRS-CS ^31^, the resulting PGI can be expressed as

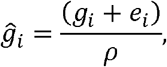

where *e*_*i*_ is mean-zero estimation error that is uncorrelated with *g*_*i*_, and 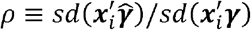 is a scaling factor that standardizes 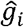. In words, the PGI is a standardized, noisy measure of the additive SNP factor, where the noise is classical measurement error.

One way to characterize the amount of measurement error is the value *ρ*. In Methods, we show that

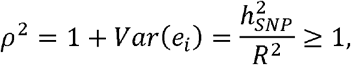

where 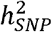 is the SNP heritability (the predictive power of *g*_*i*_) and *R*^2^ is the fraction of variance explained in a regression of the phenotype *y*_*i*_ on the PGI 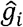 (the predictive power of 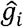). The ratio 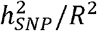 is greater than or equal to one because the weights that define *g*_*i*_ maximize the variance explained in *y*_*i*_ and therefore any other weights—including those used to construct the PGI—explain most 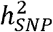 of the variation. Furthermore, the amount of measurement error *ρ* would achieve its minimum value of one only if the PGI weights were based on GWAS summary statistics from an infinite sample. Across studies, *ρ*^2^ varies. For example, *R*^2^ depends on the sample size of the GWAS underlying the PGI value of one only if the PGI weights were based on GWAS summary statistics from an infinite sample. weights and the method of constructing PGI weights (e.g., LDpred vs. PRS-CS). However, *ρ*^2^ can usually be estimated using estimates of 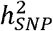 and *R*^2^ from the sample at hand or other samples that are sufficiently similar.

### Measurement-Error-Corrected Estimator for PGI Regressions

Typical research with a PGI involves running a regression with the PGI as an explanatory variable and reporting results in units of standard deviations of the PGI. This approach, however, has two shortcomings. First, it is often unclear how to interpret these units, which depend on the amount of measurement error. Second and relatedly, the effect sizes are not comparable across PGIs that differ in their amount of measurement error.

We argue that such a regression should be interpreted as aiming to approximate a regression with the standardized additive SNP factor as the explanatory factor. The PGI serves as an empirically feasible proxy for the standardized additive SNP factor. An analysis of the standardized additive SNP factor has a clearer interpretation than an analysis of the PGI and puts results in comparable units, regardless of which specific PGI was used in the analysis. Here we extend known results from errors-in-variables models to derive a consistent estimator for the coefficients from a regression with the standardized additive SNP factor as an explanatory variable.

The “theoretical regression” is what we call a regression with the (unobserved) standardized additive SNP factor as an explanatory variable. Consider an OLS regression of a phenotype *ϕ*_*i*_ on the standardized additive SNP factor *g*_*i*_ a vector of covariates ***z***_*i*_ and a vector ***w***_*i*_ of interactions between *g*_*i*_ and a subset of the regressors in ***z***_*i*_ (possibly all of them):

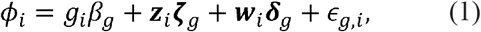

where the *g* subscripts indicate that these are parameters from the theoretical regression. (Note that the phenotype *ϕ*_*i*_ need not be the same phenotype *y*_*i*_ for which the standardized additive SNP factor is the best linear predictor. For example, some papers have studied the relationship between the PGI for educational attainment and test scores at younger ages ^14^. Note also that the covariates in ***z***_*i*_ may be measured with error; equation (1) represents whatever regression is run by a researcher except that *g*_*i*_ is measured without error.) The “feasible regression” is what we call the regression using the PGI 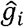 in place of *g*_*i*_:

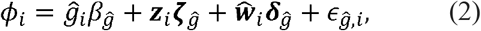

where 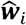 is the vector of interactions with 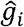 in place of *g*_*i*_ we denote the vectors of coefficients from the theoretical and feasible regressions by *α*_*g*_ ≡ (***β***_*g*_, ***ζ***_*g*_, ***δ*_*g*_**)′ and 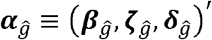 respectively.

In what follows, we sketch the derivation of an estimator for ***α***_*g*_ (for details, see the Supplementary Methods). The derivation assumes that the error in the PGI, *e*_*i*_, is uncorrelated with ***z***_*i*_ and ***w***_*i*_. In the Supplementary Methods, we show that this condition holds exactly if the PGI weights 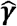 are unbiased estimates of ***γ***. We also show that if the PGI weights 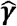 are estimated using LDpred-inf—as is true for the Repository PGIs—then the bias in our estimator due to plausible violations of this condition will typically be negligible.

Extending the standard formula for errors-in-variables bias ^32^ in a multivariate regression to this setting, and under the assumption that *e*_*i*_ is uncorrelated with ***z***_*i*_ and ***w***_*i*_, the feasible-regression coefficients can be shown to be biased:

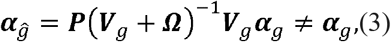

where 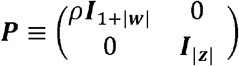**I**|*x*| is the identity matrix with the dimensionality of ***x***, ***V***_*g*_ is the variance-covariance matrix of (*g*_*i*_,***w***_*i*_,***z***_*i*_)′, and *Ω* is the component of the variance-covariance matrix of 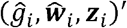 that is due to error (see Supplementary Methods). In the special case of a univariate regression, in which the only covariate is a constant term, equation (3) implies that the regression slope coefficient 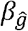 converges to 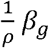. This is a familiar form of attenuation bias, in which the degree of attenuation toward zero is greater the larger the amount of measurement error. In the multivariate case, however, the amount of attenuation bias for 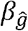 will also depend on the covariance matrix of *g*_*i*_ with ***z***_*i*_. Moreover, the other coefficients, 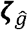 and 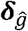 be biased as well, not necessarily toward zero. For example, a covariate whose coefficient in equation (1) is zero can have a coefficient in equation (2) that is non-zero, leading to an incorrect rejection of the null hypothesis (Abel (2017), unpublished manuscript).

The idea underlying our “corrected” estimator follows immediately from equation (3) by inverting the bias term:

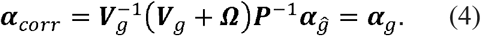

This expression is called a regression-disattenuation estimator. It cannot be implemented directly, however, because ***V***_*g*_ involves the variance and covariances of the unobserved standardized additive SNP factor *g*_*i*_. However, the variance and covariances involving *g*_*i*_ differ from analogous terms involving 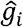 only due to measurement error, and the amount of measurement error is given by ρ. Therefore, the variance and covariances involving *g*_*i*_ can be inferred from estimable quantities. In the Supplementary Methods, we derive an expression for ***α***_*corr*_ in terms of *ρ* and population parameters that can be estimated consistently using the observed data. That expression is stated in Methods. We implement that version of the estimator. In the Supplementary Methods, we also derive standard errors for the regression coefficients, under the assumption that *ρ* is known.

If the PGI is uncorrelated with the covariates, then the estimator will inflate the naïve OLS estimate 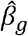 by the factor *ρ*. If, in addition, the covariates are uncorrelated with each other, then the estimator will also inflate 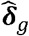 by the factor *ρ*. Correlation between the PGI and the covariates and correlation among the covariates will lead to deviations from this “rule of thumb” adjustment.

In the univariate case where ρ is estimated within the same dataset as the PGI analysis is conducted, we show that while uncertainty in 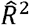 causes downward bias in the standard error, uncertainty in 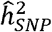 causes upward bias, and the net effect is likely to be standard errors that are slightly conservative. We conjecture that the standard errors will also typically be conservative in multivariate settings. If the *ρ* estimate is from a different dataset, then ignoring the uncertainty in *ρ* will unambiguously cause the standard errors to be anticonservative.

We provide a Python command-line tool that implements the measurement-error correction based on a user-specified value of *ρ*.^1^ The package can also estimate *ρ* by calculating estimates of 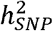 (using the GREML method ^29,33^ or, for larger datasets, BOLT-REML^34^) and *R*^2^. When possible, we recommend users estimate *ρ* within the dataset they use to analyse the PGI. If the dataset is too small to reliably estimate *ρ* or lacks a measure of the phenotype corresponding to the PGI, an estimate of *ρ* from another dataset can be used under the assumption of perfect genetic correlation of the phenotype across datasets. In the Polygenic Index Repository, we provide pre-specified estimates of *ρ* for three participating datasets for which we have access to the phenotypic data corresponding to the PGI: HRS, WLS, and the third partition of UKB (see Supplementary Table 4). For many of the cohorts, the standard error on the 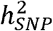 estimate is large, so we recommend a value of *ρ* based on existing 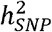 and *R*^2^ estimates from a larger sample.

Although our estimator is derived for an OLS estimation framework, it will be approximately correct for logistic regression ^35^ and survival models ^36^ as long as the coefficient on the standardized additive SNP factor, *β*_*g*_, is not too large. For example, applying a measurement-error correction that would be correct for OLS will be a very accurate approximation for the coefficient in a survival model when the hazard ratio associated with a one-standard deviation difference in the variable measured without error is 1.11 ^36^. However, the correction is roughly 20% too small when the hazard ratio is 1.65 ^36^.

### Illustrative Application

To illustrate our proposed measurement-error correction, we apply it to several analyses reported in a recent paper relating educational attainment (and labour market outcomes) to a PGI for educational attainment ^20^. The paper uses data from the HRS, one of our validation cohorts. As a preliminary analysis, the paper reports some straightforward tests of the relationship between educational attainment (EA) and the EA PGI. In Panel A of Table 3, we reproduce their univariate regression of EA on the PGI and their multivariate regression that additionally includes controls for mother’s and father’s EA. In the univariate regression, shown in column (1), a 1-standard-deviation increase in the PGI is associated with 0.823 additional years of schooling. This association is reduced to 0.619 years in column (2), once the controls are included.

**Table 3.**
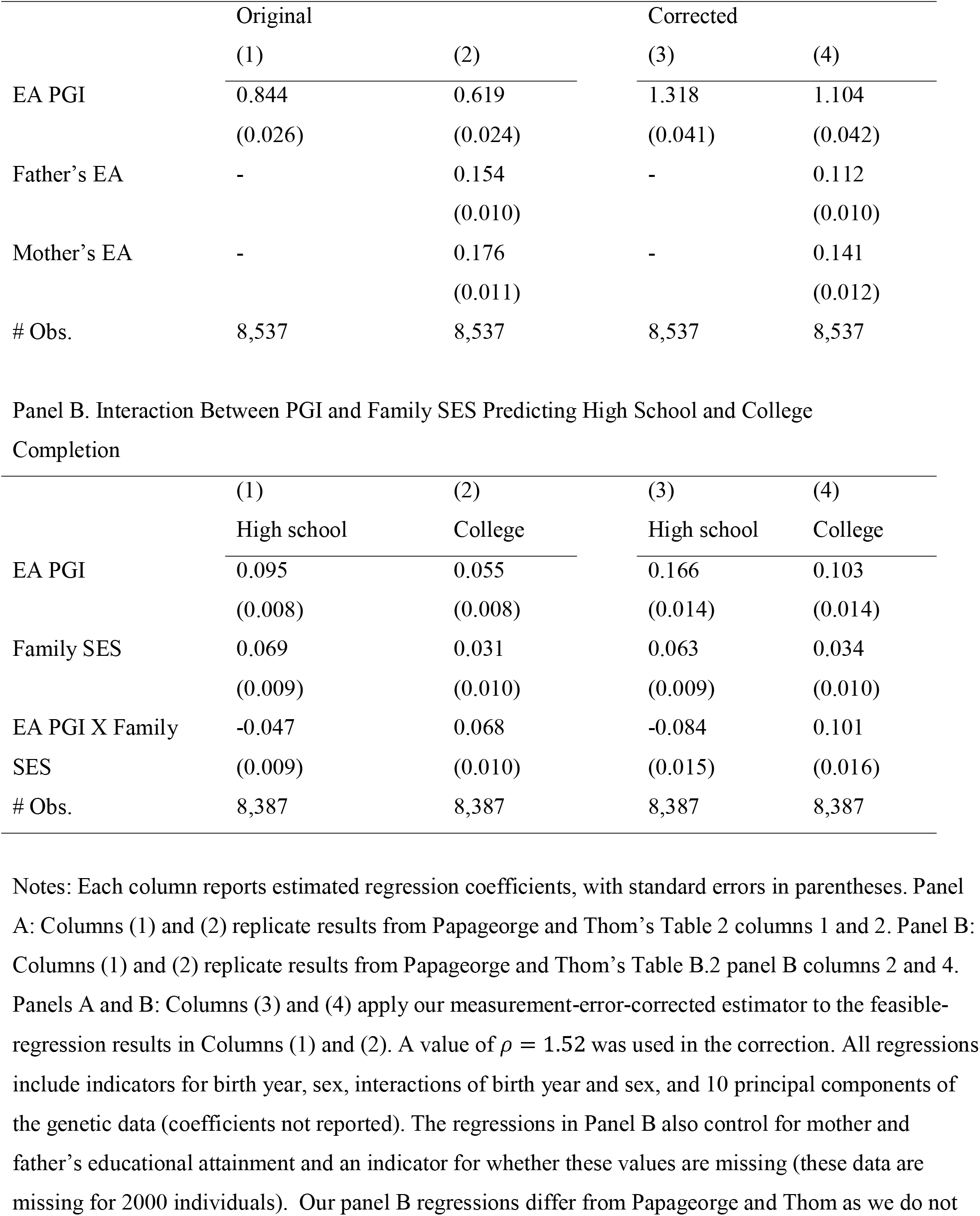
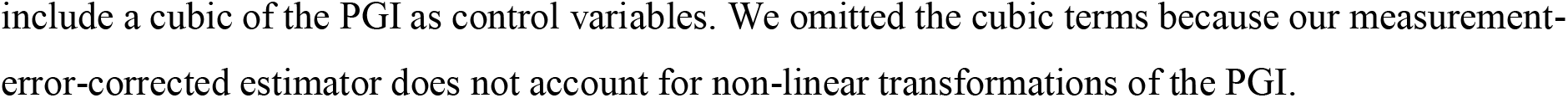
Application of measurement-error correction. Panel A. Association Between EA and the PGI, Without and With Controls for Parental EA

The measurement-error-corrected univariate regression is shown in column (3) of Panel A. We estimate that a 1-standard-deviation increase in the additive SNP factor is associated with 1.288 additional years of schooling. Relative to the PGI coefficient in column (1), this coefficient is larger by a factor of 1.288 / 0.823 = 1.57. In the regression with controls for parental education, shown in column (4), we estimate a corrected coefficient of 1.123 additional years. Relative to column (2), this is an increase by a factor of 1.123 / 0.610 = 1.84. Since for EA in the HRS, 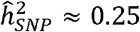 and 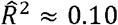, according to the rule of thumb mentioned above, both coefficients should be expected to have increased by a factor of 1.58 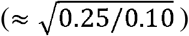. The increase is larger than that from column (2) to (4) due to the positive correlations between the PGI, the controls, and the dependent variable.

The results in Panel A illustrate a general implication of the measurement-error correction for mediation analyses: the correction deflates estimates of how much covariates mediate the effect of the PGI. There have been several mediation analyses in which researchers study how much the coefficient on a PGI is reduced when control variables—which are usually positively correlated with both the PGI and the dependent variable—are added to the regression ^37–39^. Going from column (1) to (2), the drop in the coefficient on the PGI would lead a researcher to conclude that parental education mediates (0.823 – 0.610) / 0.823 = 26% of the effect of the PGI. Going from column (3) to (4) shows the corrected estimate of mediation is only (1.288 – 1.123) / 1.288 = 13%. The drop is larger for the uncorrected regressions because in those regressions, the control variables are proxying for part of the additive SNP factor that is not well captured by the PGI. Therefore, studies that do not correct for measurement error will tend to overestimate the extent to which the control variables mediate the effect of the PGI.

The results in Panel B illustrate a fairly general implication of the measurement-error correction for PGI-by-environment interaction analyses: in contrast to how it affects mediation estimates, the correction tends to increase the magnitude of PGI-by-environment interaction estimates. A main result of Papageorge and Thom is about two such interactions: a higher PGI is associated with a weaker relationship between childhood SES and high school completion but a stronger relationship between childhood SES and college completion ^20^. Columns (1) and (2) reproduce two specifications that show this result: a regression of high school completion on the PGI, self-reported childhood SES, their interaction, and controls; and the analogous regression for college completion. The key finding is that the interaction term is negative in column (1) but positive in column (2). As shown in columns (3) and (4), once the additive SNP factor is considered instead of the PGI, the interaction coefficients for both the high school and college regressions move farther away from zero, strengthening the main result of the paper. In general, PGI-by-environment interaction studies that do not correct for measurement error will tend to underestimate the magnitude of the interaction because the interaction term will tend to be attenuated by the measurement error. Note, however, that this conclusion may not hold if other regressors are correlated with the interaction term.

## Discussion

We described the initial release of the Polygenic Index Repository, which contains PGIs for 47 phenotypes. A major goal of this effort is to disseminate PGIs with greater predictive power than the PGIs typically used. To maximize prediction accuracy of the PGIs, we meta-analysed data from multiple sources, including 23andMe and the UK Biobank.

We also derived a measurement-error-corrected estimator that can be used instead of OLS regressions where the independent variables include a PGI or a PGI and its interactions. While some lack of comparability of results across studies is inevitable (e.g., due to differences across samples in SNP heritabilities), one goal of both the Repository and the proposed estimator is to increase comparability. For example, when constructing the PGIs, we applied to each cohort uniform sets of inclusion criteria for individuals and markers in the genotype data. The estimator contributes to improving comparability by putting regression coefficients in units of the additive SNP factor, regardless of the predictive power of the particular PGI available to the researchers.

Because genetic associations are easily misinterpreted, researchers who use PGIs should be especially careful to understand and convey the appropriate interpretation of their findings. For example, it is important to keep in mind that PGI associations may be mediated by environmental factors, and these factors may be modifiable. To facilitate understanding of these and other interpretational issues, we have written a User Guide that cohorts will distribute to users of the Repository PGIs (see Supplementary Methods).

As more GWAS summary statistics become available in the years ahead, and better methods for constructing PGIs are developed, we plan to update the Repository regularly with more predictive PGIs that leverage these advances. For example, future releases will incorporate PGIs of novel phenotypes for which it is not currently feasible to construct PGIs with meaningful predictive power. We emphasize, however, that although PGIs have attained levels of predictive power that can be useful to researchers, the limited heritability of behavioural phenotypes such as those in the Repository implies that the PGIs will never be able to predict any individual’s phenotype with much precision. Additionally, since GWAS summary statistics have only been available in large samples of individuals from European ancestries, currently available PGIs have limited portability to individuals of non-European ancestries ^40^. In future releases of the Repository, once sufficient data becomes available to create PGIs that have non-negligible predictive power for other ancestry groups, we will update the Repository to contain such PGIs.

## Methods

The polygenic indexes (PGIs) shared through the Repository are based on summary statistics from three types of sources: novel GWASs conducted in UK Biobank (UKB), GWASs conducted in samples of volunteer research participants from 23andMe, and other published genome-wide association studies (GWAS). In Section I below, we begin by describing how the summary statistics used in our main analyses were generated, quality-controlled and meta-analysed to generate a set of files used as inputs into construction of the single-trait and multi-trait PGIs. In Section II, we define and justify the *R*^2^ criterion we used to determine which PGIs to include in the first release of the Repository. We then describe quality-control filters applied to the individual-level genotype data supplied by each Repository cohort. We conclude by describing the methods used to construct the cohort PGIs. In Section III we state our measurement-error-corrected estimator and its standard error in terms of estimable quantities. Section IV describes our estimation of *ρ* in the HRS, WLS and UKB. Section V describes the data underlying Figure 1.

### I. Summary Statistics

#### UKB GWAS

Supplementary Table 5 lists all UKB phenotypes for which we ran novel GWASs. Before running the GWASs, we filtered out poor-quality genotypes: (i) samples identified as putatively carrying sex-chromosome configurations that are neither XX nor XY, (ii) samples identified as outliers in heterozygosity and missingness rates, (iii) samples whose sex inferred from sex chromosomes does not match self-reported gender, and (iv) samples with missing sex, birth year, genotyping batch, or PC information. We also restricted the sample to individuals we will refer to as of “European ancestries,” defined as the first genetic PC provided by UKB being greater than 0 and individual self-reporting to be of “British”, “Irish”, or “Any other white background.”

In order to make PGIs for the UK Biobank (UKB) without having to exclude the entire UKB from the discovery GWAS, we split the UK Biobank sample into three equal-sized partitions and, for each partition, used the summary statistics from the other two partitions when generating its PGI. The first partition (UKB1) is composed of UKB participants with brain-scan data (as indicated by data field 12188), all pairs of UKB participants related up to second degree, and the pairs of relatives of third-degree relatedness with greatest relatedness. Pairs of individuals of third-degree relatedness were ordered based on the maximum relatedness coefficient they have with another participant and assigned to the first partition in decreasing relatedness order until the partition was full. Remaining individuals with third-degree relatives were assigned to the second partition. Finally, individuals with no third degree or closer relatives were randomly assigned to the second (UKB2) or third (UKB3) partition.

For all phenotypes in Supplementary Table 5, we ran three separate GWASs, one for each partition. Briefly, each GWAS in UKB was conducted using mixed-linear models implemented by the software BOLT-LMM ^41^. The dependent variable in each analysis is a phenotype that has been residualized on sex, a third-degree polynomial in birth year (defined as (*birthyear* - 1900)/10), their interactions, 106 genotyping batch dummies, and the first 40 of the PCs released by the UK Biobank. Details on how each phenotype is coded are provided in Supplementary Table 5. For the variance-component estimation in BOLT-LMM (but not the association analyses), we restricted the set of markers to the set of 622,788 hard-called SNP genotypes that remained after filtering for 1% minor allele frequency and 60% imputation accuracy and pruning with an *r*^2^ threshold of 0.3. Our subsequent association analyses were performed on imputed SNP dosages provided by UKB.

#### Using the UK Biobank split-sample PGI

Splitting the UKB into thirds as described above increases the predictive power of the PGI within each third (relative to omitting the UKB from the GWAS sample). Researchers may desire to conduct analyses that simultaneously include individuals from different partitions of the data or to meta-analyse results across different partitions. Such analyses will produce estimates that are unbiased, but the standard errors will be incorrectly calibrated. To see why, consider a linear model

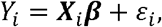

where ***X***_*i*_ is a vector of covariates that includes a PGI. Imagine that the data (***Y***, ***X***) include individuals from different partitions of the data. As a result of the sample-splitting procedure above, Cov(***X***_*i*_,*ɛ*_*i*_) = 0, which implies that the OLS estimator for ***β***, will be unbiased. However, because some of the individuals in the data were used to generate the PGI for other individuals in the data, Cov(***X***_*i*_, *ɛ*_*j*_) ≠ 0 whenever individuals *i* and *j* are in different partitions. As a result,

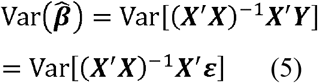

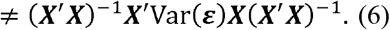

The expression (6) is the standard general formula for the sampling variance of OLS estimates. It is not equal to (5) due to the correlation between (***X***′***X***)^−1^***X***′ and ***ɛ***. If we knew the correlation between these two vectors, we could calculate correct standard errors in this setting, but the correlation structure is complex, and we are unaware of any current method that produces correct standard errors. For this reason, we recommend that researchers only do analyses on sets of individuals within a partition. If researchers choose to do analyses with individuals across different partitions, they should include the strong caveat that their standard errors may be poorly calibrated.

#### 23andMe GWAS

Our analyses use summary statistics from GWASs conducted by 23andMe in samples of European-ancestry volunteer research participants for 37 different phenotypes. Supplementary Table 6 provides an overview of these summary statistics. 28 out of the 37 are from previously published studies^5,42–55^. For these, we cite the original study in the column labelled “Citation”. The remaining 9 are based on novel, and previously unreported, GWASs. Two of the novel GWASs are for phenotypes (Subjective Well-Being and Risk) for which GWASs had been previously published by 23andMe but with a smaller sample. The remaining summary statistics have not been previously published by 23andMe. Supplementary Table 6 describes the details of the association model used for each phenotype. For details on 23andMe’s genotyping and imputation, see Supplementary Tables 17 and 18 in Lee et al.^5^

#### Quality control of summary statistics

We applied a uniform set of quality-control filters to each original file with summary statistics (both those from novel GWASs and previously published GWASs). We closely followed the quality-control pipeline detailed in section 1.5.1 of Okbay et al. ^37^ and implemented in the software EasyQC ^56^. Our QC protocol departed from Okbay et al. in the following steps:

– We used data from the Haplotype Reference Consortium reference panel (r1.1) ^57^ to check for strand misalignment, allele mismatch, chromosome and base pair position concordance, and allele frequency discrepancies (instead of using data from the 1000 Genomes Phase 1 ^58^). (Mapping file and allele frequency data were downloaded from the EasyQC website, from the following urls, respectively: https://homepages.uni-regensburg.de/~wit59712/easyqc/HRC/HRC.r1-1.GRCh37.wgs.mac5.sites.tab.rsid_map.gz, https://homepages.uni-regensburg.de/~wit59712/easyqc/HRC/HRC.r1-1.GRCh37.wgs.mac5.sites.tab.cptid.maf001.gz.)
– For simplicity and uniformity, we applied a more conservative imputation accuracy filter of 0.7 to all input files irrespective of the software that was used for imputation.
– We applied a uniform minor allele frequency filter of 0.01 to all input files. Stricter filters varying by sample size were not necessary because the studies that we analysed were much larger than some of those in Okbay et al.
– We filtered out standard-error outliers. To do so, we first estimated the standard deviation 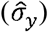 of the phenotype in each input file by regressing the reported standard errors on the following approximation to the standard error of a coefficient estimated by OLS when the phenotype is standardized:

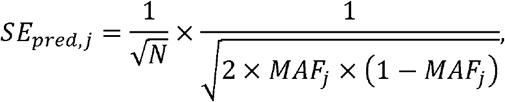

where *MAF*_*j*_ is the minor allele frequency of SNP *j* and *N* is the GWAS sample size. We filtered out markers with 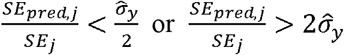. This filter allowed us to identify and remove markers for which the reported GWAS sample size deviated considerably from the sample size implied by the marker’s standard error. This filter was particularly relevant for publicly available summary statistics, where marker-specific sample sizes were typically not reported. (Having an accurate number for the sample size is important for LDpred ^30^.)

Before each filtered file was cleared for subsequent meta-analyses, we also prepared and visually inspected a number of diagnostic plots, as described in Okbay et al. Our final analyses are limited to files whose diagnostic plots did not suggest any anomalies. Finally, we examined the genetic correlation between input files (estimated using the LDSC software package ^59^) for each phenotype to make sure phenotype coding was in the same direction across 23andMe, UKB, and published studies. Supplementary Table 7 summarizes the number of SNPs dropped in each filtering step in the files that passed all diagnostic checks.

#### Single-Trait Input GWAS

In this section, we describe the construction of single-trait input GWASs used in several of our downstream analyses, including as inputs for the single-trait and multi-trait PGIs. The single-trait input GWAS for a phenotype is obtained by meta-analysing summary statistics from up to three sources of information: analyses in UKB, analyses in 23andMe, and summary statistics from a previously published study of the phenotype^5,42,49,50,52,60–73^. The input GWAS for a phenotype is the same across most cohorts. However, when there is overlap between a Repository cohort and cohorts that contributed to summary statistics from previously published studies, or in order to construct a PGI for a UKB partition that is based on summary statistics including the rest of the UKB sample, we restrict the meta-analyses to summary statistics based on non-overlapping data. Details on the construction of single-trait input GWAS are in Supplementary Table 8.

To illustrate the general procedure, consider the single-trait input GWAS for neuroticism in ELSA and EGCUT. Supplementary Table 8 shows that the largest meta-analysis of neuroticism (NEURO1) yielded a final sample of *N* = 484,560individuals by combining data from UKB (*N* = 361,688), 23andMe (*N* = 59,206) and a previously published study (*N* = 63,666). Since the column does not indicate any overlap with ELSA, the single-trait input GWAS for neuroticism in ELSA is the set of summary statistics from this meta-analysis. EGCUT, however, is listed in Supplementary Table 8 as overlapping with the NEURO1 meta-analysis. The reason is that EGCUT contributed to the summary statistics of the single-trait input is therefore generated by meta-analysing the summary statistics from UKB (*N* = previously published study (it is one of the cohorts in de Moor et al. ^66^). To eliminate overlap, EGCUT’s 361,688) and 23andMe (*N* = 59,206) only. This restricted meta-analysis is listed in the table as NEURO2. Similarly, the largest single-trait input GWAS for neuroticism includes the UKB, so all three UKB partitions are listed as overlapping with it. To eliminate overlap, the single-trait input for each UKB partition (which are labelled NEURO3, NEURO4, and NEURO5) is generated by meta-analysing 23andMe, de Moor et al., and the remaining two UKB partitions.

Each input GWAS is conducted by meta-analysing the relevant input files in MTAG ^74^. All analyses are conducted allowing for sample overlap and setting all genetic correlations equal to unity. However, we allow the SNP-heritability parameter to vary across input files. Even though MTAG produces a separate output file for each input file, the assumption of perfect genetic correlation ensures that the SNP coefficients in each output file are a constant multiple of each other (hence the PGIs generated by the file with the highest estimated SNP heritability as the input GWAS (this matters for the expected *R*^2^ output files are the same). In all analyses that follow, we adopt the convention of designating the output calculation but nothing else). The details of the heritability estimation are described below, in the subsection “Criterion for Inclusion in Repository” in Section II.

#### Multi-Trait Input GWAS

For several phenotypes in the first-wave release of the Repository, we provide multi-trait PGIs. Here, we describe the multi-trait input GWAS used to generate each of these.

In a first step, we used LDSC ^59^ to estimate genetic correlations between the phenotypes in with the largest Total *N*. This restriction leaves 53 single-trait input GWAS files, each of which is Supplementary Table 8. For phenotypes with multiple single-trait input GWAS files, we used the version associated with a distinct phenotype. Because there may be sample overlap between the meta-analysed summary statistics, we used GWAS-equivalent sample sizes as reported by MTAG when estimating genetic correlations. (This was the case for Age First Birth, Number Ever Born (men), Number Ever Born (women), and Asthma/Eczema/Rhinitis. For the first three phenotypes, we meta-analysed the publicly available summary statistics from Barban et al. ^73^, which included the first release of UKB, with UKB full release. Similarly, for Asthma/Eczema/Rhinitis, we meta-analysed publicly available summary statistics from Ferreira et al. ^49^, which included the first release of UKB, with UKB full release.) The set of pairwise genetic correlations is reported in Supplementary Table 9.

In a second step, we identified each Repository phenotype’s supplementary phenotypes. A phenotype is supplementary to a target phenotype (and vice versa) if the pairwise genetic correlation between the phenotypes exceeds 0.6 in absolute value. Under this definition, the estimates in Supplementary Table 9 identify each target phenotype’s supplementary phenotypes. These are listed in the column “Input files” of Supplementary Table 10 (set to “No Supplementary Phenotypes” if the phenotype has genetic correlation less than 0.6 with all other phenotypes). For 37 of the 53 Repository phenotypes, we identified at least one supplementary phenotype.

In a final step, for each of these 37 phenotypes, and for each Repository cohort, we ran a multivariate MTAG analysis on the target phenotype together with its supplementary phenotypes, using the version of the target phenotype and each supplementary phenotype for which the cohort is listed in the column “Repository Datasets Sumstats are Used For” in Supplementary Table 8. (In some cases, the same version of the target phenotype and each supplementary phenotype were used for more than one cohort; in those cases, we ran the MTAG analysis only once for that group of cohorts.)

Each MTAG analysis produces multiple output files—one for the target phenotype and one for each of the supplementary phenotypes—but we only retain the summary statistics for the target phenotype. In what follows, we refer to each such file as a multi-trait input GWAS.

### II. Constructing Repository PGIs

#### Criterion for Inclusion in Repository

The previous section described how we generated single-trait and multi-trait input GWASs from which it is straightforward to generate single-trait and multi-trait PGIs for a large number of phenotypes. We now describe how we determined, for each candidate phenotype, whether to include neither the single- nor multi-trait PGI, both PGIs, or one of the two in the initial release of the Repository. The structure of our algorithm is outlined in Figure 2. This section provides the details.

For both single- and multi-trait PGIs, we limited the initial set of PGIs released to those with an out-of-sample expected *R*^2^ above 1%. While the threshold itself is arbitrary, the decision to have a threshold was driven by two considerations: the value of a PGI for research is increasing in its predictive power, and we worried that a PGI with low predictive power could cause more harm than good if researchers are tempted to conduct underpowered studies.

We calculated the expected predictive power of each PGI (that might potentially be included in the Repository) using the following formula from Daetwyler et al. ^75^:

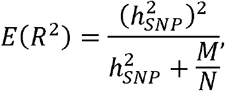

where 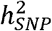 is the phenotype’s SNP heritability, *M* is the effective number of independent SNPs which we assume to be equal to 60,000 ^9^, and *N* is the GWAS sample size for the phenotype.

We first used the formula above to project the expected predictive power of each potential single-trait PGI. Our projections for the 53 potential PGIs and the underlying parameter values assumed are shown in Supplementary Table 1. We set 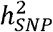 equal to the SNP heritability estimated by LDSC in the summary statistics from the single-trait input GWAS file with the largest sample size for a phenotype. We set *N* equal to the GWAS-equivalent sample size reported in the MTAG output. For the 37 phenotypes with at least one supplementary phenotype, we generated similar projections for the multi-trait PGIs, using the Multi-Trait Input GWAS files instead. The results of the 37 projections, and the underlying parameter values assumed, are shown in Supplementary Table 2.

We find that our criterion results in 47 phenotypes with at least one PGI in the Repository (see Figure 2). For 12 phenotypes, our procedure results in the release of a single-trait PGI but no multi-trait PGI; these are the phenotypes with no supplementary phenotypes. For 11 other phenotypes, our procedure results in the release of a multi-trait PGI but no single-trait PGI; these are typically phenotypes without large GWASs but for which we have multiple supplementary phenotypes with large GWASs. Finally, our procedure yields 24 phenotypes with both single- and multi-trait PGIs that satisfy our inclusion criterion and 6 phenotypes for which neither PGI qualifies.

#### Genotyping and Imputation in Repository Cohorts

Genotyping was performed using a range of commercially available arrays. Cohorts were encouraged to upload genotypes imputed against the 1000 Genomes Phase 3 ^76^ or HRC ^57^ imputation panels. Some cohorts provided only genotyped SNPs or data imputed against an older panel. In those cases, we performed the imputation against the HRC reference panel (version 1.1) using the Michigan Imputation Server ^77^. Supplementary Table 11 provides study-specific details on the genotyping arrays, pre-imputation quality control filters, imputation software used, and reference samples.

#### Genotype Data QC in Repository Cohorts

We restricted the set of markers to the SNPs present in the third phase of the international HapMap project (HapMap 3) ^78^ in order to reduce computational burden (relative to using all reported SNPs) while keeping a set of markers that covers most of the common variation in individuals with European ancestries.

#### Subject-level QC in Repository Cohorts

We restricted the samples to individuals with European ancestries. Exclusion criteria were based on the first four principal components of the genetic data. In order to obtain the principal components, for each cohort, we first converted the imputed genotype dosages for HapMap3 SNPs into hard calls. We then merged the data with all samples from the third phase of the 1000 Genomes Project, restricting to SNPs that had a call rate greater than 99% and minor allele frequency greater than 1% in the merged sample. We calculated the principal components (PCs) in the 1000 Genomes subsample and projected these onto the remaining individuals in the merged data. In order to select European-ancestry samples, we plotted the first four PCs against each other and visually identified the individuals that cluster together with the 1000 Genomes EUR sample.

#### Creation of PCs in Repository Cohorts

In the Repository cohorts, before constructing PCs, we removed markers with imputation accuracy less than 70% or minor allele frequency less than 1%, as well as markers in long-range LD blocks (chr5:44mb-51.5mb, chr6:25mb-33.5mb, chr8:8mb-12mb, chr11:45mb-57mb). Next, we restricted the sample to individuals with European ancestries, as described immediately above. We further pruned the markers to obtain a set of approximately independent markers, using a rolling window of 1000 base pairs (incremented in steps of 5) and an *R*^2^ threshold of 0.1. We used this set of markers to estimate a genetic relatedness matrix. We identified all pairs of individuals with a relatedness coefficient greater than 0.05 as calculated by Plink1.9 ^79^. We excluded one individual from each pair, calculated the first 20 PCs for the resulting sample of unrelated individuals using Plink 1.9, and projected the PCs onto the sample of unrelated individuals.

#### Constructing PGIs

All PGIs in the initial release of the Repository were constructed in Plink2 ^79^ using imputed genotype probabilities. Prior to constructing the PGIs, we adjusted the SNP weights for linkage disequilibrium (LD) using LDpred ^30^. We estimated the LD patterns using genotype data from the public release of the HRC Reference Panel (version 1.1) after applying the following quality-control filters. First, we limited the set of variants to HapMap3 SNPs and filtered out variants with genotyping call rate <0.98 and individuals with genotype missingness rate >0.02. Next, we calculated the genomic relatedness matrix and dropped one individual out of each pair with relatedness coefficient >0.025. We clustered the remaining individuals based on their identity-by-state distances using Plink1.9 and dropped an individual if the *Z*-score corresponding to their distance to their nearest neighbour is less than -5. In the remaining sample that we fed into LDpred for LD estimation, there were 1,214,408 SNPs and 14,028 individuals. At the coordination step of LDpred, we used the option “--max-freq-discrep” in order to exclude markers that used the “--z-from-se” option so that *Z* statistics were obtained from the GWAS coefficient estimates and have a frequency discrepancy greater than 0.1 between the summary statistics and genotype data. We also their standard errors rather than from *P* values (the default) because the latter led to issues in LDpred for markers with extremely small *P* values. For each PGI, we used the LD window recommended by Vilhjalmsson et al. ^30^, i.e., the number of markers common between the LD reference data, cohort genotype data and summary statistics left after the remaining LDpred quality control filters (MAF > 0.01, no allele mismatch, no ambiguous alleles), divided by 3,000. The fraction of causal markers was set to 1 for each phenotype to ensure consistency across phenotypes.

#### Prediction Analyses

We conducted a validation exercise for our new PGIs in the HRS, WLS, Dunedin, E-Risk, and UKB (third partition) cohorts. Supplementary Table 12 describes the phenotypes used as outcomes in these analyses for all cohorts except UKB. The UKB phenotypes are described in Supplementary Table 5. (The UKB phenotypes used in the prediction exercise differ slightly from the GWAS phenotypes described in Supplementary Table 5 in that they were not residualized on the PCs and genotyping batch dummies. Instead, we have controlled for these covariates in the regressions when calculating incremental *R*^2^ as described below.) As a general rule, if a single measurement in time was available, we residualized the phenotype on a second-degree polynomial in age, sex, and their interactions. If multiple measurements were available, we either did the same residualization in each wave and took the mean across waves or we took the maximum across waves and then residualized on birth year, sex, and their interactions.

Supplementary Table 3 shows the results from the prediction analyses. The incremental *R*^2^ was calculated as the difference in explained variance when adding the PGI to a regression of the residualized phenotype on the first 10 principal components of the genetic data. In the UKB prediction analyses, we included an additional 10 principal components and 106 genotyping batch dummies. We obtained 95% confidence intervals around the incremental *R*^2^’s by bootstrapping with 1000 repetitions. Supplementary Table 3 also shows the predictive power of “public PGIs”, which are PGIs constructed using our Repository pipeline based on the largest publicly available GWAS on the phenotype that does not have sample overlap with the prediction cohort ^4,5,62–70,73,37,80–85,42,49,50,52,55,60,61^ (we also use http://www.nealelab.is/uk-biobank/). The details of the input GWAS used for each validation cohort for the construction of the “public PGIs” are in Supplementary Table 13.

### III. Measurement-Error-Corrected Estimator

Equation (4) in the main text gives an expression for our measurement-error-corrected estimator, but it cannot be implemented directly because ***V***_*g*_ and Ω are based on unobserved variables. In the Supplementary Methods we derive an equivalent expression in terms of variables that can all be consistently estimated using sample analogues:

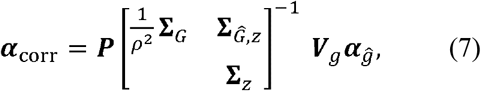

where

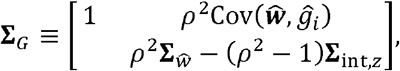

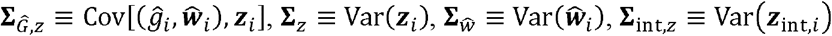, and ***z***_*int,i*_ the vector of the covariates that are interacted with *g*_*i*_ to form the vector ***w***_*i*_.

To obtain standard errors for ***α***_*corr*_, we calculate

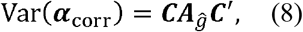

where 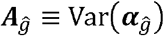 and

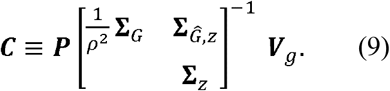

The standard errors are the square root of the diagonal of Var(***α***_*corr*_). Note that equations (7)–(9) are written in terms of population variance-covariance matrices, model coefficients, and the parameter *ρ* To implement this correction, we replace each of these terms with its sample counterpart.

### IV. Estimation of p in HRS, WLS and UKB

We estimated the value of *ρ* for all PGIs satisfying the criterion for inclusion in the Repository in three of our validation datasets: HRS, WLS and UKB (partition 3). Recall from the main text that *ρ* is defined as

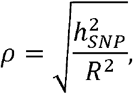

where 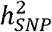 is the SNP heritability and *R*^2^ is the fraction of variance explained in a regression of the phenotype on the PGI.

In order to estimate 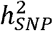 and *R*^2^, we first took the residualized phenotypes described in section “Prediction Analyses” and additionally residualized these on 20 PCs in HRS and WLS, and 40 PCs and batch effects in UKB3. We did the same for the PGIs. In HRS and WLS, we estimated 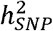 with genomic-relatedness-matrix restricted maximum likelihood (GREML) implemented in GCTA v1.93.0beta^29,33^ using HapMap3 SNPs with MAF > 1%. Prior to the 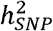 estimation, we dropped one individual from each pair with a relatedness greater than 0.025. We estimated *R*^2^ as the explained variance in a simple regression of the residualized phenotype on the residualized PGI. Standard errors for *R*^2^, 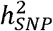, and *ρ* were estimated with a 100-block jackknife procedure.

In UKB3, because of the large sample size, we faced computational constraints. We therefore used the REML implementation in BOLT v2.3^34^ (with the `--remlNoRefine` option). Moreover, we estimated standard errors only for three phenotypes: friend satisfaction, educational attainment, and height. We chose these three phenotypes so as to have one each corresponding to a single-trait PGI with low (friend satisfaction), medium (educational attainment) and high predictive power (height).

Supplementary Table 4 lists the estimates of *ρ* for HRS, WLS and UKB3, along with the underlying 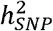 and *R*^2^ estimates and standard errors where available.

### V. Categorization of Behavior Genetics Association Annual Meeting Presentations

To obtain the data for Figure 1, we first created a dataset containing the titles, authors, and abstracts of all presentations at the 2009-2019 Behavior Genetics Association Annual Meetings. The information about the presentations is printed each year in issue six of the association journal Behavior Genetics. There were 2,034 presentations in this initial dataset. Included in the initial dataset were 36 symposia and 5 papers that were submitted as a part of symposia; all 41 of these are omitted from the final dataset. The final dataset contains a total of 1,993 presentations.

After some trial-and-error and visual inspection of several dozen abstracts, we arrived at the algorithm below for categorizing studies:

- We categorized a presentation as a “PGI study” if the title or the abstract contains at least one of the following keywords: ‘PGS’, ‘PRS’, ‘PGRS’, ‘polygenic score’, ‘polygenic risk score’, ‘genetic risk score’, ‘GRS’.
- We categorized a presentation as a “twin, family, or adoption study” if it satisfies at least one of the following conditions:

– The abstract contains ‘twin’ at least twice.
– The title contains the word ‘twin’.
– The title or abstract contain at least one of the following keywords: ‘twin registry’, ‘center for twin research’, ‘twin project’, ‘twin panel’, ‘twin study at the’, ‘twin study (LTS)’, ‘(RFAB) twin study’, ‘twin register’, ‘twin pairs’, ‘nonidentical twins’, ‘identical twins’, ‘pairs of twins’, ‘twin sample’, ‘ MZ’, ‘ DZ’, ‘monozygotic’, ‘dizygotic’, ‘pairs of twins’, ‘adopted’, ‘adoptee’, ‘adoptive’, ‘adoption design’, ‘biological parent’, ‘adoptive parent’, ‘adoption-sibling’, ‘genetically-unrelated’, ‘genetically-related’, ‘siblings reared together’, ‘siblings reared apart’, ‘mother and child’, ‘father and child’, ‘parent and child’, ‘intergenerational’, ‘transracial’, ‘biometric’, ‘path analy’, ‘Cholesky’, ‘children-of-twins’, ‘children of twins’, ‘common environment’, ‘unique environment’, ‘ACE’, ‘ACDE’.
- We categorized a presentation as a “candidate-gene study” if it satisfies at least one of the following conditions:

– The title contains ‘candidate gene’ or at least one of the following candidate gene keywords: ‘HTR2’, ‘MAOA’, ‘5-HTT’, ‘5HTT’, ‘DRD’, ‘SLC6’, ‘BDNF’, ‘COMT’, ‘TPH’, ‘MTHFR’, ‘APOE’, ‘DTNBP1’, ‘DBH’, ‘ABCB1’, ‘VNTR’, ‘CRHR’, ‘AKT’, ‘NRG’, ‘AVP’, ‘rs0’, ‘rs1’, ‘rs2’, ‘rs3’, ‘rs4’, ‘rs5’, ‘rs6’, ‘rs7’, ‘rs8’, ‘rs9’.
– The abstract contains at least one of the above candidate-gene keywords.
– The abstract contains ‘candidate’ at least twice and ‘candidate gene’ at least once.

However, a presentation was removed from the candidate-gene study category if the abstract contains GWAS keywords: ‘wide association analysis’, ‘wide association study’, ‘GWAS’.

To quantify how accurately the algorithmic classifications predict categorizations based on human evaluations, we asked two researchers with expertise in behaviour genetics to categorize 65 randomly sampled presentations. The raters worked independently, without any external assistance, and based their categorizations solely on information supplied about the title and abstract. Each rater assigned three yes/no labels—representing candidate-gene study; twin, family or adoption study; or PGI study—to each presentation. Raters sought to make labelling decisions consistent with the labels’ typical usage in the literature. We defined “agreement” on a presentation as an identical judgment about each of the three labels (i.e., if the raters disagreed about any of the three categories, they were considered as not agreeing). Even under this strict definition, we found an interrater agreement of 94%. The agreement between the algorithm’s and one rater’s categorizations was 86%, and that between the algorithm’s and the other rater’s categorizations was 83%.

## Supporting information

FAQ

## Data availability

For how to access the Repository PGIs and other data from each participating dataset, see Supplementary Note; upon publication, an up-to-date list of participating datasets and data access procedures will be maintained at https://www.thessgac.org/pgi-repository. For each phenotype that we analyse, we report GWAS and MTAG summary statistics and PGI (LDpred) weights for all SNPs from the largest discovery sample for that analysis, unless the sample includes 23andMe. SNP-level summary statistics from analyses based entirely or in part on 23andMe data can only be reported for up to 10,000 SNPs. Therefore, if the largest GWAS or MTAG analysis for a phenotype includes 23andMe, we report summary statistics for only the genome-wide significant SNPs from that analysis. In addition, we report summary statistics and PGI (LDpred) weights for all SNPs from the largest GWAS or MTAG analysis that does not include 23andMe. These summary statistics and PGI weights can be downloaded from http://www.thessgac.org/data upon publication. The data underlying Figure 1 will also be available at http://www.thessgac.org/data. Researchers at non-profit institutions can obtain access to the genome-wide summary statistics from 23andMe used in this paper by completing the 23andMe Publication Dataset Access Request Form, available at https://research.23andme.com/dataset-access/.

## Code availability

Upon publication, the software used for the measurement-error correction will be available at https://github.com/JonJala/pgi_correct. The code for constructing PGIs and principal components, for the illustrative application, and for analyzing the data displayed in Figure 1 will be available at http://www.thessgac.org/data.

## Acknowledgements

We thank C. Shulman for helpful comments. This research was carried out under the auspices of the Social Science Genetic Association Consortium (SSGAC). This research was conducted using the UK Biobank Resource under application number 11425. The study was supported by funding from the Ragnar Söderberg Foundation (E42/15, D.C.), the Swedish Research Council (421-2013-1061, M.J.; 2019-00244, S.O.), an ERC Consolidator Grant (647648 EdGe, P.K.), the Pershing Square Fund of the Foundations of Human Behavior (D.L.), Open Philanthropy (010623-00001, D.J.B.), Riksbankens Jubileumsfond P18-0782:1 (S.O.), Netherlands Organisation for Scientific Research VENI grant 016.Veni.198.058 (A.O.), and the NIA/NIH through grants R24-AG065184 (D.J.B.) and R01-AG042568 (D.J.B.) to the University of California Los Angeles; K99-AG062787-01 (P.T.) to Massachusetts General Hospital; 1R01-MH101244-02 (P.T.; PI: Benjamin M. Neale) and 5U01-MH109539-02 (P.T.; PI: B.M.N.) to the Broad Institute at Harvard and MIT; R56-AG058726 (Titus Galama) to the University of Southern California; the Government of Canada through Genome Canada and the Ontario Genomics Institute (OGI-152) (J.P.B.); the Social Sciences and Humanities Research Council of Canada (J.P.B.), the National Health and Medical Research Council through grant GNT113400 (P.M.V.); and the Australian Research Council. We thank the following consortia for sharing GWAS summary statistics: Reproductive Genetics (ReproGen) Consortium for age at first menses; Genetics of Personality Consortium (GPC) for neuroticism, extraversion, and openness; Psychiatric Genomics Consortium (PGC) for ADHD and depressive symptoms; Tobacco and Alcohol Genetics (TAG) Consortium for cigarettes per day and ever smoker; International Genomics of Alzheimer’s Project (IGAP) for Alzheimer’s disease, GWAS & Sequencing Consortium of Alcohol and Nicotine use (GSCAN) for cigarettes per day, ever smoker and drinks per week; Genetic Investigation of Anthropometric Traits (GIANT) Consortium for height and BMI; and Cognitive Genomics (COGENT) Consortium for cognitive performance. We thank the Neale Lab for making UK Biobank GWAS results available for asthma, cannabis use, COPD, hayfever, life satisfaction (family, finance, friend, work), loneliness, migraine, nearsightedness, number ever born (men, women), religious attendance, self-rated health and subjective well-being. We thank the research participants and employees of 23andMe for making this work possible. A full list of acknowledgements is provided in the Supplementary Note.

## Author contributions

D.J.B., D.C., A.O., and P.T. designed and oversaw the study. A.O. supervised all analyses and led the writing of the manuscript. J.B. was the lead analyst, responsible for the GWAS and MTAG analyses, quality control of GWAS summary statistics, and the PGI validation analyses. C.A.P.B. was responsible for quality control of genotype data and the construction of PGIs. G.G., N.W., H.J., and M.B. assisted with analyses. G.G. conducted the illustrative application and wrote the Python code. N.W. designed and implemented the algorithm used to generate Figure 1. R.K.L. ran a meta-analysis of general risk tolerance omitting validation cohorts. P.T. derived the measurement-error-correction estimator. R.A., A.Y., J.P.B., P.K., S.O., M.J., P.V., M.N.M., and D.L. contributed to study design. All authors contributed to and critically reviewed the manuscript. D.J.B., A.O., D.C. and P.T. made especially major contributions to the writing and editing. Cohort-level contributions are in the Supplementary Note.

