## Supplementary Figures for "Resource Profile and User Guide of the Polygenic Index Repository"

**Supplementary Figure 1. Predictive power of Repository multi-trait PGIs**
**(A)**

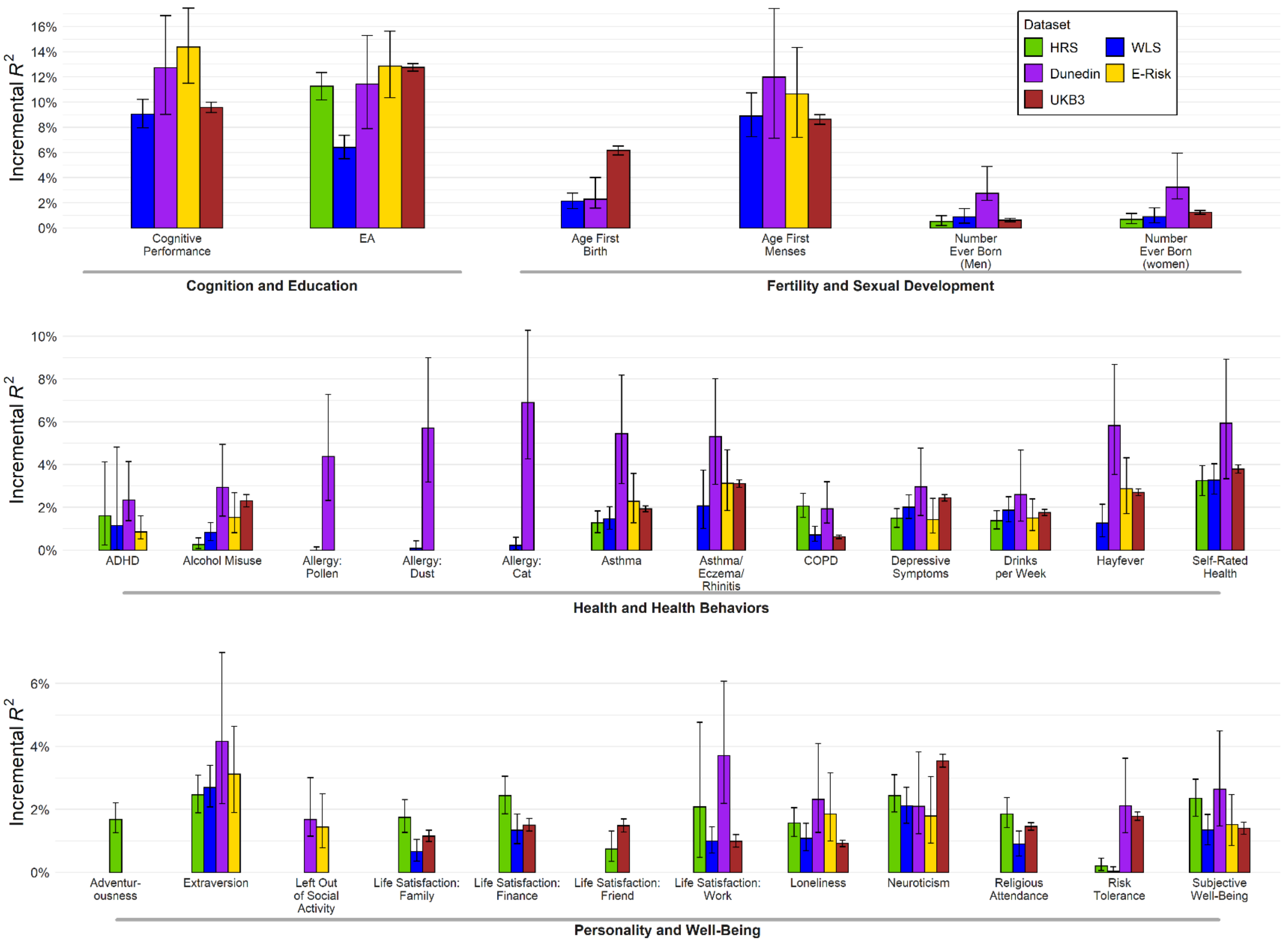

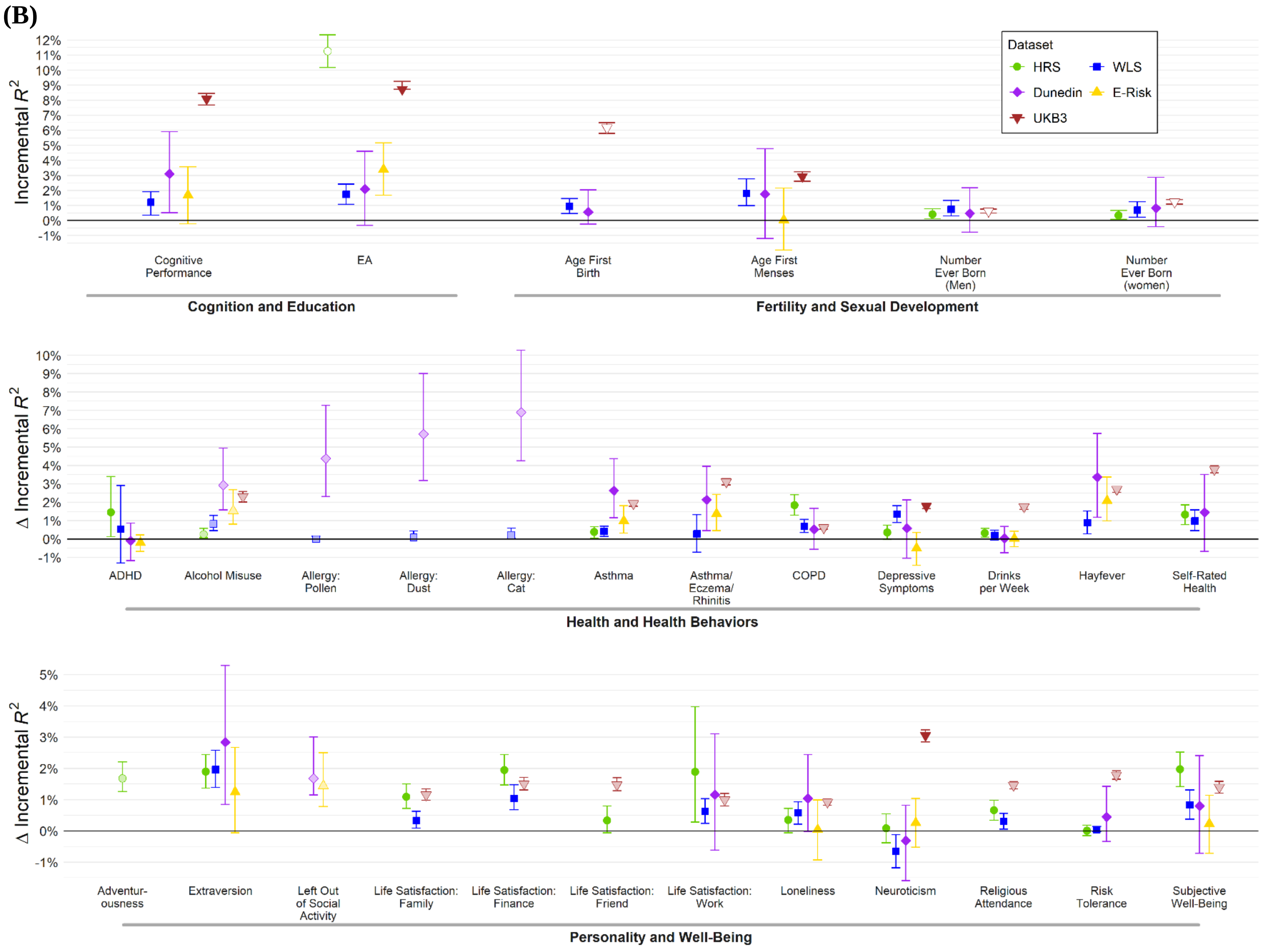

(B)
Incremental $R^2$
12%
11%
10%
9%
8%
7%
6%
5%
4%
3%
2%
1%
0%
-1%
Dataset
HRS
WLS
Dunedin
E-Risk
UKB3
Cognitive Performance
EA
Age First Birth
Age First Menses
Number Ever Born (Men)
Number Ever Born (women)
Cognition and Education
Fertility and Sexual Development
Δ Incremental $R^2$
ADHD
Alcohol Misuse
Allergy: Pollen
Allergy: Dust
Allergy: Cat
Asthma
Asthma/ Eczema/ Rhinitis
COPD
Depressive Symptoms
Drinks per Week
Hayfever
Self-Rated Health
Health and Health Behaviors
Δ Incremental $R^2$
Adventur-ousness
Extraversion
Left Out of Social Activity
Life Satisfaction: Family
Life Satisfaction: Finance
Life Satisfaction: Friend
Life Satisfaction: Work
Loneliness
Neuroticism
Religious Attendance
Risk Tolerance
Subjective Well-Being
Personality and Well-Being

**Notes**: Error bars show 95% confidence intervals from bootstrapping with 1,000 repetitions. Panel (A): Incremental $R^2$ from adding Repository's multi-trait PGI to a regression of the phenotype on 10 principal components of the genetic relatedness matrix for HRS, WLS, Dunedin, and E-Risk, and on 20 principal components and 106 genotyping batch dummies for UKB. Prior to the regression, phenotypes are residualized on a second-degree polynomial for age or birth year, sex, and their interactions (see Supplementary Tables 5 and 12). For the GWAS-equivalent sample sizes of the summary statistics that the PGIs are based on, see Supplementary Table 10. Panel (B): Difference in incremental $R^2$ between Repository multi-trait PGI and PGI constructed from publicly available summary statistics using our Repository pipeline. (Note that the latter do not include PGI directly available from datasets, such as the ones accessible from the HRS website.) If no publicly available summary statistics are available for a phenotype, then the difference in incremental $R^2$ is equal to the incremental $R^2$ of the single-trait PGI and is represented by an open circle. For the GWAS sample sizes of the PGIs based on publicly available summary statistics, see Supplementary Table 13.
