## Supplementary Methods for "Resource Profile and User Guide of the Polygenic Index Repository"

April 23, 2021

**5 Potential Bias in the Standard Errors** **14**

**6 Theoretical Framework with GWAS Controls** **15**

**7 Polygenic Index Repository User Guide** **15**

### 1 Setup

Denote individual $i$'s phenotype by $y_i^\star$. For some genetic variant $j$, denote the allele count for individual $i$ by $x_{ij}^\star$. Without loss of generality in this derivation, we use mean-centered transformation of the phenotype and allele counts, $y_i \equiv y_i^\star - \mathbb{E}(y_i^\star)$ and $x_{ij} \equiv x_{ij}^\star - \mathbb{E}(x_{ij}^\star)$. We denote the vector of mean-centered allele counts across $J$ SNPs for individual $i$ by $\mathbf{x}_i = (x_{i1}, x_{i2}, \ldots, x_{iJ})$.

As a benchmark, consider the standardized best linear predictor of $y_i$ given the $J$ SNPs, $\mathbf{x}_i$:

$$g_i \equiv \frac{\mathbf{x}_i\gamma}{\mathrm{sd}(\mathbf{x}_i\gamma)}, \tag{1}$$

where

$$\gamma \equiv \arg\min_{\tilde{\gamma}} \mathbb{E}\left[(y_i - \mathbf{x}_i\tilde{\gamma})^2\right]. \tag{2}$$

Thus, $g_i$ is the weighted sum of genotypes that maximizes the expected power for predicting $y_i$ using a linear combination of genotypes. If the set of $J$ genetic variants is the set of all genetic variants, then $g_i$ is referred to as the standardized additive genetic factor. The variance of $y_i$ explained by the standardized additive genetic factor is called the narrow-sense heritability. When the set of $J$ genetic variants is some set of genotyped SNPs, we refer to $g_i$ as the standardized additive SNP factor and the variance of $y_i$ is referred to as the SNP heritability.

We use $h^2_{SNP}$ to denote the SNP heritability, which is the variation in $y_i$ explained by the additive SNP factor. Because $g_i$ is standardized, the SNP heritability of $y_i$ is also the squared correlation of $y_i$ and $g_i$:

$$h^2_{SNP} \equiv \frac{[\mathrm{Cov}(y_i, g_i)]^2}{\mathrm{Var}(y_i)\,\mathrm{Var}(g_i)} = \frac{[\mathrm{Cov}(y_i, g_i)]^2}{\mathrm{Var}(y_i)}.$$

By basic properties of population regression, we can decompose $y_i$ into two uncorrelated components,

$$\begin{aligned} y_i &= \frac{\mathrm{Cov}(y_i, g_i)}{\mathrm{Var}(y_i)} g_i + \varepsilon_{y,i} \\ &= \frac{\mathrm{Cov}(y_i, g_i)}{\sqrt{\mathrm{Var}(y_i)}} \frac{1}{\sqrt{\mathrm{Var}(y_i)}} g_i + \varepsilon_{y,i} \\ &= \frac{h_{SNP}}{\sqrt{\mathrm{Var}(y_i)}} g_i + \varepsilon_{y,i}, \end{aligned}$$

where $\varepsilon_i$ is the component of the phenotype that is uncorrelated with $g_i$. We have ignored covariates in this definition of the additive SNP factor. For a model that includes covariates, we may define $y_i$ and each $x_{ij}$ as the phenotype and genotypes after having been residualized for the set of covariates.

A PGI for phenotype $y_i$ is also a weighted sum of genotypes,

$$\hat{g}_i \equiv \sum_j x_{ij}\hat{\gamma}_j. \tag{3}$$

The PGI will have maximum predictive power only when $\hat{\gamma}_j = \gamma_j$ for every SNP $j$. In practice, methods for constructing a PGI calculate the $\hat{\gamma}_j$'s using GWAS summary statistics together with some procedure to account for linkage disequilibrium (e.g., pruning and thresholding, LD-based shrinkage-based methods). The PGI is then usually standardized to have mean zero and variance one. In the theory below, we treat the PGI as if it has been standardized.

Projecting the PGI onto the space spanned by the standardized additive SNP factor, we can express the PGI as

$$\hat{g}_i = \frac{g_i + e_i}{\mathrm{sd}(g_i + e_i)}, \tag{4}$$

where $\mathrm{Cov}(g_i, e_i) = 0$. In Section 2.2 below, we assume that $e_i$ is uncorrelated with all other variables when the prediction sample is independent of the sample used to estimate $\hat{\gamma}$. (We highlight there where the assumption is used.) As shown in Section 4, this assumption will be a very good approximation when the PGI is constructed using LDpred-inf, as is the case for all the PGIs in the Repository.

The predictive power of the PGI, $R^2$, is therefore

$$\begin{aligned}
R^2 &\equiv \frac{\left[\operatorname{Cov}\left(y_i, \hat{g}_i\right)\right]^2}{\operatorname{Var}\left(y_i\right) \operatorname{Var}\left(\hat{g}_i\right)} \\
&= \frac{\left[\operatorname{Cov}\left(y_i, \hat{g}_i\right)\right]^2}{\operatorname{Var}\left(y_i\right)} \\
&= \frac{\left[\operatorname{Cov}\left(y_i, \frac{g_i+e_i}{\operatorname{sd}\left(g_i+e_i\right)}\right)\right]^2}{\operatorname{Var}\left(y_i\right)} \\
&= \frac{\left[\operatorname{Cov}\left(y_i, \frac{g_i}{\operatorname{sd}\left(g_i+e_i\right)}\right)\right]^2}{\operatorname{Var}\left(y_i\right)} \\
&= \frac{1}{\operatorname{Var}\left(g_i+e_i\right)} \frac{\left[\operatorname{Cov}\left(y_i, g_i\right)\right]^2}{\operatorname{Var}\left(y_i\right)} \\
&= \frac{h_{SNP}^2}{\operatorname{Var}\left(g_i+e_i\right)} \\
&= \frac{h_{SNP}^2}{\operatorname{Var}\left(g_i\right)+\operatorname{Var}\left(e_i\right)} \\
&= \frac{h_{SNP}^2}{1+\operatorname{Var}\left(e_i\right)} < h_{SNP}^2 .
\end{aligned}$$

This inequality shows that the predictive power of the PGI is strictly less than the heritability, but the predictive power increases asymptotically towards $h_{SNP}^2$ as the error in the PGI decreases. Also note that this calculation implies

$$\frac{h_{SNP}^2}{R^2} = 1 + \operatorname{Var}\left(e_i\right).$$

We denote this ratio by

$$\rho^2 \equiv \frac{h_{SNP}^2}{R^2}. \tag{5}$$

Using this notation, the PGI can be written as

$$\hat{g}_i = \frac{g_i+e_i}{\operatorname{sd}\left(g_i+e_i\right)} = \frac{g_i+e_i}{\sqrt{1+\operatorname{Var}\left(e_i\right)}} = \frac{g_i+e_i}{\rho}. \tag{6}$$

The error in the PGI will bias any analysis that uses the PGI as a regressor instead of the additive SNP factor. (We use the term *bias* in this case to refer to a difference between expected parameter estimates of a model that includes the additive SNP factor and expected parameter estimates of a model that instead uses the PGI. This is in contrast to whether the PGI itself is a biased predictor of the additive SNP factor or of the phenotype[1].) The "errors-in-variables" bias described here is closely related to what is often referred to as "attenuation bias" because in special cases (such as a univariate regression), measurement error in the regressor attenuates the coefficient on that regressor.

In the following sections, we describe a correction for this bias. We assume that $R^2$ and $h_{SNP}^2$ are known parameters. In practice, these parameters are not known and would need to be estimated. Using estimates of $R^2$ and $h_{SNP}^2$, as opposed to the true value of the parameters, affects the standard errors of the regression estimates. We discuss this issue in section 5 below.

### 2 Measurement-Error-Corrected Estimator

#### 2.1 The Theoretical Regression and the Feasible Regression

Let $\phi_i$ denote a mean zero phenotype of interest, and let $g_i$ denote a standardized additive SNP factor. Note that $\phi_i$ and $g_i$ may correspond to different phenotypes. In addition to $g_i$, the model may include a vector of mean-zero covariates, which we denote by $\mathbf{z}_i$, where $\mathbf{\Sigma}_z \equiv \text{Var}(\mathbf{z}_i)$. The model may also include interactions between $g_i$ and some subset of covariates in $\mathbf{z}_i$. Let $\mathbf{z}_{\text{int},i}$ denote the subset of covariates that we would like to interact with $g_i$ where $\mathbf{\Sigma}_{\text{int},z} \equiv \text{Var}(\mathbf{z}_{\text{int},i})$. We denote the vector of interactions by $\mathbf{w}_i \equiv g_i\mathbf{z}_{int,i}$, where $\mathbf{\Sigma}_w \equiv \text{Var}(\mathbf{w}_i)$ and $\mathbf{\Sigma}_{g,w} \equiv \text{Cov}(g_i, \mathbf{w}_i)$. The *theoretical regression* model we would like to estimate is:

$$\phi_i = g_i\beta_g + \mathbf{w}_i\zeta_g + \mathbf{z}_i\delta_g + \varepsilon_{g,i}. \tag{7}$$

We collect the coefficients of this regression into a single vector, denoted $\alpha_g \equiv (\beta_g, \zeta_g, \delta_g)$. We group all genetic variables (which are the variables affected by the measurement error) using the vector $\mathbf{G}_i \equiv (g_i,\, \mathbf{w}_i)$, where $\mathbf{\Sigma}_G \equiv \text{Var}(\mathbf{G}_i)$ and $\mathbf{\Sigma}_{G,z} \equiv \text{Cov}(\mathbf{G}_i, \mathbf{z}_i)$. The coefficients $\alpha_g \equiv (\beta_g, \zeta_g, \delta_g)$ of the regression (7) are:

$$\begin{aligned}
\alpha_g &= \begin{bmatrix} \text{Var}(\mathbf{G}_i) & \text{Cov}(\mathbf{G}_i, \mathbf{z}_i) \\ & \text{Var}(\mathbf{z}_i) \end{bmatrix}^{-1} \begin{bmatrix} \text{Cov}(\mathbf{G}_i, \phi_i) \\ \text{Cov}(\mathbf{z}_i, \phi_i) \end{bmatrix} \\
&= \begin{bmatrix} \mathbf{\Sigma}_G & \mathbf{\Sigma}_{G,z} \\ & \mathbf{\Sigma}_z \end{bmatrix}^{-1} \begin{bmatrix} \text{Cov}(\mathbf{G}_i, \phi_i) \\ \text{Cov}(\mathbf{z}_i, \phi_i) \end{bmatrix} \\
&= \mathbf{V}_g^{-1} \begin{bmatrix} \text{Cov}(\mathbf{G}_i, \phi_i) \\ \text{Cov}(\mathbf{z}_i, \phi_i) \end{bmatrix},
\end{aligned}$$

where

$$\begin{aligned}
\mathbf{V}_g &\equiv \text{Var}\begin{bmatrix} \mathbf{G}_i \\ \mathbf{z}_i \end{bmatrix} \\
&= \begin{bmatrix} \mathbf{\Sigma}_G & \mathbf{\Sigma}_{G,z} \\ & \mathbf{\Sigma}_z \end{bmatrix}
\end{aligned}$$

is the variance-covariance of all the regressors in the theoretical regression (7).

In practice, however, we do not observe $g_i$ and instead observe the PGI $\hat{g}_i$. Similarly, instead of $\mathbf{w}_i$, we observe $\hat{\mathbf{w}}_i \equiv \hat{g}_i\mathbf{z}_{\text{int},i}$, where $\mathbf{\Sigma}_{\hat{w}} \equiv \text{Var}(\hat{\mathbf{w}}_i)$; and instead of $\mathbf{G}_i$, we observe $\hat{\mathbf{G}}_i = (\hat{g}_i,\, \hat{\mathbf{w}}_i)$, where $\mathbf{\Sigma}_{\hat{G}} \equiv \text{Var}\left(\hat{\mathbf{G}}_i\right)$. We use $\mathbf{\Sigma}_{\hat{G}z} \equiv \text{Cov}\left(\hat{\mathbf{G}}_i, \mathbf{z}_i\right)$ to denote the covariance between $\hat{\mathbf{G}}_i$ and $\mathbf{z}_i$. So the *feasible regression* in this case is

$$\phi_i = \hat{g}_i\beta_{\hat{g}} + \hat{\mathbf{w}}_i\zeta_{\hat{g}} + \mathbf{z}_i\delta_{\hat{g}} + \varepsilon_{\hat{g},i}, \tag{8}$$

where we now subscript the regressors and error term by $\hat{g}$ instead of $g$ to distinguish them from the regressors and error term in the theoretical regression. As before, we collect the coefficients of this regression into a single vector, denoted $\alpha_{\hat{g}} \equiv (\beta_{\hat{g}}, \zeta_{\hat{g}}, \delta_{\hat{g}})$.

#### 2.2 Bias from Estimating the Feasible Regression

To construct a measurement-error correction, we must first derive the relationship between $\alpha_g$ and $\alpha_{\hat{g}}$. The relationship we derive is closely related to results previously derived by Abel (2017). The primary differences between the relationship we derive and that of Abel (2017) are: (i) in our context, we can describe the amount of measurement error as a function of estimable parameters, $h^2_{SNP}$ and $R^2$ (whereas Abel's formula treats the amount of measurement error as known), and (ii) our formula accounts for the standardization of PGIs rather than just assuming that the measurement error is an additive component of the observed value of the regressor.

We begin by writing the coefficients of the feasible regression, $\alpha_{\hat{g}} \equiv (\beta_{\hat{g}}, \zeta_{\hat{g}}, \delta_{\hat{g}})$ in terms of the notation defined above. The coefficient vector is equal to

$$\begin{aligned}
\alpha_{\hat{g}} &= \begin{bmatrix} \text{Var}\left(\hat{\mathbf{G}}_i\right) & \text{Cov}\left(\hat{\mathbf{G}}_i, \mathbf{z}_i\right) \\ & \text{Var}(\mathbf{z}_i) \end{bmatrix}^{-1} \begin{bmatrix} \text{Cov}\left(\hat{\mathbf{G}}_i, \phi_i\right) \\ \text{Cov}(\mathbf{z}_i, \phi_i) \end{bmatrix} \\
&= \begin{bmatrix} \mathbf{\Sigma}_{\hat{G}} & \mathbf{\Sigma}_{\hat{G}z} \\ & \mathbf{\Sigma}_z \end{bmatrix}^{-1} \begin{bmatrix} \text{Cov}\left(\hat{\mathbf{G}}_i, \phi_i\right) \\ \text{Cov}(\mathbf{z}_i, \phi_i) \end{bmatrix} \\
&= \mathbf{V}_{\hat{g}}^{-1} \begin{bmatrix} \text{Cov}\left(\hat{\mathbf{G}}_i, \phi_i\right) \\ \text{Cov}(\mathbf{z}_i, \phi_i) \end{bmatrix},
\end{aligned} \tag{9}$$

where

$$\mathbf{V}_{\hat{g}} \equiv \begin{bmatrix} \mathbf{\Sigma}_{\hat{G}} & \mathbf{\Sigma}_{\hat{G}z} \\ & \mathbf{\Sigma}_z \end{bmatrix}.$$

We now take each term that is related to the PGI and derive its relationship with a term related to the additive SNP factor. Considering the first term in the right column vector of (9),

$$\begin{aligned}
\text{Cov}\left(\hat{\mathbf{G}}_i, \phi_i\right) &= \begin{bmatrix} \text{Cov}\left(\hat{g}_i, \phi_i\right) \\ \text{Cov}\left(\hat{\mathbf{w}}_i, \phi_i\right) \end{bmatrix} \\
&= \begin{bmatrix} \text{Cov}\left(\frac{g_i+e_i}{\rho}, \phi_i\right) \\ \text{Cov}\left(\frac{g_i+e_i}{\rho}\mathbf{z}_{int,i}, \phi_i\right) \end{bmatrix} \\
&= \frac{1}{\rho}\begin{bmatrix} \text{Cov}\left(g_i, \phi_i\right) + \text{Cov}\left(e_i, \phi_i\right) \\ \text{Cov}\left(g_i\mathbf{z}_{int,i}, \phi_i\right) + \text{Cov}\left(e_i\mathbf{z}_{int,i}, \phi_i\right) \end{bmatrix} \\
&= \frac{1}{\rho}\begin{bmatrix} \text{Cov}\left(g_i, \phi_i\right) \\ \text{Cov}\left(g_i\mathbf{z}_{int,i}, \phi_i\right) \end{bmatrix} \\
&= \frac{1}{\rho}\begin{bmatrix} \text{Cov}\left(g_i, \phi_i\right) \\ \text{Cov}\left(\mathbf{w}_i, \phi_i\right) \end{bmatrix} \\
&= \frac{1}{\rho}\text{Cov}\left(\mathbf{G}_i, \phi_i\right).
\end{aligned}$$

Note that in the fourth line, we have used the assumption that $e_i$ is independent of all other varables. The above calculation implies that

$$\begin{bmatrix} \text{Cov}\left(\hat{\mathbf{G}}_i, \phi_i\right) \\ \text{Cov}\left(\mathbf{z}_i, \phi_i\right) \end{bmatrix} = \mathbf{P}^{-1}\begin{bmatrix} \text{Cov}\left(\mathbf{G}_i, \phi_i\right) \\ \text{Cov}\left(\mathbf{z}_i, \phi_i\right) \end{bmatrix}, \tag{10}$$

where

$$\mathbf{P} = \begin{bmatrix} \rho\mathbf{I}_{|G|} & \mathbf{0} \\ & \mathbf{I}_{|z|} \end{bmatrix}.$$

Substituting (10) into (9), we get

$$\begin{aligned}
\alpha_{\hat{g}} &= \mathbf{V}_{\hat{g}}^{-1}\begin{bmatrix} \text{Cov}\left(\hat{\mathbf{G}}_i, \phi_i\right) \\ \text{Cov}\left(\mathbf{z}_i, \phi_i\right) \end{bmatrix} \\
&= \mathbf{V}_{\hat{g}}^{-1}\mathbf{P}^{-1}\begin{bmatrix} \text{Cov}\left(\mathbf{G}_i, \phi_i\right) \\ \text{Cov}\left(\mathbf{z}_i, \phi_i\right) \end{bmatrix} \\
&= \mathbf{V}_{\hat{g}}^{-1}\mathbf{P}^{-1}\mathbf{V}_g\mathbf{V}_g^{-1}\begin{bmatrix} \text{Cov}\left(\mathbf{G}_i, \phi_i\right) \\ \text{Cov}\left(\mathbf{z}_i, \phi_i\right) \end{bmatrix} \\
&= \mathbf{V}_{\hat{g}}^{-1}\mathbf{P}^{-1}\mathbf{V}_g\alpha_g. \qquad (11)
\end{aligned}$$

We now need to derive the relationship between $\mathbf{V}_{\hat{g}}$ and $\mathbf{V}_g$. We begin by calculating

$$\begin{aligned}
\Sigma_{\hat{G}} &= \begin{bmatrix} \text{Var}\left(\hat{g}_i\right) & \text{Cov}\left(\hat{\mathbf{w}}_i, \hat{g}_i\right) \\ & \text{Var}\left(\hat{\mathbf{w}}_i\right) \end{bmatrix} \\
&= \begin{bmatrix} \text{Var}\left(\hat{g}_i\right) & \text{Cov}\left(\hat{g}_i\mathbf{z}_{\text{int},i}, \hat{g}_i\right) \\ & \text{Var}\left(\hat{g}_i\mathbf{z}_{\text{int},i}\right) \end{bmatrix} \\
&= \begin{bmatrix} \text{Var}\left(\frac{g_i+e_i}{\rho}\right) & \text{Cov}\left(\frac{g_i+e_i}{\rho}\mathbf{z}_{\text{int},i}, \frac{g_i+e_i}{\rho}\right) \\ & \text{Var}\left(\frac{g_i+e_i}{\rho}\mathbf{z}_{\text{int},i}\right) \end{bmatrix} \\
&= \left(\frac{1}{\rho}\mathbf{I}_{|G|}\right)\begin{bmatrix} \text{Var}\left(g_i\right) + \text{Var}\left(e_i\right) & \text{Cov}\left(g_i\mathbf{z}_{\text{int},i}, g_i\right) + \text{Cov}\left(e_i\mathbf{z}_{\text{int},i}, e_i\right) \\ & \text{Var}\left(g_i\mathbf{z}_{\text{int},i}\right) + \text{Var}\left(e_i\mathbf{z}_{\text{int},i}\right) \end{bmatrix}\left(\frac{1}{\rho}\mathbf{I}_{|G|}\right) \\
&= \left(\frac{1}{\rho}\mathbf{I}_{|G|}\right)\left(\mathbf{\Sigma}_G + \mathbf{\Omega}_G\right)\left(\frac{1}{\rho}\mathbf{I}_{|G|}\right),
\end{aligned}$$

where

$$\mathbf{\Omega}_G \equiv \begin{bmatrix} \text{Var}(e_i) & \text{Cov}(e_i \mathbf{z}_{\text{int},i}, e_i) \\ & \text{Var}(e_i \mathbf{z}_{\text{int},i}) \end{bmatrix}$$

is the component of the variance-covariance matrix of $\hat{\mathbf{G}}_i$ that is due to error before standardization. Notice that, in the fourth line above, we have again used the assumption of the independence of $e_i$. Defining

$$\mathbf{\Omega} \equiv \begin{bmatrix} \mathbf{\Omega}_G & \mathbf{0} \\ & \mathbf{0} \end{bmatrix},$$

we have

$$\mathbf{V}_{\hat{g}} = \mathbf{P}^{-1} (\mathbf{V}_g + \mathbf{\Omega}) \mathbf{P}^{-1}. \tag{12}$$

Finally, substituting (12) into (11) gives us an equation for how the coefficients from the feasible regression, $\alpha_{\hat{g}}$, are biased relative to the coefficients from the theoretical regression, $\alpha_g$:

$$\begin{aligned} \alpha_{\hat{g}} &= \mathbf{V}_{\hat{g}}^{-1} \mathbf{P}^{-1} \mathbf{V}_g \alpha_g \\ &= \left[\mathbf{P}^{-1} (\mathbf{V}_g + \mathbf{\Omega}) \mathbf{P}^{-1}\right]^{-1} \mathbf{P}^{-1} \mathbf{V}_g \alpha_g \\ &= \mathbf{P} (\mathbf{V}_g + \mathbf{\Omega})^{-1} \mathbf{V}_g \alpha_g. \end{aligned} \tag{13}$$

#### 2.3 Estimator for Coefficients from the Theoretical Regression

Equation (13) can be rearranged to yield the simple regression-disattenuation estimator mentioned in the main text:

$$\alpha_{corr} \equiv \mathbf{V}_g^{-1} (\mathbf{V}_g + \mathbf{\Omega}) \mathbf{P}^{-1} \alpha_{\hat{g}}. \tag{14}$$

This estimator, however, is written in terms of $\mathbf{V}_g$ and $\mathbf{V}_g + \mathbf{\Omega}$, which are unobserved. To obtain the estimator we implement, we now derive expressions for $\mathbf{V}_g$ and $\mathbf{V}_g + \mathbf{\Omega}$ in terms of estimable quantities.

Beginning with $\mathbf{V}_g + \mathbf{\Omega}$, equation (12) gives us

$$\begin{aligned} (\mathbf{V}_g + \mathbf{\Omega}) &= \mathbf{P}\mathbf{P}^{-1} (\mathbf{V}_g + \mathbf{\Omega}) \mathbf{P}^{-1} \mathbf{P} \\ &= \mathbf{P} \mathbf{V}_{\hat{g}} \mathbf{P}. \end{aligned} \tag{15}$$

Now, turning to $\mathbf{V}_g$, we begin by deriving expressions for $\mathbf{\Sigma}_G$ and $\mathbf{\Sigma}_{G,z}$, which are quadrants of the matrix $\mathbf{V}_g$. We calculate

$$\begin{aligned} \mathbf{\Sigma}_G &= \begin{bmatrix} \text{Var}(g_i) & \text{Cov}(\mathbf{w}_i, g_i) \\ & \text{Var}(\mathbf{w}_i) \end{bmatrix} \\ &= \begin{bmatrix} \text{Var}(g_i) & \text{Cov}(\mathbf{w}_i, g_i) - \rho^2 \text{Cov}(\hat{\mathbf{w}}_i, \hat{g}_i) + \rho^2 \text{Cov}(\hat{\mathbf{w}}_i, \hat{g}_i) \\ & \text{Var}(\mathbf{w}_i) - \rho^2 \text{Var}(\hat{\mathbf{w}}_i) + \rho^2 \text{Var}(\hat{\mathbf{w}}_i) \end{bmatrix} \\ &= \begin{bmatrix} \text{Var}(g_i) & \text{Cov}(\mathbf{w}_i, g_i) - \rho^2 \text{Cov}\left(\frac{g_i + e_i}{\rho} \mathbf{z}_{\text{int},i}, \frac{g_i + e_i}{\rho}\right) + \rho^2 \text{Cov}(\hat{\mathbf{w}}_i, \hat{g}_i) \\ & \text{Var}(\mathbf{w}_i) - \rho^2 \text{Var}\left(\frac{g_i + e_i}{\rho} \mathbf{z}_{\text{int},i}\right) + \rho^2 \text{Var}(\hat{\mathbf{w}}_i) \end{bmatrix} \\ &= \begin{bmatrix} \text{Var}(g_i) & \text{Cov}(\mathbf{w}_i, g_i) - \text{Cov}(g_i \mathbf{z}_{\text{int},i}, g_i) - \text{Cov}(e_i \mathbf{z}_{\text{int},i}, e_i) + \rho^2 \text{Cov}(\hat{\mathbf{w}}_i, \hat{g}_i) \\ & \text{Var}(\mathbf{w}_i) - \text{Var}(g_i \mathbf{z}_{\text{int},i}) - \text{Var}(e_i \mathbf{z}_{\text{int},i}) + \rho^2 \text{Var}(\hat{\mathbf{w}}_i) \end{bmatrix} \\ &= \begin{bmatrix} \text{Var}(g_i) & \text{Cov}(\mathbf{w}_i, g_i) - \text{Cov}(\mathbf{w}_i, g_i) - \text{Var}(e_i) \mathbb{E}(\mathbf{z}_{\text{int},i}) + \rho^2 \text{Cov}(\hat{\mathbf{w}}_i, \hat{g}_i) \\ & \text{Var}(\mathbf{w}_i) - \text{Var}(\mathbf{w}_i) - \text{Var}(e_i) \text{Var}(\mathbf{z}_{\text{int},i}) + \rho^2 \text{Var}(\hat{\mathbf{w}}_i) \end{bmatrix} \\ &= \begin{bmatrix} 1 & \rho^2 \text{Cov}(\hat{\mathbf{w}}_i, \hat{g}_i) - (\rho^2 - 1) \mathbb{E}(\mathbf{z}_{\text{int},i}) \\ & \rho^2 \mathbf{\Sigma}_{\hat{w}} - (\rho^2 - 1) \mathbf{\Sigma}_{\text{int},z} \end{bmatrix}. \end{aligned} \tag{16}$$

Also,

$$\begin{aligned}
\mathbf{\Sigma}_{\hat{G},z} &= \begin{bmatrix} \text{Cov}\left(\hat{g}_i, \mathbf{z}_i\right) \\ \text{Cov}\left(\hat{\mathbf{w}}_i, \mathbf{z}_i\right) \end{bmatrix} \\
&= \begin{bmatrix} \text{Cov}\left(\hat{g}_i, \mathbf{z}_i\right) \\ \text{Cov}\left(\hat{g}_i \mathbf{z}_{\text{int},i}, \mathbf{z}_i\right) \end{bmatrix} \\
&= \begin{bmatrix} \text{Cov}\left(\frac{g_i+e_i}{\rho}, \mathbf{z}_i\right) \\ \text{Cov}\left(\frac{g_i+e_i}{\rho}\mathbf{z}_{\text{int},i}, \mathbf{z}_i\right) \end{bmatrix} \\
&= \frac{1}{\rho}\begin{bmatrix} \text{Cov}\left(g_i, \mathbf{z}_i\right) \\ \text{Cov}\left(g_i \mathbf{z}_{\text{int},i}, \mathbf{z}_i\right) \end{bmatrix} \\
&= \frac{1}{\rho}\begin{bmatrix} \text{Cov}\left(g_i, \mathbf{z}_i\right) \\ \text{Cov}\left(\mathbf{w}_i, \mathbf{z}_i\right) \end{bmatrix} \\
&= \frac{1}{\rho}\mathbf{\Sigma}_{G,z}.
\end{aligned}$$

Hence,

$$\mathbf{\Sigma}_{G,z} = \rho \mathbf{\Sigma}_{\hat{G},z}. \tag{17}$$

Equations (16) and (17) then give us an expression for $\mathbf{V}_g$ in terms of observables:

$$\begin{aligned}
\mathbf{V}_g &= \begin{bmatrix} \mathbf{\Sigma}_G & \rho\mathbf{\Sigma}_{\hat{G},z} \\ & \mathbf{\Sigma}_z \end{bmatrix} \\
&= \mathbf{P}\begin{bmatrix} \frac{1}{\rho^2}\mathbf{\Sigma}_G & \mathbf{\Sigma}_{\hat{G},z} \\ & \mathbf{\Sigma}_z \end{bmatrix}\mathbf{P},
\end{aligned} \tag{18}$$

where the estimable expression for $\mathbf{\Sigma}_G$ is given by(16) above.

Substituting these expressions for $\mathbf{V}_g$ and $\mathbf{V}_g + \mathbf{\Omega}$, equations (18) and (15), into the estimator, equation (14), gives us our estimator in terms of estimable quantities:

$$\begin{aligned}
\alpha_{\text{corr}} &\equiv \mathbf{V}_g^{-1}\left(\mathbf{V}_g + \mathbf{\Omega}\right)\mathbf{P}^{-1}\alpha_{\hat{g}} \\
&= \left(\mathbf{P}\begin{bmatrix} \frac{1}{\rho^2}\mathbf{\Sigma}_G & \mathbf{\Sigma}_{\hat{G},z} \\ & \mathbf{\Sigma}_z \end{bmatrix}\mathbf{P}\right)^{-1}\left(\mathbf{P}\mathbf{V}_{\hat{g}}\mathbf{P}\right)\mathbf{P}^{-1}\alpha_{\hat{g}} \\
&= \mathbf{P}^{-1}\begin{bmatrix} \frac{1}{\rho^2}\mathbf{\Sigma}_G & \mathbf{\Sigma}_{\hat{G},z} \\ & \mathbf{\Sigma}_z \end{bmatrix}^{-1}\mathbf{V}_{\hat{g}}\alpha_{\hat{g}} \\
&= \mathbf{C}\alpha_{\hat{g}},
\end{aligned} \tag{19}$$

where

$$\mathbf{C} \equiv \mathbf{P}^{-1}\begin{bmatrix} \frac{1}{\rho^2}\mathbf{\Sigma}_G & \mathbf{\Sigma}_{\hat{G},z} \\ & \mathbf{\Sigma}_z \end{bmatrix}^{-1}\mathbf{V}_{\hat{g}}. \tag{20}$$

#### 2.4 Standard Errors

To obtain standard errors for our estimator, note that equation (19) implies

$$\text{Var}\left(\alpha_{\text{corr}}\right) = \mathbf{C}\mathbf{A}_{\hat{g}}\mathbf{C}'. \tag{21}$$

where $\mathbf{A}_{\hat{g}} \equiv \text{Var}\left(\alpha_{\hat{g}}\right)$. We calculate standard errors by taking the square root of the diagonal of this matrix.

#### 2.5 Two Special Cases

To understand the intuition for the estimator, we consider two special cases. First consider a univariate regression of the phenotype on the PGI. In this case, $\mathbf{G}_i = [g_i]$ and $\mathbf{z}_i$ is empty. Thus

$$\begin{aligned}\alpha_{\text{corr}} &= \left[\frac{1}{\rho}\right]\left[\frac{1}{\rho^2}\right]^{-1}[1]\,\alpha_{\hat{g}} \\ &= \rho\alpha_{\hat{g}}.\end{aligned}$$

In this case, our estimator simply re-inflates the coefficient corresponding to the amount of attenuation due to error in the PGI.

Turning to the standard error, by (21), the sampling variance of the corrected estimate is

$$\begin{aligned}\text{Var}\left(\alpha_{\text{corr}}\right) &= \mathbf{C}\mathbf{A}_{\hat{g}}\mathbf{C}' \\ &= \rho\text{Var}\left(\alpha_{\hat{g}}\right)\rho \\ &= \rho^2\text{Var}\left(\alpha_{\hat{g}}\right).\end{aligned}$$

This means the standard error of the corrected estimate is

$$\text{s.e.}\left(\hat{\alpha}_{corr}\right) = \rho\ \text{s.e.}\left(\hat{\alpha}_{\hat{g}}\right).$$

Since the standard error is inflated by exactly the same factor $\rho$ as the regression coefficient, the $t$-statistic and $p$-value of the measurement-error-corrected regression coefficient remains the same as without the measurement-error correction.

As a second special case, consider a multivariate regression with a single covariate that is independent of the PGI and an interaction between the covariate and the PGI. In this case, we have

$$\begin{aligned}\alpha_{\text{corr}} &= \begin{bmatrix}\frac{1}{\rho} & 0 & 0 \\ & \frac{1}{\rho} & 0 \\ & & 1\end{bmatrix}\begin{bmatrix}\frac{1}{\rho^2} & 0 & 0 \\ & \frac{1}{\rho^2}\Sigma_z & 0 \\ & & \Sigma_z\end{bmatrix}^{-1}\begin{bmatrix}1 & 0 & 0 \\ & \Sigma_z & 0 \\ & & \Sigma_z\end{bmatrix}\alpha_{\hat{g}} \\ &= \begin{bmatrix}\rho & 0 & 0 \\ & \rho & 0 \\ & & 1\end{bmatrix}\alpha_{\hat{g}},\end{aligned}$$

where the first element of $\alpha_{\hat{g}}$ is the coefficient associated with the PGI, the second element is the coefficient associated with the interaction, and the third element is the coefficient associated with the covariate. Similar to the univariate special case, the estimator in the simple, independent gene-by-environment interaction case with no other covariates simply inflates the coefficients corresponding to covariates related to the PGI. We conclude from this special that that as a first approximation, we should expect that the estimator will inflate each of the coefficients associated with the PGI or its interactions by $\rho$. The estimator will deviate from this benchmark to the extent that the PGI is correlated with the interacted environmental factor or any other covariates included in the model.

Using (21), we calculate the variance of the corrected estimates:

$$\begin{aligned}\text{Var}\left(\alpha_{\text{corr}}\right) &= \mathbf{C}\mathbf{A}_{\hat{g}}\mathbf{C}' \\ &= \begin{bmatrix}\rho & 0 & 0 \\ & \rho & 0 \\ & & 1\end{bmatrix}\text{Var}\left(\alpha_{\hat{g}}\right)\begin{bmatrix}\rho & 0 & 0 \\ & \rho & 0 \\ & & 1\end{bmatrix} \\ &= \begin{bmatrix}\rho^2\text{Var}\left(\alpha_{\hat{g},1}\right) & \rho^2\text{Cov}\left(\alpha_{\hat{g},1},\alpha_{\hat{g},2}\right) & \rho\text{Cov}\left(\alpha_{\hat{g},1},\alpha_{\hat{g},3}\right) \\ & \rho^2\text{Var}\left(\alpha_{\hat{g},2}\right) & \rho\text{Cov}\left(\alpha_{\hat{g},2},\alpha_{\hat{g},3}\right) \\ & & \text{Var}\left(\alpha_{\hat{g},3}\right)\end{bmatrix},\end{aligned}$$

where $\alpha_{\hat{g},1}$, $\alpha_{\hat{g},2}$, and $\alpha_{\hat{g},3}$ are the three elements of $\alpha_{\hat{g}}$. The standard errors are the square root of the diagonal of this matrix, giving us

$$\text{s.e.}\left(\hat{\alpha}_{corr}\right) = \begin{bmatrix} \sqrt{\rho^2 \text{Var}\left(\hat{\alpha}_{\hat{g},1}\right)} \\ \sqrt{\rho^2 \text{Var}\left(\hat{\alpha}_{\hat{g},2}\right)} \\ \sqrt{\text{Var}\left(\hat{\alpha}_{\hat{g},3}\right)} \end{bmatrix}$$
$$= \begin{bmatrix} \rho & 0 & 0 \\ & \rho & 0 \\ & & 1 \end{bmatrix} \text{Var}\left(\hat{\alpha}_{\hat{g}}\right).$$

Thus, each of the standard errors of the corrected estimates is inflated by exactly the same proportion as the inflation of its corresponding corrected estimates. Therefore, in this special case, the $t$-statistics and $p$-values of all three measurement-error-corrected regression coefficients remain the same as without the measurement-error correction.

### 3 Implementation of the Estimator

In the derivation above, we have expressed everything in terms of population parameters. In order to obtain a consistent estimator of $\alpha_{\text{corr}}$ and its standard error, we must write them in terms of the data that we observe.

First, consider the parameter $\rho$. Our estimator $\hat{\rho}$ is

$$\hat{\rho} \equiv \sqrt{\frac{\hat{h}^2_{SNP}}{\hat{R}^2}}.$$

The value $\hat{h}^2_{SNP}$ is an estimate of SNP heritability of the phenotype $y_i$ in the prediction dataset, based on the same set of $J$ SNPs that make up the PGI. The value $\hat{R}^2$ is the estimated predictive power of the PGI for $y_i$ in the prediction dataset.

Note that $\hat{h}^2_{SNP}$ and $\hat{R}^2$ each correspond to the PGI phenotype $y_i$ rather than the (possibly different) phenotype in the regression analysis $\phi_i$. If the phenotype $y_i$ is not available in the prediction dataset or if the sample is too small to obtain reliable estimates, $\hat{h}^2_{SNP}$ and $\hat{R}^2$, $\hat{\rho}$ could instead be estimated from a different sample without introducing any bias as long as the genetic correlation of $y_i$ is perfect between the two samples. (The heritability may differ in the samples, but the genetic correlation must be one. This may happen if the individuals are drawn from the same population in the two samples, but the phenotype is measured with greater error in one of them.) Given that a researcher would choose to do this only in the absence of enough data on $y_i$ in the regression sample, perfect genetic correlation cannot be reliably tested and would therefore become an important assumption underlying use of the correction.

Turning to the other parameters besides $\rho$, in all cases we replace the population variance-covariance matrices with the consistent (sample-analog) estimates of these matrices. For example,

$$\hat{\mathbf{\Sigma}}_z \equiv \frac{1}{N}\mathbf{Z}'\mathbf{Z},$$

were $N$ is the sample size in the regression sample and $\mathbf{Z}$ is the $N \times |z_i|$ matrix of covariates in the regression.

Since $\hat{\rho}$ and each variance-covariance matrix is a consistent estimator of its population counterpart, $\hat{\alpha}_{\text{corr}}$ is a consistent estimator of $\alpha_{\text{corr}}$.

### 4 Assumption That $e_i$ is Uncorrelated With Other Variables

Recall from equation (4), we have expressed the PGI as

$$\hat{g}_i = \frac{g_i + e_i}{\text{sd}\left(g_i + e_i\right)},$$

where, by construction, $\text{Cov}\left(g_i, e_i\right) = 0$. We assumed that $e_i$ is uncorrelated with all other covariates in the model. In this section, we show that if the SNP weights for the PGI are unbiased estimates of the

SNP weights for the additive SNP factor, then this uncorrelatedness assumption is exactly true. We then show that when SNP weights for the PGI are estimated using LDpred-inf—which is the method we use for the Repository and which does not generate unbiased estimates—given typical parameter values for PGIs in the Repository, the bias in our measurement-error-corrected estimator due to the violation of the uncorrelatedness assumption is negligible.

#### 4.1 Uncorrelatedness Is Implied When Unbiased Estimates of $\gamma_j$ Are Used

A sufficient condition for our $\text{Cov}(g_i, e_i) = 0$ assumption to hold is that the SNP weights for the PGI are unbiased estimates of the SNP weights for the additive SNP factor. To state it more formally, recall that the standardized additive SNP factor is

$$g_i = \frac{\mathbf{x}_i \gamma}{\text{sd}(\mathbf{x}_i \gamma)},$$

and the PGI is

$$\hat{g}_i = \frac{\mathbf{x}_i \hat{\gamma}}{\text{sd}(\mathbf{x}_i \hat{\gamma})}.$$

The sufficient condition is that $\hat{\gamma}$ is an unbiased estimate of $\gamma$. This would be the case, for example, if $\hat{\gamma}$ is estimated by ordinary least squares or logistic regression (rather than a Bayesian approach, such as LDpred, which tends to shrink coefficient estimates relative to those from ordinary least squares). This is roughly equivalent to what is done when PGIs are constructed using "Pruning and Thresholding" methods as long as the PGI weights are estimated in a different sample than the sample used to select the SNPs that are included in the PGI. (Note, however, that because "Pruning and Thresholding" methods construct a PGI using fewer SNPs, the resulting PGI is proxying for an additive SNP factor that is based on fewer SNPs and hence has a lower $h^2_{SNP}$.) Because $\hat{\gamma}$ is unbiased,

$$\hat{\gamma} = \gamma + \mathbf{e}_\gamma,$$

where $\mathbb{E}(\mathbf{e}_\gamma) = \mathbf{0}$. Since $\mathbf{e}_\gamma$ is sampling error, $\mathbf{e}_\gamma$ is independent of all variables in independent samples. Therefore, the measurement error in the PGI, $e_i = \mathbf{x}_i \mathbf{e}_\gamma$, is also independent of all variables in independent samples.

#### 4.2 Magnitude of the Bias When $\gamma_j$ Is Estimated Using LDpred-inf

For the Repository, we construct the PGI weights using LDpred-inf. To be precise about this method, it is helpful to express the length-$N$ vector of phenotype values for the $N$ individuals in the discovery sample as

$$\mathbf{Y} = \mathbf{X}\gamma + \mathbf{E},$$

where $\mathbf{X}$ is the $N \times K$ matrix of $K$ genotypes, and $\mathbf{E}$ is a vector of residuals, which is uncorrelated with $\mathbf{X}$. The LDpred-inf estimator for $\hat{\gamma}$ is:

$$\hat{\gamma} = \left( \mathbf{X}'\mathbf{X} + \frac{1}{\sigma^2_\gamma} \mathbf{I} \right)^{-1} \mathbf{X}'\mathbf{Y},$$

where $\sigma^2_\gamma \equiv \text{Var}(\gamma) = \frac{h^2_{SNP}}{K}$ is the prior variance of the additive SNP factor weights. This is equivalent to the ridge regression estimator, with a particular choice of the regularization parameter. This estimator reduces the problem of multicollinearity (due to LD) at the cost of some bias in the estimates. We can calculate the relationship between $\hat{\gamma}$ and $\gamma$ as

$$\begin{aligned}
\hat{\gamma} &= \left(\mathbf{X}'\mathbf{X} + \frac{1}{\sigma_\gamma^2}\mathbf{I}\right)^{-1}\mathbf{X}'\mathbf{Y} \\
&= \left(\mathbf{X}'\mathbf{X} + \frac{1}{\sigma_\gamma^2}\mathbf{I}\right)^{-1}\mathbf{X}'\left(\mathbf{X}\gamma + \mathbf{E}\right) \\
&= \left(N\hat{\Sigma}_X + \frac{1}{\sigma_\gamma^2}\mathbf{I}\right)^{-1}\left(N\hat{\Sigma}_X\right)\gamma + e_{\hat{\gamma}} \\
&= \left(\hat{\Sigma}_X + \frac{1}{N\sigma_\gamma^2}\mathbf{I}\right)^{-1}\hat{\Sigma}_X\gamma + e_{\hat{\gamma}},
\end{aligned} \tag{22}$$

where $\hat{\Sigma}_X \equiv \frac{1}{N}\mathbf{X}'\mathbf{X}$ is the sample variance-covariance matrix of $\mathbf{X}$ and $e_{\hat{\gamma}} \equiv \left(\mathbf{X}'\mathbf{X} + \frac{1}{\sigma_\gamma^2}\mathbf{I}\right)^{-1}\mathbf{X}'\mathbf{E}$ is the estimation error of $\hat{\gamma}$.

To evaluate the magnitude of the bias in finite samples, we quantify it in a simple case where we regress the phenotype $\phi_i$ on the standardized additive SNP factor $g_i$ and a single (scalar) covariate $z_i$. Without loss of generality, we orient $z_i$ such that $g_i$ and $z_i$ have positive covariance. As in the main text, we use $\alpha_g$ to denote the coefficients of this theoretical regression:

$$\phi_i = \begin{bmatrix} g_i & z_i \end{bmatrix}\alpha_g + \varepsilon_i.$$

The coefficients from the feasible regression are

$$\alpha_{\hat{g}} = \begin{bmatrix} \mathrm{Var}(\hat{g}_i) & \mathrm{Cov}(\hat{g}_i, z_i) \\ & \mathrm{Var}(z_i) \end{bmatrix}^{-1}\begin{bmatrix} \mathrm{Cov}(\hat{g}_i, \phi_i) \\ \mathrm{Cov}(z_i, \phi_i) \end{bmatrix}.$$

Our measurement-error-corrected estimator is

$$\begin{aligned}
\alpha_{\text{corr}} &= \begin{bmatrix} \frac{1}{\rho} & 0 \\ & 1 \end{bmatrix}\begin{bmatrix} \frac{1}{\rho^2} & \mathrm{Cov}(\hat{g}_i, z_i) \\ & \mathrm{Var}(z_i) \end{bmatrix}^{-1}\begin{bmatrix} \mathrm{Var}(\hat{g}_i) & \mathrm{Cov}(\hat{g}_i, z_i) \\ & \mathrm{Var}(z_i) \end{bmatrix}\alpha_{\hat{g}} \\
&= \begin{bmatrix} \frac{1}{\rho} & 0 \\ & 1 \end{bmatrix}\begin{bmatrix} \frac{1}{\rho^2} & \mathrm{Cov}(\hat{g}_i, z_i) \\ & \mathrm{Var}(z_i) \end{bmatrix}^{-1}\begin{bmatrix} \mathrm{Cov}(\hat{g}_i, \phi_i) \\ \mathrm{Cov}(z_i, \phi_i) \end{bmatrix} \\
&= \begin{bmatrix} \frac{1}{\rho} & 0 \\ & 1 \end{bmatrix}\begin{bmatrix} \frac{1}{\rho^2} & \mathrm{Cov}(\hat{g}_i, z_i) \\ & \mathrm{Var}(z_i) \end{bmatrix}^{-1}\begin{bmatrix} \mathrm{Cov}\left(\hat{g}_i, \begin{bmatrix} g_i & z_i \end{bmatrix}\alpha + \varepsilon_i\right) \\ \mathrm{Cov}\left(z_i, \begin{bmatrix} g_i & z_i \end{bmatrix}\alpha + \varepsilon_i\right) \end{bmatrix} \\
&= \begin{bmatrix} \frac{1}{\rho} & 0 \\ & 1 \end{bmatrix}\begin{bmatrix} \frac{1}{\rho^2} & \mathrm{Cov}(\hat{g}_i, z_i) \\ & \mathrm{Var}(z_i) \end{bmatrix}^{-1}\begin{bmatrix} \mathrm{Cov}(\hat{g}_i, g_i) & \mathrm{Cov}(\hat{g}_i, z_i) \\ \mathrm{Cov}(g_i, z_i) & \mathrm{Var}(z_i) \end{bmatrix}\alpha \\
&= \frac{1}{\frac{\mathrm{Var}(z_i)}{\rho^2} - \mathrm{Cov}(\hat{g}_i, z_i)^2}\begin{bmatrix} \frac{1}{\rho} & 0 \\ & 1 \end{bmatrix}\begin{bmatrix} \mathrm{Var}(z_i) & -\mathrm{Cov}(\hat{g}_i, z_i) \\ & \frac{1}{\rho^2} \end{bmatrix}\begin{bmatrix} \mathrm{Cov}(\hat{g}_i, g_i) & \mathrm{Cov}(\hat{g}_i, z_i) \\ \mathrm{Cov}(g_i, z_i) & \mathrm{Var}(z_i) \end{bmatrix}\alpha \\
&= \frac{1}{\frac{\mathrm{Var}(z_i)}{\rho^2} - \mathrm{Cov}(\hat{g}_i, z_i)^2}\begin{bmatrix} \frac{1}{\rho}\mathrm{Var}(z_i) & -\frac{1}{\rho}\mathrm{Cov}(\hat{g}_i, z_i) \\ -\mathrm{Cov}(\hat{g}_i, z_i) & \frac{1}{\rho^2} \end{bmatrix}\begin{bmatrix} \mathrm{Cov}(\hat{g}_i, g_i) & \mathrm{Cov}(\hat{g}_i, z_i) \\ \mathrm{Cov}(g_i, z_i) & \mathrm{Var}(z_i) \end{bmatrix}\alpha \\
&= \begin{bmatrix} \frac{\frac{1}{\rho}\mathrm{Var}(z_i)\mathrm{Cov}(\hat{g}_i, g_i) - \frac{1}{\rho}\mathrm{Cov}(g_i, z_i)\mathrm{Cov}(\hat{g}_i, z_i)}{\frac{\mathrm{Var}(z_i)}{\rho^2} - \mathrm{Cov}(\hat{g}_i, z_i)^2} & 0 \\ \frac{\frac{1}{\rho^2}\mathrm{Cov}(g_i, z_i) - \mathrm{Cov}(\hat{g}_i, z_i)\mathrm{Cov}(\hat{g}_i, g_i)}{\frac{\mathrm{Var}(z_i)}{\rho^2} - \mathrm{Cov}(\hat{g}_i, z_i)^2} & 1 \end{bmatrix}\alpha \\
&= \begin{bmatrix} \frac{\frac{1}{\rho}\mathrm{Var}(z_i)\mathrm{Cov}(\hat{g}_i, g_i) - \frac{1}{\rho}\mathrm{Cov}(g_i, z_i)\mathrm{Cov}(\hat{g}_i, z_i)}{\frac{\mathrm{Var}(z_i)}{\rho^2} - \mathrm{Cov}(\hat{g}_i, z_i)^2} & 0 \\ \frac{\frac{1}{\rho^2}\mathrm{Cov}(g_i, z_i) - \mathrm{Cov}(\hat{g}_i, z_i)\mathrm{Cov}(\hat{g}_i, g_i)}{\frac{\mathrm{Var}(z_i)}{\rho^2} - \mathrm{Cov}(\hat{g}_i, z_i)^2} & 1 \end{bmatrix}\alpha.
\end{aligned}$$

Using $\text{Cov}(\hat{g}_i, g_i) = \text{Cov}\left(\frac{g_i+e_i}{\rho}, g_i\right) = \frac{1}{\rho}$, we have

$$\begin{aligned}
\alpha_{\text{corr}} &= \begin{bmatrix} \frac{\frac{1}{\rho^2}\text{Var}(z_i) - \frac{1}{\rho}\text{Cov}(g_i,z_i)\text{Cov}(\hat{g}_i,z_i)}{\frac{\text{Var}(z_i)}{\rho^2} - \text{Cov}(\hat{g}_i,z_i)^2} & 0 \\ \frac{\frac{1}{\rho^2}\text{Cov}(g_i,z_i) - \frac{1}{\rho}\text{Cov}(\hat{g}_i,z_i)}{\frac{\text{Var}(z_i)}{\rho^2} - \text{Cov}(\hat{g}_i,z_i)^2} & 1 \end{bmatrix} \alpha \\
&= \begin{bmatrix} \frac{\frac{1}{\rho^2}\text{Var}(z_i) - \text{Cov}(\hat{g}_i,z_i)^2 + \text{Cov}(\hat{g}_i,z_i)^2 - \frac{1}{\rho}\text{Cov}(g_i,z_i)\text{Cov}(\hat{g}_i,z_i)}{\frac{\text{Var}(z_i)}{\rho^2} - \text{Cov}(\hat{g}_i,z_i)^2} & 0 \\ \frac{\frac{1}{\rho^2}\text{Cov}(g_i,z_i) - \frac{1}{\rho}\text{Cov}(\hat{g}_i,z_i)}{\frac{\text{Var}(z_i)}{\rho^2} - \text{Cov}(\hat{g}_i,z_i)^2} & 1 \end{bmatrix} \alpha \\
&= \begin{bmatrix} 1 + \frac{\text{Cov}(\hat{g}_i,z_i)\left[\text{Cov}(\hat{g}_i,z_i) - \frac{1}{\rho}\text{Cov}(g_i,z_i)\right]}{\frac{\text{Var}(z_i)}{\rho^2} - \text{Cov}(\hat{g}_i,z_i)^2} & 0 \\ \frac{-\frac{1}{\rho}\left[\text{Cov}(\hat{g}_i,z_i) - \frac{1}{\rho}\text{Cov}(g_i,z_i)\right]}{\frac{\text{Var}(z_i)}{\rho^2} - \text{Cov}(\hat{g}_i,z_i)^2} & 1 \end{bmatrix} \alpha.
\end{aligned}$$

This means that the bias is

$$\begin{aligned}
\alpha_{\text{corr}} - \alpha &= \begin{bmatrix} 1 + \frac{\text{Cov}(\hat{g}_i,z_i)\left[\text{Cov}(\hat{g}_i,z_i) - \frac{1}{\rho}\text{Cov}(g_i,z_i)\right]}{\frac{\text{Var}(z_i)}{\rho^2} - \text{Cov}(\hat{g}_i,z_i)^2} & 0 \\ \frac{-\frac{1}{\rho}\left[\text{Cov}(\hat{g}_i,z_i) - \frac{1}{\rho}\text{Cov}(g_i,z_i)\right]}{\frac{\text{Var}(z_i)}{\rho^2} - \text{Cov}(\hat{g}_i,z_i)^2} & 1 \end{bmatrix} \alpha - \alpha \\
&= \begin{bmatrix} \frac{\text{Cov}(\hat{g}_i,z_i)\left[\text{Cov}(\hat{g}_i,z_i) - \frac{1}{\rho}\text{Cov}(g_i,z_i)\right]}{\frac{\text{Var}(z_i)}{\rho^2} - \text{Cov}(\hat{g}_i,z_i)^2} & 0 \\ \frac{-\frac{1}{\rho}\left[\text{Cov}(\hat{g}_i,z_i) - \frac{1}{\rho}\text{Cov}(g_i,z_i)\right]}{\frac{\text{Var}(z_i)}{\rho^2} - \text{Cov}(\hat{g}_i,z_i)^2} & 0 \end{bmatrix} \alpha \\
&= \left[\text{Cov}(\hat{g}_i, z_i) - \frac{1}{\rho}\text{Cov}(g_i, z_i)\right] \begin{bmatrix} \frac{\text{Cov}(\hat{g}_i,z_i)}{\frac{\text{Var}(z_i)}{\rho^2} - \text{Cov}(\hat{g}_i,z_i)^2} \\ \frac{-\frac{1}{\rho}}{\frac{\text{Var}(z_i)}{\rho^2} - \text{Cov}(\hat{g}_i,z_i)^2} \end{bmatrix} \alpha_1, \qquad (23)
\end{aligned}$$

where $\alpha_1$ is the first element of $\alpha$.

We next express the first factor, $\left[\text{Cov}(\hat{g}_i, z_i) - \frac{1}{\rho}\text{Cov}(g_i, z_i)\right]$, in terms of observable or estimable quantities. To do this, consider the best linear predictor of $z_i$ using the same SNPs that make up $g_i$. That predictor would have weights

$$\xi \equiv \arg\min_{\tilde{\xi}} \mathbb{E}\left[\left(y_i - \mathbf{x}_i\tilde{\xi}\right)^2\right]$$

(so $\mathbf{x}_i\xi$ is the additive SNP factor for $z_i$). Therefore,

$$\begin{aligned}
\left[\text{Cov}(\hat{g}_i, z_i) - \frac{1}{\rho}\text{Cov}(g_i, z_i)\right] &= \left[\text{Cov}\left(\frac{x_i\hat{\gamma}}{\rho}, x_i\xi\right) - \frac{1}{\rho}\text{Cov}(x_i\gamma, x_i\xi)\right] \\
&= \frac{1}{\rho}\left[\text{Cov}\left(x_i\left[\left(\hat{\Sigma}_X + \frac{1}{N\sigma_\gamma^2}\mathbf{I}\right)^{-1}\hat{\Sigma}_X\gamma + e_{\hat{\gamma}}\right], x_i\xi\right) - \text{Cov}(x_i\gamma, x_i\xi)\right] \\
&= \frac{1}{\rho}\left[\text{Cov}\left(x_i\left(\hat{\Sigma}_X + \frac{1}{N\sigma_\gamma^2}\mathbf{I}\right)^{-1}\hat{\Sigma}_X\gamma, x_i\xi\right) - \text{Cov}(x_i\gamma, x_i\xi)\right] \\
&= \frac{1}{\rho}\left[\gamma'\hat{\Sigma}_X\left(\hat{\Sigma}_X + \frac{1}{N\sigma_\gamma^2}\mathbf{I}\right)^{-1}\text{Var}(x_i) - \gamma'\text{Var}(x_i)\right]\xi \\
&= \frac{1}{\rho}\gamma'\left[\hat{\Sigma}_X\left(\hat{\Sigma}_X + \frac{1}{N\sigma_\gamma^2}\mathbf{I}\right)^{-1}\Sigma_X - \Sigma_X\right]\xi.
\end{aligned}$$

The second line follows from substituting equation (22) into the first line. By Woodbury's Identity, we have

$$\begin{aligned}\left[\text{Cov}\left(\hat{g}_i, z_i\right) - \frac{1}{\rho}\text{Cov}\left(g_i, z_i\right)\right] &= \frac{1}{\rho}\gamma'\left[\hat{\Sigma}_X\left(\hat{\Sigma}_X^{-1} - \hat{\Sigma}_X^{-1}\left(\hat{\Sigma}_X^{-1} + n\sigma_\gamma^2\mathbf{I}\right)^{-1}\hat{\Sigma}_X^{-1}\right)\Sigma_X - \Sigma_X\right]\xi \\ &= \frac{1}{\rho}\gamma'\left[\hat{\Sigma}_X\hat{\Sigma}_X^{-1}\Sigma_X - \hat{\Sigma}_X\hat{\Sigma}_X^{-1}\left(\hat{\Sigma}_X^{-1} + n\sigma_\gamma^2\mathbf{I}\right)^{-1}\hat{\Sigma}_X^{-1}\Sigma_X - \Sigma_X\right]\xi \\ &= \frac{1}{\rho}\gamma'\left[\Sigma_X - \left(\hat{\Sigma}_X^{-1} + n\sigma_\gamma^2\mathbf{I}\right)^{-1}\hat{\Sigma}_X^{-1}\Sigma_X - \Sigma_X\right]\xi \\ &= -\frac{1}{\rho}\gamma'\left(\hat{\Sigma}_X^{-1} + n\sigma_\gamma^2\mathbf{I}\right)^{-1}\hat{\Sigma}_X^{-1}\Sigma_X\xi. \end{aligned} \tag{24}$$

Next imagine a weighted regression of $\xi$ onto $\gamma$ with weights $\left(\Sigma_X^{-1} + n\sigma_\gamma^2\mathbf{I}\right)^{-1}$. This produces

$$\xi = \frac{\sigma_{\xi\gamma}}{\sigma_\gamma^2}\gamma + \mu,$$

with $\sigma_{\gamma\xi} \equiv \text{Cov}\left(\gamma, \xi\right)$ and with the residual $\mu$ having the property $\gamma'\left(\Sigma_X^{-1} + n\sigma_\gamma^2\mathbf{I}\right)^{-1}\hat{\Sigma}_X^{-1}\Sigma_X\mu = 0$. Since we have oriented $g_i$ and $z_i$ to have positive covariance, $\sigma_{\gamma\xi} > 0$. Substituting both of these equations into (24) gives us

$$\begin{aligned}\left[\text{Cov}\left(\hat{g}_i, z_i\right) - \frac{1}{\rho}\text{Cov}\left(g_i, z_i\right)\right] &= -\frac{1}{\rho}\gamma'\left(\hat{\Sigma}_X^{-1} + n\sigma_\gamma^2\mathbf{I}\right)^{-1}\hat{\Sigma}_X^{-1}\Sigma_X\left(\frac{\sigma_{\xi\gamma}}{\sigma_\gamma^2}\gamma + \mu\right) \\ &= -\frac{1}{\rho}\left[\frac{\sigma_{\xi\gamma}}{\sigma_\gamma^2}\gamma'\left(\Sigma_X^{-1} + n\sigma_\gamma^2\mathbf{I}\right)^{-1}\hat{\Sigma}_X^{-1}\Sigma_X\gamma + \gamma'\left(\Sigma_X^{-1} + n\sigma_\gamma^2\mathbf{I}\right)^{-1}\hat{\Sigma}_X^{-1}\Sigma_X\mu\right] \\ &= -\frac{1}{\rho}\frac{\sigma_{\xi\gamma}}{\sigma_\gamma^2}\gamma'\left(\Sigma_X^{-1} + n\sigma_\gamma^2\mathbf{I}\right)^{-1}\hat{\Sigma}_X^{-1}\Sigma_X\gamma. \end{aligned}$$

Next, in order to put an upper bound on the magnitude of bias, we will show that $\gamma'\left(\Sigma_X^{-1} + n\sigma_\gamma^2\mathbf{I}\right)^{-1}\hat{\Sigma}_X^{-1}\Sigma_X\gamma < \frac{1}{n\sigma_\gamma^2}\gamma'\gamma$. Again using Woodbury's Identity, we have

$$\begin{aligned}\gamma'\left(\Sigma_X^{-1} + n\sigma_\gamma^2\mathbf{I}\right)^{-1}\hat{\Sigma}_X^{-1}\Sigma_X\gamma - \frac{1}{n\sigma_\gamma^2}\gamma'\gamma &= \gamma'\left[\left(\Sigma_X^{-1} + n\sigma_\gamma^2\mathbf{I}\right)^{-1}\hat{\Sigma}_X^{-1}\Sigma_X - \frac{1}{n\sigma_\gamma^2}\mathbf{I}\right]\gamma \\ &= \gamma'\left[\frac{1}{n\sigma_\gamma^2}\mathbf{I} - \left(\frac{1}{n\sigma_\gamma^2}\mathbf{I}\right)\left(\Sigma_X + \frac{1}{n\sigma_\gamma^2}\mathbf{I}\right)^{-1}\left(\frac{1}{n\sigma_\gamma^2}\mathbf{I}\right)\hat{\Sigma}_X^{-1}\Sigma_X - \frac{1}{n\sigma_\gamma^2}\mathbf{I}\right]\gamma \\ &= -\frac{1}{n^2\sigma_\gamma^4}\gamma'\left(\Sigma_X + \frac{1}{n\sigma_\gamma^2}\mathbf{I}\right)^{-1}\hat{\Sigma}_X^{-1}\Sigma_X\gamma \\ &< 0. \end{aligned}$$

The last step here follows because $\Sigma_X$ and $\frac{1}{n\sigma_\gamma^2}\mathbf{I}$ are positive definite matrices. So this implies that

$\gamma' \left(\Sigma_X^{-1} + n\sigma_\gamma^2 \mathbf{I}\right)^{-1} \hat{\Sigma}_X^{-1} \Sigma_X \gamma < \frac{1}{n\sigma_\gamma^2}\gamma'\gamma$. Since both sides of this inequality are positive, this implies that

$$\begin{aligned}
\left[\mathrm{Cov}\left(\hat{g}_i, z_i\right) - \frac{1}{\rho}\mathrm{Cov}\left(g_i, z_i\right)\right] &= -\frac{1}{\rho}\frac{\sigma_{\xi\gamma}}{\sigma_\gamma^2}\gamma' \left(\Sigma_X^{-1} + n\sigma_\gamma^2 \mathbf{I}\right)^{-1} \hat{\Sigma}_X^{-1} \Sigma_X \gamma \\
&> -\frac{1}{\rho}\frac{\sigma_{\xi\gamma}}{\sigma_\gamma^2}\frac{1}{n\sigma_\gamma^2}\gamma'\gamma \\
&= -\frac{1}{\rho}\frac{\sigma_{\xi\gamma}}{\sigma_\gamma^2}\frac{1}{n\sigma_\gamma^2} M_e \sigma_\gamma^2 \\
&= -\frac{1}{\rho}\frac{\sigma_{\xi\gamma}}{\sigma_\gamma^2}\frac{M_e}{n} \\
&= -\frac{1}{\rho} r_{\xi\gamma}\frac{\sigma_\xi}{\sigma_\gamma}\frac{M_e}{n} \\
&= -\frac{1}{\rho} r_{\xi\gamma}\sqrt{\frac{h_z^2}{h_{SNP}^2}}\frac{M_e}{n}.
\end{aligned}$$

where $M_e$ is the effective population size, $r_{\xi\gamma}$ is the correlation of $\xi$ and $\gamma$, and $h_z^2$ is the SNP heritability of $z_i$. The parameters $r_{\xi\gamma}$ and $h_z^2$ are unknown, but the magnitude of this term will be largest when $r_{\xi\gamma} = h_z^2 = 1$. So we have

$$\left[\mathrm{Cov}\left(\hat{g}_i, z_i\right) - \frac{1}{\rho}\mathrm{Cov}\left(g_i, z_i\right)\right] > -\frac{1}{\rho}\frac{M_e}{n h_{SNP}}. \tag{25}$$

Substituting (25) into (23) therefore gives us an upper bound on the magnitude of the bias of the corrected estimates:

$$b_{upper} = -\frac{1}{\rho}\frac{1}{n h_{SNP}} \begin{bmatrix} \frac{\mathrm{Cov}(\hat{g}_i, z_i)}{\frac{\mathrm{Var}(z_i)}{\rho^2} - \mathrm{Cov}(\hat{g}_i, z_i)^2} \\ \frac{-\frac{1}{\rho}}{\frac{\mathrm{Var}(z_i)}{\rho^2} - \mathrm{Cov}(\hat{g}_i, z_i)^2} \end{bmatrix} \alpha_1. \tag{26}$$

Each of the values in this expression is observed or estimable. This means we can replace each of these parameters with their corresponding estimates to approximate the magnitude of the bias. Calculations using equation (26) imply that when Repository PGIs are used (for which the weights are calculated using LDpred-inf), the bias due to the violation of the $\mathrm{Cov}\left(e_i, z_i\right) = 0$ assumption will typically be small.

For example, using values from the Papageorge and Thom application used in this paper, if $z_i$ represents mother's educational attainment, we estimate $\hat{\rho} = 1.51$, $\hat{h}_{SNP}^2 = 0.25$, $n = 293,723$, $\mathrm{Var}\left(z_i\right) = 9.02$, $\mathrm{Cov}\left(\hat{g}_i, z_i\right) = 0.53$ and $\hat{\alpha}_1 = 1.16$. (Note that this value of $\alpha_1$ is actually based on controlling for mother and father's education, but since this exercise is just meant to get a approximation of the order or magnitude of the bias, we have not re-evaluated the model with only one covariate.) Substituting these values into (26) gives us

$$b_{upper} = \begin{bmatrix} -6.50 \times 10^{-7} \\ 8.13 \times 10^{-7} \end{bmatrix}.$$

This is several orders of magnitude smaller than the measurement-error correction.

### 5 Potential Bias in the Standard Errors

The standard errors from (21) ignore the estimation error introduced by using $\hat{\rho}$ rather than $\rho$. We argue here that ignoring this source of uncertainty induces little bias to our standard errors for $\hat{\alpha}_{\mathrm{corr}}$ if $\rho$ is estimated in the same sample as $\alpha_{\hat{g}}$.

Consider the univariate case: regressing the phenotype $\phi_i$ on only the PGI $\hat{g}_i$. In that case,

$$\begin{aligned}
\hat{\alpha}_{\mathrm{corr}} &= \hat{\rho}\hat{\alpha}_{\hat{g}} \\
&= \frac{\hat{h}_{SNP}}{\hat{R}}\hat{\alpha}_{\hat{g}}.
\end{aligned}$$

Note that $\hat{R}$ and $\hat{\alpha}_{\hat{g}}$ correspond to how well the PGI predicts $y_i$ and $\phi_i$, respectively, in sample. For this reason, the error in $\hat{R}$ and $\hat{\alpha}_{\hat{g}}$ will be positively correlated, which will reduce the standard error of $\hat{\alpha}_{\text{corr}}$. In contrast, the error in $\hat{h}_{SNP}$ will also be correlated with the error in $\hat{\alpha}_{\hat{g}}$, which will increase the standard error. In the further simple setting where $\phi_i = y_i$, note that $\hat{\alpha}_{\hat{g}} = \hat{R}$, which implies that

$$\begin{aligned}\hat{\alpha}_{\text{corr}} &= \hat{\rho}\hat{\alpha}_{\hat{g}} \\ &= \frac{\hat{h}_{SNP}}{\hat{R}}\hat{R} \\ &= \hat{h}_{SNP}.\end{aligned}$$

So the true standard error is equal to the standard error of $\hat{h}_{SNP}$.

If $\hat{\rho}$ is calculated in a different dataset than the dataset used in the regression, the error in $\hat{h}_{SNP}$ and $\hat{R}$ will be uncorrelated with the error in $\hat{\alpha}_{\hat{g}}$. This means that the standard errors reported by our measurement-error software will be anti-conservative. However, since the error in $\hat{h}_{SNP}$ and $\hat{R}$ will be positively correlated, the sampling variance in $\hat{\rho}$ will likely be small, suggesting that the bias in the standard errors will also likely be small relative to the magnitude of the reported standard error.

### 6 Theoretical Framework with GWAS Controls

The theoretical framework in the main text is derived for PGI weights estimated in a GWAS conducted using ordinary least squares (OLS), without any control variables. In practice, PGI weights are virtually always derived from a GWAS that includes at least some basic set of control variables (typically sex, age, and at least four principal components (PCs) of the genotype data). We omit the covariates in the main text because doing so simplifies the exposition without altering any of the theoretical properties of the true additive SNP factor that we focus on in the main text. However, the choice of covariates is one of many dimensions of GWAS methodology that may matter in important ways in practical applications where a researcher is trying to interpret a PGI from a specific GWAS. To illustrate, we show below that the theoretical framework can be extended to account for the vector of control variables, $C$, included in the GWAS. The theoretical regression equation that defines the vector needs to be modified to include the control variables:

$$(\gamma^C, \kappa) = \arg \min_{(\tilde{\gamma}^C, \tilde{\kappa})} E[(y_i - x_i'\tilde{\gamma}^C - C_i'\tilde{\kappa})^2],$$

where we use the $C$ superscript to highlight the fact that the optimal weight vector with controls, $\gamma^c$, generally differs from the optimal weight vector without controls, which we denoted $\gamma$ in the main text. Although the additive SNP factor $g_i^C \equiv x_i'\gamma^C$ is in general different from $g_i \equiv x_i'\gamma$, as it is derived from $\gamma^C$ rather than $\gamma$, everything proceeds from here onward like in the main text. Since $g_i^c$ is a best linear predictor, it can be understood as a standardized, noisy measure of an unobserved, latent variable, and the error-in-variables bias and the measurement-error-corrected estimator formulas remain the same, with coefficients from the conditional analyses replacing the univariate coefficients. Compared to the main text, the only difference is that $g_i^c$ is now the best linear predictor of the phenotype *conditional* on the controls.

### 7 Polygenic Index Repository User Guide

In this section, we lay out some of the interpretational issues that are likely to arise as researchers begin to use PGIs from the Repository, and we outline how we suggest thinking through those issues. The executive summary is as follows:

1. The methodologies used to conduct the GWAS and to construct the PGI weights jointly determine the additive SNP factor that is proxied for by the PGI.

2. These methodologies, together with the PGI phenotype, determine the relative importance of various potential confounds to a causal interpretation of PGI associations.

3. Whether and which confounds should be highlighted (or can be safely ignored) depends on the application.
4. While a multi-trait PGI generally has higher predictive power than its corresponding single-trait PGI, it is subject to additional potential confounds. This tradeoff should be evaluated when deciding whether to use a single-trait or multi-trait PGI.
5. Currently, the most feasible way to cleanly identify causal effects of a PGI is to conduct a within-family analysis (where the PGI is analyzed in a sibling sample, with sibling fixed effects). In the absence of clean identification of a causal effect, researchers should highlight the potential confounds to a causal interpretation.
6. In interpreting PGI associations (whether causal or not), it is important to keep in mind that genetic effects can operate through environmental mechanisms, and these mechanisms may be modifiable. For this reason, terminology such as "genetic endowment" should be avoided. Researchers should remind readers of the potential role of environmental mechanisms in explaining PGI associations.

The following subsections, numbered 1 through 6, provide more detail on the points above. In addition to attending to these interpretational issues, we urge users of the Repository to conduct power calculations prior to undertaking analyses; to pursue analyses only if they are adequately powered; and, when feasible, to preregister planned analyses (along with the power calculations).

We note that the GWAS from which the Repository PGIs are constructed were conducted in European-ancestry samples (where "European-ancestry" is operationalized differently depending on the study but almost always involves sample restrictions based on the genetic PCs; e.g., for our UKB GWAS, see the "UKB GWAS" subsection of Section I in Methods). Due to the limited portability of such GWAS results to other ancestries, for the PGIs released to participating datasets, the current version of the Repository is restricted to individuals of European ancestries, as defined by how their genetic PCs cluster together with those classified as having European ancestries in the 1000 Genomes Project (see the "Subject-level QC in Repository Cohorts" subsection of Section II in Methods).

#### 7.1 GWAS and PGI-Weight Methodologies and the Additive SNP Factor

In the Supplementary Methods section 6, we showed how the set of control variables used in a GWAS affects the additive SNP factor proxied for by a PGI. The choice of controls, however, is just one of many dimensions of GWAS methodology. A change to any of these dimensions is likely to result in a different additive SNP factor (with a different interpretation). For example, it is increasingly common for datasets to conduct association analyses using mixed-linear models[2,3] rather than OLS. Since mixed-linear models often produce estimates that are more robust to stratification, the additive SNP factor will be akin to that generated by an OLS-based GWAS with some additional controls for stratification. Knowledge of the methodology of the GWAS underlying a particular PGI is therefore often a necessary first step for understanding what additive SNP factor a specific PGI is proxying for. For example, the methodologies underlying the GWASs we conducted in UKB for the PGIs in the Repository are described in the "UKB GWAS" subsection of Section I in Methods. Information about association models in 23andMe GWAS can be found in Supplementary Table 6.

The PGI-weight methodology can matter, as well. For example, our Repository PGI weights are calculated from the GWAS results using the HapMap3 set of SNPs, which primarily captures common genetic variation. If PGI weights were instead calculated based on results from SNPs that capture a different mix of common and rare genetic variation, then the additive SNP factor corresponding to that PGI would have a different interpretation: it would be the best linear predictor based on that set of SNPs.

#### 7.2 Potential Confounds to a Causal Interpretation

It is increasingly understood that standard GWAS approaches with a limited set of controls – for example, sex, age, and up to 10 PCs, as in most of the GWAS underlying the Repository PGIs – generate PGIs that can be subject to a number of confounds to a causal interpretation[4–7]. For example, PGIs for educational attainment derive a substantial share of their overall predictive power from their positive association with rearing environment. In behavior-genetic parlance, this positive correlation arises due to the vertical transmission of the parental phenotypes (parents' phenotypes impact their children's phenotypes). In recent molecular-genetic research, this source of positive gene-environment correlation has been labelled "genetic nurture"[5]. This effect can be further exacerbated by assortative mating at the genetic level.

As another example, when the PCs are estimated in a small sample, they are often not very accurate proxies for ancestry. Failure to adequately control for genetic ancestry gives rise to "population stratification"[8]: because the PGI is correlated with ancestry, which in turn is correlated with ethnicity and regional background, it picks up cultural or environmental factors that are correlated with these factors. In many empirical applications, the goal is to estimate an association that is net of any such cultural and environmental confounds. In such cases, it may be possible to mitigate concerns that the underlying GWAS may have relied on inaccurate ancestry controls by including a richer-than-usual set of environmental controls in the analysis of the PGI (i.e., in the vector $\mathbf{z}_i$ in equations (1) and (2) in the main text).[9].

The relevance of potential confounds could vary across phenotypes[4,6,7]. For example, genetic nurture effects are much smaller for height than educational attainment. Although the noisiness of PCs as measures of ancestry in a given sample is the same across phenotypes, the noisiness is likely to be substantially more problematic for educational attainment than for height because finer ancestral distinctions (which require more PCs to capture) probably matter for the social and environmental factors that influence educational attainment. More generally, it seems likely that potential confounds to a causal interpretation matter more for PGIs for social and behavioral phenotypes than for PGIs for more biologically proximal phenotypes.

#### 7.3 Importance of Confounds Depends On the Application

The degree to which potential confounds to a causal interpretation matter depends on how the PGI is used. For example, if a PGI is used as a control variable to increase precision for a randomized treatment evaluation[10,11], then the goal is simply to use controls that absorb as much residual variance as possible (and avoid controlling for any variables realized after the randomized intervention). Since the PGI is simply being used as a predictive variable, its interpretation is irrelevant in that case. As a contrasting example, consider the illustrative application in the main text that tests how much parental education mediates the predictive power of the PGI for educational attainment. There, the PGI should be understood as capturing some of the genetic nurture effects and ancestry associations with education. In most applications, the potential confounds do matter and should be highlighted.

#### 7.4 Single- Versus Multi-Trait PGIs

MTAG coefficient estimates are a weighted sum of GWAS coefficient estimates. Relative to GWAS estimates, MTAG coefficients have a lower expected mean-squared error, which means that multi-trait PGIs will in general have greater predictive power.

Multi-trait PGIs, however, do not necessarily have the same interpretation as single-trait PGIs. Because MTAG estimates are a weighted average of GWAS estimates for several traits, the multi-trait PGI based on MTAG estimates is roughly a weighted average of PGIs for the set of included traits. As a result, a multi-trait PGI may be correlated with an outcome variable if that outcome variable is genetically correlated with a supplementary phenotype for the multi-trait PGI. This can even be the case if the outcome variable and the target phenotype are not genetically correlated.

Therefore, **results using the multi-trait PGI have the same interpretation as results using the single-trait PGI in analyses where**

(i) the dependent variable and the PGI correspond to the same phenotype, *and*

(ii) no other covariates are included in the regression that are genetically correlated with any of the supplementary phenotypes used to construct the multi-trait PGI.

However, **results from the multi-trait PGI should be interpreted differently than results from the single-trait PGI—perhaps being driven by a supplementary phenotype rather than the target phenotype—if either (i) or (ii) is violated**. In that case, the risk of spurious results increases when (a) the GWAS sample size for the target GWAS is small relative to the GWAS sample size of the supplementary phenotypes, and (b) the genetic correlation between the target phenotype and the supplementary phenotypes is only moderate. Researchers who use multi-trait PGIs should make clear to readers how large the potential for a confounded interpretation is and how much it matters for the application at hand.

As described in Section 3 above, in settings where the PGI is just being used as a covariate (e.g., as a control variable in a randomized controlled trial), the confounds associated with using the multi-trait PGI may be less important. In all settings, however, it is good practice to describe which supplementary phenotypes were included in the multi-trait PGI if they are included in an analysis.

#### 7.5 Identifying Causal Effects of a PGI

A clean way to identify the causal effects of a PGI is to conduct the analysis in a sibling sample and control for family fixed effects. This strategy exploits a natural experiment: conditional on a pair of biological parents, genetic inheritance is random. A robustly estimated non-zero within-family association from a large and attrition-free sample would provide strong evidence of a causal effect of the PGI. The coefficient estimate could be interpreted as a weighted average of treatment effects from hypothetical experiments that randomly modify PGIs at conception[12,13].

The additive SNP factors corresponding to the PGIs in the Repository are not the best linear predictors conditional on a pair of biological parents (because the GWAS underlying the PGI weights do not control for the biological parents). The PGIs proxying for additive SNP factors that would be the best linear predictors for such a "within-family analysis" would be PGIs constructed from GWAS that control for parental genotypes or from GWAS (in sibling samples) that control for family fixed effects. Unfortunately, to date genotyped family-based samples have been too small to produce reliable "within-family PGIs." The Repository does not yet contain any such PGIs. Ultimately, however, when genotyped family-based samples become sufficiently large, the resulting within-family PGIs will be more predictive for within-family analyses than PGIs constructed from currently-standard (between-family) GWAS.

#### 7.6 Genetic Effects Can Operate Through Environmental Mechanisms

We encourage researchers who use PGIs in their research to be mindful of three important issues of interpretation for the causal effects of a PGI. First, a PGI could exert its effects through the environment[14]. Consider a PGI for BMI[10]. Suppose a within-family association analysis yields unambiguous evidence of a within-family association between the PGI and BMI. Even though the within-family design provides strong support for a causal interpretation, this does *not* imply that the SNPs in the PGI must be influencing BMI through some narrowly physiological mechanism. In principle, the sibling differences in BMI could arise because of sibling differences in genes that influence the proneness to eat sweets, exercise habits, or myriads of other behaviors with downstream effects on BMI. PGIs for seemingly "biological" phenotypes can thus have a substantial behavioral component. A PGI for lung health may similarly derive predictive power from SNPs that influence lung health very indirectly, through smoking habits[15,16].

Second and relatedly, it is therefore a fallacy to assume that any genetic sources of heterogeneity captured by a PGI are immutable, or at least harder to modify than environmental sources of heterogeneity. Indeed, the possibility of identifying modifiable mechanisms through which PGIs exert some of their effects motivates some of the research using PGIs[17,18]. To continue the BMI example, the widespread replacement of sugar by low-calorie sweeteners or better behavioral tools for avoiding temptation could eliminate or reduce the effect of the PGI on BMI. Because of these issues, we urge caution in describing PGIs as "genetic endowments," or related terminology that may, however inadvertently, promote the common misunderstanding that genes are a resource that is easily separable from choices made in light of that resource.

Third, because the additive genetic factor depends on the environment, the PGI may be context dependent. That is, the same PGI may have a different predictive power in two different samples if there are differences in the population sampled, the sampling methodology, or the phenotype measure. For example, the research participants from the UKB were recruited through the mail and had a 5.5% response rate. Those that responded to the recruitment mailers were more healthy and more educated than the UK population as a whole[19,20]. Because UKB participants make up a large fraction of the discovery sample for many phenotypes, it may be that the PGI from this Repository does not correspond to a PGI that would be produced from a representative sample or a sample of individuals not from the UK.
