## Supplementary Note for "Resource Profile and User Guide of the Polygenic Index Repository"

January 15, 2021

#### Contents


**1 Data Access Procedures** **2**

**2 Dataset Profiles** **4**

**3 Dataset-Specific Acknowledgments** **5**

**4 Dataset Authorship Contributions** **7**

### 1 Data Access Procedures

#### 23andMe

Upon publication of this paper, investigators at non-profit institutions can obtain access to the genome-wide summary statistics from 23andMe used in this paper by completing the 23andMe Publication Dataset Access Request Form. The information provided on this form will be used to generate a Statement of Work (SOW) that will allow 23andMe to transfer data for use in the described research project. The SOW and a Data Transfer Agreement will need to be signed by the institution and 23andMe before data can be shared. The form, as well as additional information and requirements, are available at https://research.23andme.com/dataset-access/.

#### Add Health

Access to the polygenic indexes and full phenotype data in Add Health is publicly available via a restricted data use contract with the University of North Carolina at Chapel Hill. Obtain access by visiting the CPC Data Portal at data.cpc.unc.edu/projects/2/view or see the Add Health contracts page at www.cpc.unc.edu/projects/addhealth/contracts. Add Health genotype data can be accessed via the database of Genotypes and Phenotypes (dbGaP, www.ncbi.nlm.nih.gov/gap, accession number phs001367.v1.p1).

#### Dunedin Multidisciplinary Health and Development Study

The datasets reported in the current article are available on request by qualified scientists. Requests require a concept paper describing the purpose of data access, ethical approval at the applicant's university, and provision for secure data access. We offer secure access on the Duke, Otago and King's College campuses. All data analysis scripts and results files are available for review. For more information, see moffittcaspi.trinity.duke.edu/research-topics/dunedin.

#### ELSA

Polygenic indexes and genotype data are publicly available and are available here: https://www.elsa-project.ac.uk/genetics. Phenotype and other publicly available data can be downloaded from the UK Data Service: https://beta.ukdataservice.ac.uk/datacatalogue/studies/study?id=5050. Use is limited to non-profit research use only. For more information regarding the data please contact.

#### E-Risk

The datasets reported in the current article are available on request by qualified scientists. Requests require a concept paper describing the purpose of data access, ethical approval at the applicant's university, and provision for secure data access. We offer secure access on the Duke and King's College campuses. All data analysis scripts and results files are available for review. For more information, see moffittcaspi.trinity.duke.edu/research-topics/erisk.

#### EGCUT

Estonian Biobank data is available for academic research. To request phenotype, polygenic index, and/or genotype data, researchers need to fill out a preliminary request form (available at genomics.ut.ee/en/biobank.ee/data-access) and submit it via e-mail to. The preliminary request will be evaluated by the Estonian Committee on Bioethics and Human Research. Upon positive review, researchers need to fill out a request form (also available at genomics.ut.ee/en/biobank.ee/data-access) and submit it via e-mail to. The data will then be shared pursuant to a Data Use Agreement. For further details, see genomics.ut.ee/en/biobank.ee/data-access.

#### HRS

Polygenic scores are publicly available and can be downloaded here: hrs.isr.umich.edu/data-products/genetic-data. Phenotype and other publicly available data can be downloaded here: hrs.isr.umich.edu/data-products. Genotype data can be accessed via the database of Genotypes and Phenotypes (dbGaP, www.ncbi.nlm.nih.gov/gap, accession number phs000428.v1.p1 and phs000428.v2.p2) with the most recent version forthcoming via NIAGADS (www.niagads.org/). Use is limited to non-profit research use only.

#### MCTFR

Access to the MCTFR PGIs is available by contacting Matt McGue, who will provide access authorization. Access to MCTFR phenotypic data will require a research proposal the structure of which can be provided by Matt McGue. Please note that the MCTFR is a complex, longitudinal study with thousands of relevant phenotypes assessed at multiple points in time. An overview of the range of phenotypes and developmental periods can be found in Wilson et al. (2019). Use of phenotypic data requires an approved proposal that is approved by the MCTFR Principal Investigator Committee; access to the MCTFR PGIs does not require an approved proposal. Because of the complexities involved, developing a proposal typically involves multiple iterations with MCTFR staff and are dealt with on a case-by-case basis.

#### STR

Researchers interested in using STR data must obtain approval from the Swedish Ethical Review Authority and from the Steering Committee of the Swedish Twin Registry. Researchers using STR data are required to follow the terms of a number of clauses designed to ensure protection of privacy and compliance with relevant laws. For further information please visit ki.se/en/research/the-swedish-twin-registry.

#### TTP

Access to the polygenic indexes and phenotype data from the Texas Twin Project is available via a restricted data use contract with the University of Texas at Austin. Restricted data users must develop an IRB-approved research proposal and security plan that ensures secure use of the data to minimize deductive disclosure risks. To apply for restricted-use data, please visit https://redcap.prc.utexas.edu/redcap/surveys/?s=FHJW9KCW8K.

#### UKB

All bona fide researchers can apply to use the UK Biobank resource for health related research that is in the public interest. Researchers can register and apply for data access at https://www.ukbiobank.ac.uk/register-apply/. Prior to publication of this paper, we will return the Repository PGIs to the UKB in accordance with their "returning results" procedure: https://biobank.ndph.ox.ac.uk/showcase/exinfo.cgi?src=returning_results. UKB will subsequently make the PGIs available to researchers as "Derived data-fields."

#### WLS

In addition to phenotype data, the polygenic index data is publicly available. As of February 2019, researchers who wish to use these polygenic indexes should email a brief research proposal and a copy or link to their CV to. Given the need to preserve participant confidentiality, to access the complete genotyped data, researchers will additionally need to receive IRB approval from their home institution and enter into a Data Use Agreement between the researcher's home institution and the University of Wisconsin-Madison. For the most up-do-date instructions, see www.ssc.wisc.edu/wlsresearch/documentation/GWAS/.

#### 2 Dataset Profiles

##### 23andMe

Eriksson, N. *et al.* Web-Based, Participant-Driven Studies Yield Novel Genetic Associations for Common Traits. *PLOS Genetics* 6(6), 1–20 (2010).

##### Add Health

Harris, K. M. *et al.* Cohort Profile: The National Longitudinal Study of Adolescent to Adult Health (Add Health). *International Journal of Epidemiology* 48(5), 1415–1425 (2019).

##### Dunedin Multidisciplinary Health and Development Study

Poulton, R, *et at.*. The Dunedin Multidisciplinary Health and Development Study: Overview of the first 40 years, with an eye to the future. *Social Psychiatry and Psychiatric Epidemiology* 50, 679–693 (2015).

##### ELSA

Steptoe, A. *et al.* "Cohort Profile: The English Longitudinal Study of Ageing (ELSA)." *International Journal of Epidemiology* 42(6), 1640–1648 (2013).

##### E-Risk

None.

##### EGCUT

Leitsalu, L. *et al.* Cohort Profile: Estonian Biobank of the Estonian Genome Center (EGCUT), *International Journal of Epidemiology* 44, 1137–1147 (2015).

##### HRS

Sonnega, A. *et al.* Cohort Profile: the Health and Retirement Study (HRS), *International Journal of Epidemiology* 43(2), 576–85 (2014).

##### MCTFR

Wilson, S. *et al.* Minnesota Center for Twin and Family Research (MCTFR). *Twin Research and Human Genetics* 22(6), 746–752 (2019).

##### STR

Zagai, U. *et al.* The Swedish Twin Registry (STR): Content and Management as a Research Infrastructure. *Twin Research and Human Genetics* 22(6), 672–680 (2019).

##### TTP

Harden, K.P. *et al.* The Texas Twin Project (TTP). *Twin Research and Human Genetics* 16(1), 385–90 (2013).

##### UKB

Sudlow, C *et al.* UK Biobank: An Open Access Resource for Identifying the Causes of a Wide Range of Complex Diseases of Middle and Old Age. *PLoS Med* 12(3) (2015).

#### WLS

Herd, P. *et al.* Cohort profile: Wisconsin Longitudinal Study (WLS). *International Journal of Epidemiology* 43, 34–41 (2014).

### 3 Dataset-Specific Acknowledgments

We gratefully acknowledge research participants from all cohorts.

#### 23andMe

We gratefully acknowledge the contributions of members of 23andMe's Research Team, whose names are listed below: Michelle Agee, Babak Alipanahi, Adam Auton, Robert K. Bell, Katarzyna, Bryc, Sarah L. Elson, Pierre Fontanillas, Nicholas A. Furlotte, Karen E. Huber, Nadia K. Litterman, Jennifer C. McCreight, Matthew H. McIntyre, Joanna L. Mountain, Carrie A.M. Northover, Steven J. Pitts, J. Fah Sathirapongsasuti, Olga V. Sazonova, Janie F. Shelton, Suyash Shringarpure, Chao Tian, Joyce Y. Tung, Vladimir Vacic, and Catherine H. Wilson.

#### Add Health

The National Longitudinal Study of Adolescent to Adult Health (Add Health) is supported by grant P01 HD031921 to Kathleen Mullan Harris from the Eunice Kennedy Shriver National Institute of Child Health and Human Development (NICHD), with cooperative funding from 23 other federal agencies and foundations. Add Health GWAS data were funded by NICHD grants to Harris (R01 HD073342) and to Harris, Boardman, and McQueen (R01 HD060726). For information about access to the data from this study, contact.

#### Dunedin Multidisciplinary Health and Development Study

Dunedin Multidisciplinary Health and Development Study research is supported by National Institute on Aging grants R01AG032282, R01AG049789, UK Medical Research Council grant MR/P005918, the New Zealand Health Research Council and New Zealand Ministry of Business, Innovation, and Employment.

#### ELSA

The English Longitudinal Study of Ageing is jointly run by University College London, Institute for Fiscal Studies, University of Manchester and National Centre for Social Research. Genetic analyses have been carried out by UCL Genomics and funded by the Economic and Social Research Council (ES/K005774/1) and the National Institute on Aging (R01 AG017644). All GWAS data has been deposited in the European Genome-phenome Archive. For more information please refer to www.elsa-project.ac.uk/genetics, or contact.

#### E-Risk

The E-Risk study is funded by grant G1002190 from the UK Medical Research Council and grant HD077482 from the National Institute of Child Health and Development.

#### EGCUT

EGCUT received funding from the Estonian Research Council Grant PUT1660 and PRG184, Mobilitas Pluss ERA-NET grant SP1GI18045T, Horizon 2020 program grants MMVCM18418R, and European Union through the European Regional Development Fund SLTMR16142T. For more information, please contact Tõnu Esko.

#### MCTFR

This project was led by William G. Iacono, PhD. and Matt McGue, PhD (Co-Principal Investigators) at the University of Minnesota, Minneapolis, MN, USA. Co-investigators from the same institution included: Irene J. Elkins, Margaret A. Keyes, James J. Lee, Lisa N. Legrand, Stephen M. Malone, William S. Oetting, Michael B. Miller, Saonli Basu and Scott Vrieze. Funding support for this project was provided through NIDA (U01DA024417). Other support for sample ascertainment and data collection came from several grants: R37DA05147, R01AA09367, R01AA11886, R01DA13240, R01MH66140.

#### STR

The Swedish Twin Registry (STR) is managed by Karolinska Institutet and receives additional funding through the Swedish Research Council under the grant no 2017-00641. Other funding for the project come from the Ragnar Söderberg Foundation (E9/11), the Swedish Research Council (421-2013-1061).

#### TTP

The Texas Twin Project is supported by grants R01HD083613 and R01HD092548 from NIH/NICHD and Jacobs Foundation Research Fellowships.

#### WLS

This research uses data from the Wisconsin Longitudinal Study (WLS) of the University of Wisconsin-Madison. Since 1991, the WLS has been supported principally by the National Institute on Aging (AG-9775, AG-21079, AG-033285, and AG-041868, R01 AG041868-01A1), with additional support from the Vilas Estate Trust, the National Science Foundation, the Spencer Foundation, and the Graduate School of the University of Wisconsin-Madison. Since 1992, data have been collected by the University of Wisconsin Survey Center. The opinions expressed herein are those of the authors. A public use file of data from the Wisconsin Longitudinal Study is available from the Wisconsin Longitudinal Study, University of Wisconsin-Madison, 1180 Observatory Drive, Madison, Wisconsin 53706 and at www.ssc.wisc.edu/WLSresearch/data/.

### 4 Dataset Authorship Contributions

| Dataset | Author | Study design & mgmt. | Data collection | Genotyping | Genotype prep. | Phenotype prep. | Data analysis | Writing |
|---|---|---|---|---|---|---|---|---|
| **23andMe** | Aaron Kleinman | | | | | X | X | X |
| **23andMe** | David A. Hinds | X | | | | X | | X |
| **23andMe** | 23andMe Research Group | | X | X | | X | X | |
| **Add Health** | Kathleen Mullan Harris | X | X | X | X | X | | |
| **Dunedin Study** | Daniel W. Belsky | | | | | X | X | X |
| **Dunedin Study** | Avshalom Caspi | X | X | | | | X | X |
| **Dunedin Study** | David L. Corcoran | | | X | X | | X | |
| **Dunedin Study** | Terrie E. Moffitt | X | X | | | X | | X |
| **Dunedin Study** | Richie Poulton | X | X | | | X | | X |
| **Dunedin Study** | Karen Sugden | | | X | X | X | X | X |
| **Dunedin Study** | Benjamin S. Williams | | | X | X | X | | |
| **ELSA** | Andrew Steptoe | X | X | | | | | |
| **ELSA** | Olesya Ajnakina | | | | X | | | |
| **E-Risk** | Daniel W. Belsky | | | | | X | X | X |
| **E-Risk** | Avshalom Caspi | X | X | | | | X | X |
| **E-Risk** | David L. Corcoran | | | X | X | | X | |
| **E-Risk** | Terrie E. Moffitt | X | X | | | X | | X |
| **E-Risk** | Karen Sugden | | | X | X | X | X | X |
| **E-Risk** | Benjamin S. Williams | | | X | X | X | | |
| **EGCUT** | Lili Milani | X | X | X | X | X | X | |
| **EGCUT** | Tõnu Esko | X | X | X | X | X | X | |
| **MCTFR** | William G. Iacono | X | X | | | | | |
| **MCTFR** | Matt McGue | X | X | | | X | | |
| **STR** | Rafael Ahlskog | X | | | | | | |
| **STR** | Patrik K.E. Magnusson | X | X | | | X | | |
| **TTP** | Travis T. Mallard | | | | X | | | |
| **TTP** | K. Paige Harden | X | | | | | | |
| **TTP** | Elliot M. Tucker-Drob | X | | | | | | |
| **WLS** | Pamela Herd | X | X | X | | | | |
| **WLS** | Jeremy Freese | X | X | X | | | | |
