## Supplementary material for "Resource Profile and User Guide of the Polygenic Index Repository": FAQ

**Frequently Asked Questions (FAQs)**

This document provides information about the study:

Becker *et al.* (2021) "Resource Profile and User Guide of the Polygenic Index Repository" *Nature Human Behaviour*, in press.

The document was prepared by Daniel Benjamin, David Laibson, Michelle N. Meyer, and Patrick Turley. It draws from and builds on the FAQs for earlier SSGAC papers. It has the following sections:

1. **Background**
2. **Study design and results**
3. **Social and ethical implications of the study**
4. **Appendices**

For clarifications or additional questions, please contact Daniel Benjamin.

### 1. Background

#### 1.1. Who conducted this study? What are the group's overarching goals?

The authors of the study are researchers affiliated with the Social Science Genetic Association Consortium (SSGAC) as well as data providers (i.e., individuals who act as stewards for datasets and provide other researchers with access to these data for research purposes). The SSGAC is a multi-institutional, international research group that aims to identify statistically robust associations between variation in DNA and variation in social-science-relevant outcomes.

We study the most common sources of genetic variation—single-nucleotide polymorphisms (SNPs). SNPs are sites in the genome where single DNA base pairs commonly differ across individuals. Each SNP usually has two different possible base pairs, which are called alleles. Although there are tens of millions of sites where SNPs are located in the human genome, our work (like most genetic research today that aims to link variation in DNA to variation in disease and other outcomes) investigates only SNPs that can be easily measured with a high level of accuracy. These days, we can easily and accurately measure millions of SNPs, which together capture most of the common genetic variation across people.

The social-science-relevant outcomes that we analyze include differences across people in behavior, preferences, and personality that are traditionally studied by social and behavioral scientists (e.g., anthropologists, economists, political scientists, psychologists, and sociologists). These traits are often also of interest to health and other researchers.

The SSGAC was formed in 2011 to address a specific set of scientific challenges. Most outcomes and behaviors are weakly associated with a very large number of SNPs. Although their collective effect can be meaningful (see FAQs 1.2 & 2.3), we now know that almost every one of these SNPs has an extremely weak association on its own. To identify specific SNPs with such small effects, scientists must study at least hundreds of thousands of people (to separate weak signals from sampling noise). One promising strategy for doing this is for many investigators to pool their data into one large study. This approach has borne considerable fruit when used by medical geneticists interested in a range of medical conditions (Visscher et al., 2017). Most of these advances would not have been possible without large research collaborations between multiple research groups interested in similar questions. The SSGAC was formed in an attempt by social scientists to adopt this research model.

The SSGAC is organized as a working group of the Cohorts for Heart and Aging Research in Genomic Epidemiology (CHARGE), a successful medical consortium. (In genetics research, "cohort" is a term that means "dataset.") The SSGAC was founded by three social scientists—Daniel Benjamin (University of California – Los Angeles), David Cesarini (New York University), and Philipp Koellinger (University of Wisconsin and Vrije Universiteit Amsterdam)—who believe that studying SNPs associated with social scientific outcomes can have substantial positive impacts across many research fields. This includes research that aims to better understand the effects of the environment (e.g., research on policy interventions) and interactions between genetic and environmental effects. The potential benefits also span a diverse set of research questions in the biomedical sciences, such as why and how educational attainment is linked to longevity and better overall health outcomes.

To conduct such research, the SSGAC implements genome-wide association studies (GWAS) of social-scientific outcomes. For example, to conduct a GWAS of educational attainment (e.g., Lee et al., 2018) every participating cohort calculates the cross-sectional (i.e., within-cohort) correlation between educational attainment and DNA-base-pair variation at a single location on the genome: a SNP. As first discussed above, a SNP is a base-pair of the genome where there is common variation in the human population. This statistical analysis is repeated for each SNP on the genome. The cohort-level results do not contain individual-level data—just summary statistics about these within-cohort statistical associations. The SSGAC then combines these cohort results to produce the overall GWAS results. By using existing datasets and combining cohort results, we can study the genetics of large numbers of individuals (for example, ~1.1 million people in Lee et al. (2018)) at very low cost. The SSGAC publicly shares overall, aggregated results (subject to some Terms of Service; see FAQ 3.7) so that other scientists can build on this work. These publicly available data have already catalyzed many research projects and analyses across the social and biomedical sciences. Among the most useful products of these GWASs for other research are the polygenic indexes that are based on GWAS associations. Polygenic indexes are variables that aggregate the predictive power of many SNPs for predicting the outcome of the GWAS (see FAQ 1.2), and they are the focus on the current paper.

The Advisory Board for the SSGAC is composed of prominent researchers representing various disciplines: Dalton Conley (Sociology, Princeton University), George Davey Smith (Epidemiology, University of Bristol), Tõnu Esko (Molecular Biology and Human Genetics, University of Tartu and Estonian Genome Center), Albert Hofman (Epidemiology, Harvard University), Robert Krueger (Psychology, University of Minnesota), David Laibson (Economics, Harvard University), James Lee (Psychology, University of Minnesota), Sarah Medland (Genetic Epidemiology, QIMR Berghofer Medical Research Institute), Michelle Meyer (Bioethics and Law, Geisinger Health System), and Peter Visscher (Statistical Genetics, University of Queensland).

The SSGAC is committed to the principles of reproducibility and transparency. Major SSGAC publications are usually accompanied by a FAQ document (such as this one). The FAQ document is written to communicate what was found less tersely and technically than in the paper, as well as what can and cannot be concluded from the research findings more broadly. FAQ documents produced for SSGAC publications are available on the SSGAC website.

To date, SSGAC-affiliated papers have studied educational attainment, cognitive performance, subjective well-being, reproductive behavior, risk tolerance, and dietary intake. The SSGAC website contains a list of our major publications, which have been published in journals such as *Science*, *Nature*, *Nature Genetics*, *Proceedings of the National Academy of Sciences*, *Psychological Science*, and *Molecular Psychiatry*.

#### 1.2. What is a polygenic index (PGI)? Why this terminology?

A polygenic index (we use the acronym PGI throughout the paper) is an index composed of a large number of SNPs from across the genome. Each polygenic index is associated with a particular outcome (for details, see FAQ 1.3). Because a polygenic index aggregates the information from many SNPs, it can "predict" (see FAQ 1.6) far more of the variation among individuals than any single SNP. (Note that even polygenic indexes are not good predictors of outcomes for one person; see FAQ 3.3). Often, the polygenic indexes with the most predictive power are those created using *all* the (millions of) SNPs measured in a SNP array. A SNP array is the currently standard way of measuring common genetic differences across individuals. A SNP array data does not measure the entire genetic sequence of each individual, but it does measure most of the places on the genome where individuals differ.

Our terminology of *polygenic index* is currently non-standard, but most of the authors of the paper prefer it to current terms and hope that this paper, and the Polygenic Index Repository introduced in this paper, make polygenic index a standard term. The traditional terms include polygenic risk score and polygenic score. The word *risk* makes little sense when the polygenic index is for a non-disease outcome (such as height). The word *score* was intended to echo statistical nomenclature but can instead convey an unintended value judgment or valence (i.e., "a higher score must be better"). The word *index* is at least as accurate statistically and does not convey a value judgment.

#### 1.3. How is a polygenic index constructed?

A polygenic index is constructed in three steps. First, a genome-wide association study (GWAS) is conducted, looking at SNPs measured across the entire human genome to see which of them are associated with higher or lower levels of some outcome. As explained above, SNPs are sites in the genome where single DNA base pairs commonly differ across individuals. SNPs usually have two different possible base pairs, or alleles. Although there are tens of millions of sites where SNPs are located in the human genome, GWASs typically investigate only SNPs that can be easily measured (or imputed) with a high level of accuracy. These days, we can easily and accurately measure millions of SNPs, which together capture most of the common genetic variation across people. For each of these millions of SNPs, the GWAS generates an "effect size" corresponding to the (typically miniscule) magnitude of the association between that SNP and the outcome. (We use the term "effect size" because it is a common scientific shorthand for "magnitude of association," but we emphasize that use of the term is not intended to imply that the SNP, or polygenic index, *causes* the outcome; see FAQ 1.5.)

Second, the effect sizes are used to determine the "weight" each SNP will get in the polygenic index. The simplest scheme is to weight each SNP by its effect size as estimated in the GWAS. This simple weighting scheme has one main problem: because SNPs tend to be correlated with nearby SNPs on the genome (a phenomenon called linkage disequilibrium), if one SNP is associated with the outcome, nearby SNPs will also be associated with the outcome. State-of-the-art approaches to determining the weights for a polygenic index are designed to address this problem. We use a common approach called LDpred (Vilhjálmsson et al., 2015). Using the results of a GWAS, LDpred generates a weight for each SNP. These weights are not equal to the SNPs' effect sizes as estimated in the GWAS, mostly because the weights take into account each SNP's correlation with other SNPs. (Even though LDpred addresses the issue of linkage disequilibrium, it does so only for the purpose of generating weights for optimal prediction. LDpred will not necessarily assign more weight to the SNP whose association with the outcome is responsible for nearby SNPs' associations with the outcome. Thus, LDpred is a tool to address the issue of linkage disequilibrium for the purpose of prediction—which is the purpose of a polygenic index—but not for the purpose of unbiased estimation of SNPs' effect sizes. See FAQ 1.5.)

Third, the set of weights for the SNPs are used in a formula for calculating a polygenic index for any particular individual. The formula is a weighted sum of alleles at each SNP (using the weights from the second step). The formula is used to calculate a numerical value of the polygenic index for each individual in some dataset (that was not included in the GWAS).

The sample used for the GWAS in the first step is the training sample for the polygenic index. The larger the GWAS sample size, the greater the predictive power of a polygenic index constructed in the third step. However, this predictive power of a polygenic index has a maximum for each outcome that the polygenic index can approach as the sample size gets bigger, but it can never exceed.

#### 1.4. How might polygenic indexes be useful?

A polygenic index for an outcome provides one measure of the genetic influence on that outcome that can be used in research in a variety of ways. For example, polygenic indexes have been used to:

- partially control for genetic influences in order to generate less noisy estimates of how changes in school policy influence health outcomes (Davies et al., 2018);
- examine how the effect of school policy on health outcomes depends in part on genetic influences (Barcellos, Carvalho and Turley, 2018a);
- study why SNPs predict educational attainment – for example, it appears that some genetic effects on educational attainment operate through associations with cognitive function and traits such as self-control (Belsky et al., 2016), which in turn affect educational attainment;
- investigate how genetic influences on educational attainment differ across environmental contexts (Schmitz and Conley, 2017; Barcellos, Carvalho and Turley, 2018b);
- investigate how genetic influences on BMI vary over the lifecycle (Khera et al., 2019);
- infer the degree of assortative mating (Robinson et al., 2017; Yengo et al., 2018);
- trace recent migration patterns (Domingue et al., 2018; Abdellaoui et al., 2019);
- examine whether polygenic indexes for disease risk are sufficiently predictive to be incorporated into clinical practice for preventative medicine (Khera et al., 2018); and
- develop new statistical tools that may advance our understanding of how parenting and other features of a child's rearing environment influence his or her developmental outcomes (Koellinger and Harden, 2018; Kong et al., 2018).

The idea of using GWAS results to create a polygenic index was initially proposed in 2007 (Wray, Goddard and Visscher, 2007), and the first polygenic index was created in 2009 in a GWAS of schizophrenia and bipolar disorder (Purcell et al., 2009). Since then, polygenic indexes have become a significant part of research that builds on genetics in the medical and social sciences. For example, in the current paper we analyze presentations at the annual meeting of the Behavior Genetics Association. We report that the fraction of presentations that used polygenic indexes increased from 0% in 2009 to 20% in 2019. The list above represents a few illustrative examples of research that uses polygenic indexes.

As discussed in FAQ 1.9 below, one goal of this paper, and the Polygenic Index Repository it introduces, is to facilitate further work using polygenic indexes by making a much wider range of more predictive polygenic indexes available to researchers.

#### 1.5. Does a polygenic index "cause" the outcome of interest?

Polygenic indexes available today, including those we construct in this paper, should not be interpreted as a measure of causal mechanisms.

The genome-wide association studies (GWASs) used as the training data for the polygenic indexes (see FAQ 1.3) identify SNPs that are associated with the outcome, but an observed empirical correlation with a specific SNP need not imply that the SNP *causes* the outcome, for a variety of reasons. First, SNPs are often highly correlated with other, nearby SNPs on the same chromosome. As a result, when one or more SNPs in a region causally influence an outcome (in that particular environment), many non-causal SNPs in that region may also be identified as associated with the outcome (in FAQ 1.3, see the parenthetical "Even though LDpred…" for why LDpred does not solve this problem for the purpose of identifying the causal SNP). In fact, the causal SNP may not have even been measured directly. For example, GWAS that focus on common SNPs would not be able to identify rare or structural types of genetic variation (e.g., deletions or insertions of an entire genetic region) that are causal, but they may identify SNPs that are correlated with these unobserved variants. For these and other reasons, polygenic indexes are likely to be composed of a mix of causal and non-causal SNPs, and the weights used in the formula for constructing the polygenic index (see FAQ 1.3) should not be interpreted as estimates of the causal effects of the SNPs. As a very rough estimate, for social and behavioral outcomes, no more than about one-third of the predictive power of a polygenic index (i.e., the percentage of the variance in the outcome among individuals that the polygenic index explains) is explained by causal genetic effects (Howe et al., 2021). For instance, the most predictive polygenic index for educational attainment currently available explains about 12% of the variance between people, but only one-third of that—about 4%—is causal. (These causal SNPs may be among the SNPs included in the polygenic index or may be physically close to, and therefore correlated with, SNPs that are included.) In contrast, for anthropometric outcomes such as height, it is possible that nearly all of the predictive power of a polygenic index is explained by causal SNPs.

Second, at a particular SNP the frequency of different alleles might vary systematically across environments. If those environmental factors are not accounted for in the association analyses, some of the measured SNP associations with social-science outcomes may be spurious. To use a well-known example often used to explain this idea (Lander and Schork, 1994), any genetic variants common in people of Asian ancestries will be associated statistically with more frequent than average chopstick use, but these variants would not *cause* greater chopstick use; rather, these genetic variants and the outcome of chopstick use are both distributed unevenly among people with different ancestries. This is called the problem of "population stratification." The GWAS underlying the polygenic indexes in this paper employ standard strategies to try to minimize this problem, but the issues raised by population stratification cannot be ruled out entirely. As a result, the polygenic indexes likely reflect population stratification to some extent. In the User Guide that accompanies the Polygenic Index Repository (reproduced in the Supplementary Methods of the paper), we discuss this problem in more detail and discuss strategies for addressing the population stratification in the polygenic indexes

Even in GWAS (such as those we rely on or conduct ourselves) that attempt to address and correct for heterogeneity in genetic ancestry, allele frequencies may nonetheless vary systematically with environmental factors even *within* a group of people of similar genetic ancestry. For example, a SNP that is associated with improved educational outcomes in the parental generation may have downstream effects on parental income and other factors known to influence children's educational outcomes (such as neighborhood characteristics). This same SNP is likely to be inherited by the children of these parents, creating a correlation between the presence of the SNP in a child's genome and the extent to which the child was reared in an environment with specific characteristics. A recent study of Icelandic families showed that a parental allele associated with higher educational attainment of the parent that is *not* passed on to the parent's offspring is still associated with the child's educational attainment, suggesting that GWAS results for educational attainment partly represent these intergenerational environmental pathways (Kong et al., 2018).

Third, a SNP's effects on an outcome may be indirect, so a SNP that may be "causal" in one environment may have a diminished effect or no effect at all in other environments. For example, variation in a particular SNP on chromosome 15 is associated with lung cancer (Amos et al., 2008; Hung et al., 2008; Thorgeirsson et al., 2008). From this observation alone we cannot conclude that variation in this SNP can cause lung cancer through some direct *biological* mechanism. In fact, it is likely that variation in this SNP, which is part of the nicotinic acetylcholine receptor gene cluster that affects nicotine metabolism, increases lung cancer risk through effects on smoking behavior. In a tobacco-free environment, it is plausible that this association with lung cancer would be substantially weaker and perhaps disappear altogether. Thus, even *if* we have credible evidence that a specific association is not spurious, it is entirely possible that the SNP in question influences the outcome through channels that we, in common parlance, would label environmental (e.g., smoking). Nearly forty years ago, the sociologist Christopher Jencks criticized the widespread tendency to mistakenly treat environmental and genetic sources of variation as mutually exclusive (see also Turkheimer, 2000). As the example of smoking illustrates, it is often overly simplistic to assume that "genetic explanations of behavior are likely to be exclusively physical explanations while environmental explanations are likely to be social" (Jencks, 1980, p723).

#### 1.6. In what sense does a polygenic index "predict" the outcome of interest?

When we and other scientists say that polygenic indexes (and other variables, such as demographics or other environmental factors) "predict" certain outcomes, our use of the word differs in several important ways from how "predict" is used in standard language (e.g., outside of social science research papers). First, we do not mean that the polygenic index guarantees an outcome with 100% probability, or even with a high degree of likelihood. Rather, we mean that the polygenic index is, on average across people, statistically associated with an outcome. In other words, on average, people with a higher numerical value of the polygenic index have a higher likelihood of the outcome compared to people with a lower numerical value. A polygenic index is said to be statistically "predictive" of an outcome even if the polygenic index has only a *weak* association with the outcome—as is the case, for instance, with almost all of the polygenic indexes in this paper. In such cases, the polygenic index is only weakly predictive of the outcome.

Second, in standard language, "prediction" usually refers to the future. In contrast, when scientists say that a polygenic index "predicts" an outcome, they mean that they expect to see the association in *new data*. "New data" means data that haven't been analyzed yet—regardless of whether those data will be collected in the future or have already been collected. In other words, in social science, it makes perfect sense to ask how well a polygenic index predicts outcomes that have already occurred, like how many years of education were attained by older adults.

Finally, in standard language, a "prediction" is often an unconditional guess about what will happen. Instead of meaning it unconditionally, scientists mean that they expect to see an association in new data under certain conditions, for example, that the environment for the new data is the same as the environment in which the GWAS that underlies the polygenic index (see FAQ 1.3) was conducted. In the example given in FAQ 1.5, in which a SNP is associated with lung cancer due to an effect on smoking, we would *not* expect the SNP to be as strongly predictive of lung cancer, or predictive at all, in an environment where tobacco-based products are hard to obtain or absent entirely.

#### 1.7. What polygenic indexes were available to researchers prior to this project?

Prior to this project, only a few datasets had constructed polygenic indexes that researchers could download and use. Notable examples of data providers that did make polygenic indexes directly available to researchers —all of which recognized early on the value of doing so—are the Health and Retirement Study, the Wisconsin Longitudinal Study, and the National Longitudinal Adolescent to Adult Health Study. The UK Biobank does not construct polygenic indexes for its users, but it provides a mechanism by which researchers who use the data and construct polygenic indexes can "return" them to the UK Biobank for use by other researchers. Through this mechanism, polygenic indexes constructed from several GWASs have been made available for researchers to download from the UK Biobank.

To study polygenic indexes in other datasets or for other outcomes, prior to this paper, researchers would need to construct the polygenic indexes themselves, following the steps described in FAQ 1.3. For the first step, most researchers would need to rely on publicly available GWAS results, which include less data and are therefore less predictive than some polygenic indexes in published work that rely on non-public GWAS results (see FAQ 2.3). Recently, to make it easier for researchers to construct polygenic indexes themselves, the Polygenic Score Catalog (Lambert *et al.*, 2020) collected together weights for a range of polygenic indexes (also based on publicly available GWAS results).

As we discuss in more detail in FAQ 2.1, for the Polygenic Index Repository, we constructed a large number of polygenic indexes in each of 11 datasets (including the four mentioned above) and have made the polygenic indexes directly available for researchers to download. The polygenic indexes are often based on more data than is publicly available, and the polygenic indexes are constructed according to a uniform methodology across both outcomes and datasets. For examples of Repository polygenic indexes that were previously not available at all or that were less accurate (i.e., predictive), see FAQ 2.3.

#### 1.8. How do different polygenic indexes for the same outcome differ? How comparable are results across studies that use different polygenic indexes for the same outcome?

There are several reasons why polygenic indexes for the same outcome can differ from each other. As described in FAQ 1.3, there are three steps to creating a polygenic index, and differences can arise at each of these steps. For example, in the first step, researchers could base the polygenic index on different GWAS studies of the same outcome. Different GWAS studies may be based on samples who live under different environmental conditions, may have different measures of the outcome, and/or may have measured different SNPs. As another example, in the second step, researchers could use a different method of determining polygenic-index weights from the results of a GWAS. For these and other reasons, it has been common for different studies to use different polygenic indexes, even when the polygenic indexes are for the same outcome and are being studied in the same dataset.

The results are typically difficult to compare across such studies for three main reasons:

1. If the polygenic indexes are constructed using different methods, then even though they are both measuring genetic influences on the outcome, the precise definition of these "genetic influences" may differ (see FAQs 3.1 and 3.2).
2. The units for measuring the strength of associations between the polygenic index and other variables generally differ across studies. Researchers usually report results in terms of standard deviations (a statistical unit) of the polygenic index, but if the polygenic index in one study is a more powerful predictor than that in the other study, then one standard deviation of one polygenic index means something different than one standard deviation of the other.
3. If one of the polygenic indexes is a more powerful predictor than the other, then they differ in their signal-to-noise ratio for capturing genetic influences on the outcome. Whenever an explanatory variable is measured with noise, results based on that variable will be distorted, sometimes in unanticipated ways. Since the signal-to-noise ratio differs across the polygenic indexes, results based on them are distorted differentially, further making the results difficult to compare.

##### 1.9. Why create the Polygenic Index Repository?

In brief, the Polygenic Index Repository introduced in this paper has three main goals: (i) to make polygenic indexes for a large number of outcomes more accessible to a wider range of researchers from many fields and disciplines, including early career researchers, researchers without access to the data and/or training required to create the most state-of-the-art polygenic indexes, and researchers who wish to probe the limitations of polygenic indexes; (ii) to increase the use of polygenic indexes that are more accurate (i.e., predictive) than polygenic indexes researchers could construct from publicly available GWAS results; and (iii) to facilitate the comparability of results across studies that use these polygenic indexes.

In more detail, the Polygenic Index Repository addresses several practical obstacles that researchers interested in using polygenic indexes must often confront, including:

1. Constructing a polygenic index from genotype data requires special expertise. Even for researchers with that expertise, it can be a time-consuming process.
2. It is generally desirable to generate polygenic-index weights from the GWAS with the largest sample size because the predictive accuracy of a polygenic index is expected to be largest in that case. However, there are administrative hurdles for accessing some GWAS results, such as those from 23andMe. In practice, researchers often end up constructing polygenic indexes using only publicly available GWAS results. Such polygenic indexes tend to have less predictive power.
3. Publicly available GWAS results are sometimes based on a sample that includes the dataset (or close relatives of dataset members) in which the researcher wants to analyze the polygenic index. Such "sample overlap" spuriously inflates the predictive power of the polygenic index, which can lead to highly misleading results.

4. Because different researchers construct polygenic indexes in different ways, it is hard to compare and interpret results from different studies (see FAQ 1.8)

As we explain in the paper:

> We overcome #1 by constructing the [polygenic indexes] ourselves and releasing them to the data providers, who in turn will make them available to researchers. This simultaneously addresses #2 because we use all the data available to us that may not be easily available to other researchers or to the data providers, including genome-wide summary statistics from 23andMe. Using these genome-wide summary statistics from 23andMe is what primarily distinguishes our Repository from existing efforts by data providers to construct PGIs and make them available…It also distinguishes our Repository from efforts to make publicly available [polygenic index] weights directly available for download (although we also do that, for weights constructed without 23andMe data). To deal with #3, for each [outcome] and each dataset, we construct a [polygenic index] from GWAS summary statistics that excludes that dataset. We overcome #4 by using a uniform methodology across the [outcomes].

In addition to providing polygenic indexes constructed using a uniform methodology (which deals with problem #1 listed in FAQ 1.8), we aim to improve comparability of results based on polygenic indexes in another way (which deals with problems #2 and #3 listed in FAQ 1.8): we derive a "measurement-error-corrected estimator" and provide software for calculating it. This estimator deals with the fact that polygenic indexes can differ from each other in their signal-to-noise ratios. It estimates what the results of an analysis would be if the polygenic index had no noise. It thereby avoids the distortions in results that arise from having a noisy measure. Because it puts results about the polygenic index in the units of the "noiseless" polygenic index, the results from polygenic indexes with different signal-to-noise ratios are expressed in the same units. For more details, see FAQ 2.4.

#### 2. Study Design and Results

##### 2.1. What outcomes are included in the Polygenic Index Repository? How did you choose the outcomes?

We constructed polygenic indexes for 47 outcomes in 11 datasets, using a consistent methodology. The outcomes (listed in Table 1 in the paper) can be categorized into five groups:

- anthropometric (height and body mass index);
- cognition and education (including number of years of formal schooling and performance on cognitive tests);
- fertility and sexual development (including number of children separately for men and women, and age at first menses);
- health and health behaviors (the largest category, which includes self-rated overall health, several alcohol and smoking-related behaviors, and depressive symptoms); and
- personality and well-being (the next largest category, which includes self-rated risk tolerance, subjective well-being, and adventurousness).

The set of 47 outcomes we studied was selected from a larger set of 53 outcomes; we did not create polygenic indexes for the 6 outcomes for which statistical calculations indicated that, based on the GWAS results we had available, a polygenic index was predicted to explain less than 1% of the variation across individuals. Although the specific threshold of 1% is somewhat arbitrary (but see further discussion in FAQ 2.3 below), polygenic indexes with low predictive power are less useful and more likely to generate misleading results (such as false positives) if used.

#### 2.2. How did you create these polygenic indexes?

In order to construct the polygenic indexes, we combined GWAS results from three sources. First, for the 34 outcomes where we could find previously published GWAS, we obtained the publicly available results. Second, we collaborated with the personal genomics company 23andMe. 23andMe contributes to academic research by analyzing the data of customers who consent to participate in research. For this paper, 23andMe provided GWAS results for 37 outcomes, 9 of which had not previously been published. Third, for 25 outcomes, we conducted a GWAS ourselves in the UK Biobank, a large-scale biomedical database accessible to researchers. When more than one of these sources of GWAS results was available for an outcome, we combined the GWAS results together using a statistical method called meta-analysis. In some cases, we constructed "multi-trait polygenic indexes" using GWAS results for multiple outcomes (Turley et al., 2018); these polygenic indexes are often more predictive than a standard "single-trait polygenic index" constructed from GWAS results from a single outcome (FAQ 1.3), but the results from analyzing multi-trait polygenic indexes are sometimes more difficult to interpret (FAQ 2.5).

#### 2.3. How predictive are the polygenic indexes in the Repository?

To assess the predictive power of the polygenic indexes, we used data from 5 of the 11 participating datasets (those for which we had access to both the outcome and genotype data we needed to construct the polygenic indexes). In each of these 5 datasets, we calculated the predictive power of every polygenic index for which the dataset contained data on the relevant outcome (see FAQ 2.1).

The predictive power of the polygenic indexes varies substantially across the outcomes and validation datasets. The polygenic index for height has the greatest predictive power. It predicts 26% to 34% of the variation across individuals, depending on the validation dataset. Next is the polygenic index for body mass index (BMI), whose predictive power ranges from 13% to 15% in our validation datasets. Several outcomes—cognitive performance, age at first menses, and educational attainment—have a polygenic index with predictive power in the range of 6% to 12%. Among the least predictive are the polygenic indexes for satisfaction with family and satisfaction with friendships, whose predictive powers in our validation datasets range from 0.3% to 0.7% (they were included because their predictive power was statistically expected to exceed 1%; see FAQ 2.1). The predictive powers for the other polygenic indexes in the Repository lie somewhere between 1% and 6%.

Although the effects explained by these polygenic indexes are small-to-modest, they can nevertheless be useful in research. For instance, the environmental factors studied in economics research typically have predictive power smaller than 5%, often 1% or smaller. Among the strongest predictors of educational attainment is family socioeconomic status, which has predictive power of roughly 15%. In a standard categorization used in psychology (Cohen, 1992; percentages here are squared *r* values) predictive power less than 9% is "small" while predictive power greater than 25% (rarely attained in psychological research) is "large." We caution, however, that these comparisons of the effect sizes of polygenic indexes and environmental influences aren't apples-to-apples because researchers usually study one particular environmental factor or many on an outcome, whereas a polygenic index summarizes the predictive power of SNPs across the genome. As discussed further in FAQ 3.3, for social and behavioral outcomes, the sum of all environmental (i.e., non-genetic) influences substantially outweigh the sum of all genetic influences that a polygenic index aims to capture.

As we discuss in FAQ 3.3, an individual's polygenic indexes (even for height) do *not* very accurately predict that individual's outcomes. However, polygenic indexes are useful for *scientific studies* (including social science, health research, etc.). Such studies are concerned with aggregate population trends and averages rather than with individual outcomes. For example, for a polygenic index that predicts 1% of the variation across individuals, studies of its association with other variables can be well powered in sample sizes as small as 785 individuals; 10 out of the 11 datasets participating in the Repository have sample sizes larger than that.

A major goal of the Polygenic Index Repository is to enable other research that is valuable to social scientists and health researchers. Such studies are already being conducted with some polygenic indexes (see FAQ 1.9). For some outcomes, the polygenic indexes in the Repository are more predictive than those that were previously possible to construct; examples include having asthma/eczema/rhinitis, number of cigarettes smoked per day, having migraines, nearsightedness, self-reported physical activity, self-rated overall health, extraversion (i.e., being outgoing), and subjective well-being (i.e., self-reported happiness or life satisfaction). For other outcomes, polygenic indexes were not available prior to this paper because there had been no large published GWASs for those outcomes; examples include childhood reading, self-rated math ability, and self-reported narcissism, and several allergies including to pollen.

#### 2.4. What is the "measurement-error-corrected estimator"? How will it and the Repository improve comparability of results across future studies?

To understand this tool, it's helpful to imagine the theoretically ideal polygenic index that could result from an infinitely large GWAS. In the paper, we call the predictor that would result from this ideal GWAS the "additive SNP factor." The actual polygenic indexes that exist in the world are "noisy" measures of, and therefore only proxies for, this additive SNP factor. The signal-to-noise ratio of a polygenic index—i.e., the extent to which it reflects the additive SNP factor—is determined by the sample size of the GWAS from which the polygenic index is constructed (a larger GWAS leads to less noise and therefore a higher signal-to-noise ratio). The fact that the polygenic index is noisy distorts the results of most analyses that use the polygenic index (relative to what the results would be with the ideal predictor). These distortions can lead researchers to reach incorrect conclusions. For example, in an analysis of how genes and environments interact in influencing some outcome, the noise in the polygenic index will usually cause a researcher to underestimate how strongly genes and environments interact.

Moreover, as discussed in FAQ 1.8, there are many reasons why two polygenic indexes for the same outcome could differ from each other, including differences in the GWAS that the polygenic index is based on and different methods for constructing the polygenic index. Many of these differences among GWASs produce differences in the signal-to-noise ratios of their resulting polygenic indexes. Two studies using polygenic indexes with different signal-to-noise ratios will, in turn, have results that are distorted to differing degrees, reducing comparability of results across studies that use the polygenic indexes.

The "measurement-error-corrected estimator" we derive in the paper enables researchers to conduct analyses *without* the distortion that comes from the noise. It works because we (often) have a good estimate of how much noise a given polygenic index has. We can use that information to calculate what the results of an analysis would have been if the polygenic index had no noise. The estimator improves comparability of results across papers because it avoids the distortions in results that arise from having a noisy polygenic index. Rather than being distorted to different degrees, two studies using polygenic indexes with different signal-to-noise ratios that use our estimator will both have undistorted results. We have made available the software for this estimator. We will maintain and provide user support for this software.

Moreover, across all the polygenic indexes and across all the datasets participating in the Repository, we constructed the polygenic indexes in a uniform way. To the extent that future studies use the polygenic indexes from the Repository, their results will therefore be more comparable.

#### 2.5. What is in the User Guide that accompanies the Repository?

Along with the polygenic indexes, we have distributed to the participating datasets a User Guide. Data providers will distribute this User Guide to researchers as part of the Repository. The User Guide contains technical details about the construction of the polygenic indexes, as well as details about data and software availability. It also describes a set of key interpretational considerations that researchers should keep in mind when analyzing polygenic indexes. These include when to use a single-trait versus multi-trait polygenic index (see FAQ 2.1) and reasons why associations between a polygenic index and an outcome generally cannot be interpreted as causal (see FAQ 1.5). Finally, the User Guide contains a discussion of six "interpretational considerations" that we urge researchers who use polygenic indexes to consider as part of the responsible conduct and communication of their research (see FAQ 3.7).

#### 2.6. Who can access the Repository polygenic indexes, and how?

Researchers can access the Repository polygenic indexes through the data access procedures for each of the datasets participating in the Repository. These are summarized in the Supplementary Note of the paper. Typically, data providers require researchers to submit a brief a description of the planned research and to sign a Data Use Agreement. The Data Use Agreement usually requires researchers to agree to protect the confidentiality of individuals in the dataset and, to that end, to analyze the data on computers that satisfy certain security protocols.

We provided the polygenic indexes we created to the 11 datasets participating in the Repository, so that the data providers can distribute them to users of the datasets. We designed the Repository this way for three reasons (corresponding to problems #1, #2, and #3 in FAQ 1.9; problem #4 is addressed by using a consistent methodology for constructing the polygenic indexes). First, because we are making available the polygenic indexes (rather than the GWAS results from which they are constructed), researchers do not need to spend time constructing the polygenic indexes from GWAS results. Second, for many outcomes, the polygenic indexes we construct are based on more data than are in the largest previously published GWAS. Because the Repository polygenic indexes for those outcomes are based on more data, they are more accurate (i.e., predictive) than polygenic indexes that could be constructed based only on publicly available GWAS results. Third, we tailored the polygenic indexes we constructed to each of the 11 datasets. Specifically, we ensured that for a given dataset, its polygenic indexes were *not* based on GWAS results that included that dataset (which would have led to "sample overlap" that would make it problematic to use the polygenic index with that dataset).

##### 2.7. How will the Repository be updated?

We plan to update the Repository regularly as new GWAS are published or new data become available in which we can conduct our own GWAS. The updates will increase the predictive power of polygenic indexes already in the Repository, as well as expand the set of outcomes for which polygenic indexes are available. We also expect to include additional datasets whose stewards want to participate in the Repository and make their data broadly available to the research community.

#### 3. Ethical and social implications of the study

##### 3.1. Do GWAS or the polygenic indexes they produce identify the gene—or genes—"for" a particular outcome?

No. GWAS of complex outcomes identify *many* SNPs that are associated with an outcome like height or educational attainment. Although it was once believed that scientists would discover numerous strong one-to-one associations between specific genes and outcomes, we have known for a number of years that the vast majority of human traits and other outcomes are complex and are influenced by thousands of genes, each of which alone tends to have a small influence on the relevant outcome.

Furthermore, many complex outcomes are also influenced by parts of the genome that are not genes at all but instead serve to regulate genes (e.g., influencing when a gene is turned on or off). Genes typically contain many SNPs (often dozens or hundreds, in some cases thousands), and there are even more SNPs outside of genes than inside genes. Complex outcomes are often influenced by millions of SNPs.

Although the GWAS that produced the polygenic indexes included in the Repository did find several SNPs that are associated with particular outcomes, we believe that characterizing these as "genes for X"—or, more accurately—"SNPs for X" (e.g., educational attainment, height) is still likely to mislead, for many reasons, and we urge researchers and reporters to avoid this usage.

As an example, consider the outcome of educational attainment. First, most of the variation in people's educational attainment is accounted for by social and other environmental factors, not by additive genetic effects (See FAQ 3.3). “Genes for educational attainment” might be read to imply, incorrectly, that genes are the strongest predictor of variation in educational attainment.

Second, the SNPs that are associated with educational attainment are also associated with many other things. These SNPs are no more “for” educational attainment than for the other outcomes with which they are associated.

Third, the “predictive” power (see FAQ 1.6) of each individual SNP that we identify is very small. Our previous work (Lee et al., 2018) has shown that genetic associations with educational attainment are comprised of thousands, or even millions, of SNPs, each of which has a tiny effect size. Each SNP is therefore weakly associated with, rather than a strong influence on, educational attainment. “Genes for educational attainment” might misleadingly imply the latter.

Fourth, environmental factors can increase or decrease the impact of specific SNPs (see FAQ 3.3). Put differently, even if a SNP is associated with higher or lower levels of educational attainment *on average*, it may have a much larger or smaller effect depending on environmental conditions. Indeed, in our most recent GWAS of educational attainment (Lee et al., 2018) and elsewhere, we report exploratory analyses that provide evidence of such gene-environment interactions. Educational attainment couldn’t even exist as a meaningful object of measurement if we didn’t have schools, and having schools introduces societal mechanisms that influence who goes to them. Accordingly, genetic associations with educational attainment necessarily will be mediated by societal systems and therefore genetic variation should often be expected to interact with environmental factors when it influences social phenomena, such as educational attainment. “Genes for educational attainment” suggests a stability in the relationship between these genes and the outcome of educational attainment that does not exist.

Finally, SNPs do not affect educational attainment directly. As described in our previous work (Lee et al., 2018), the genes identified as associated with educational attainment tend to be especially active in the brain and involved in neural development and neuron-to-neuron communication. The “predictive” power (see FAQ 1.6) of SNPs on educational attainment may therefore be the result of a long process starting with brain development, followed by the emergence of particular psychological traits (e.g., cognitive abilities and personality). These traits may then lead to behavioral tendencies as well as experiences and treatment by parents, peers, and teachers. All of these factors may additionally interact with the environment in which a person lives. Eventually these traits, behaviors, and experiences may influence (but not completely determine) educational attainment.

#### 3.2. Do polygenic indexes show that these outcomes are determined, or fixed, at conception?

Absolutely not. Social and other environmental factors account for most variation in most of the outcomes for which the Repository contains polygenic indexes. But even if it were true that genetic factors accounted for *all* of the differences among individuals in an outcome, it would *still* not follow that an individual’s outcome is “determined” at conception. There are at least three reasons for this.

First, some genetic effects may operate through environmental channels (Jencks, 1980). Again, consider educational attainment as an example. Suppose—hypothetically— that some of the SNPs in the index help students to memorize and, as a result, to become better at taking tests that rely on memorization. In this example, changes to the intermediate environmental channels—the type of tests administered in schools—could have large effects on individuals' educational attainment, even though individuals' genome would not have changed. Certain SNPs may not be associated with educational attainment *at all* if schools did not use tests that rely on memorization. More generally, the polygenic index for educational attainment in the Repository might be less predictive if the education system were organized differently than it is at present (see also FAQ 3.3).

Second, even if the genetic associations with educational attainment operated entirely through non-environmental mechanisms that are difficult to modify (such as direct influences on the formation of neurons in the brain and the biochemical interactions among them), there could still exist powerful environmental interventions that could change the genetic relationships. In a famous example suggested by the economist Arthur Goldberger, even if all variation in unaided eyesight were due to genes, there would still be enormous benefits from introducing eyeglasses (Goldberger, 1979). Similarly, policies such as a required minimum number of years of education and dedicated resources for individuals with learning disabilities can increase educational attainment in the entire population and/or reduce differences among individuals.

Third, even if the genetic effects on an outcome were not influenced by changes in the environment, those environmental changes themselves could still have a major impact on the outcome in the population as a whole. For example, if young children were given more nutritious diets, then everyone's school performance might improve, and college graduation rates might increase. Or consider the outcome of height: 80%-90% of the variation across individuals in height is due to genetic factors. Yet the current generation of people is much taller than past generations due to changes in the environment such as improved nutrition.

##### 3.3. Can the polygenic indexes from the Repository be used to accurately predict a particular person's outcomes?

No. While the "predictive" power (see FAQ 1.6) of our polygenic indexes makes most of them useful in research for some purposes (see FAQ 2.3), these polygenic indexes *fail to predict* the majority of variation across individuals. Even for height—the outcome for which our polygenic index has the greatest predictive power—the index fails to predict 70% of the variation.

Indeed, an important message of a number of our earlier papers is that DNA does *not* "determine" an individual's behavioral and social outcomes, for at least four reasons: First, in the environments in which the outcomes have been measured, other studies have estimated that the additive effects of SNPs will only ever account (even with arbitrarily large samples used to construct polygenic indexes) for a minority of the variation across individuals in the outcomes we study. For example, we estimate that the theoretical upper bound for additive effects of SNPs would account for 46% of the variation in height, 24% in body mass index, 20% in age at first menses, and less than 10% for most of the social/behavioral outcomes we study. So even a hypothetical polygenic index that perfectly reflects the additive SNP factor (see FAQ 2.4) could only explain a small fraction of the variation across individuals. Second, *today's* polygenic indexes are *not* perfect; they are only able to predict a fraction of that already small fraction of cross-sectional predictive power. Third, since SNPs matter more or less depending on environmental context (see FAQ 3.2), a polygenic index might be less (or more) predictive for individuals in some environments than for individuals in others. Finally, and similarly, polygenic predictions only hold for as long as the environment in which they were developed remains substantially the same.

To illustrate these final two reasons, consider the example of educational attainment (for which we have included a polygenic index in the Repository and on which we have done previous research): if the pedagogy underlying the educational system in which the GWAS that produced the polygenic index was conducted is substantially different than the pedagogy of the *different population* to which that polygenic index is being applied, the polygenic index may be less (or, conceivably, more) predictive in this second population (for an example, see FAQ 3.2). The same is true if the polygenic index is applied to the same population, but at a *later time* when the pedagogy has changed substantially. Just as eyeglasses allow those genetically predisposed to poor vision to have nearly perfect vision, innovations in education (say, an innovation that makes education irresistibly engaging, thus mitigating the risk to those with SNPs associated with lower ability to pay attention or maintain self-control) might result in those with lower polygenic indexes now achieving just as much education, on average, as those with higher polygenic indexes.

As sample sizes for GWAS continue to grow, it will likely be possible to construct polygenic indexes for many outcomes whose predictive power comes closer to the total amount of variation that is theoretically predictable from additive effects of common SNPs for those outcomes (the upper bounds given above). Even these levels of predictive power would pale in comparison to some other scientific predictors. For example, professional weather forecasts correctly predict about 95% of the variation in day-to-day temperatures. Weather forecasters are therefore vastly more accurate forecasters than social science geneticists will ever be.

Note: Polygenic indexes created by GWASs are increasingly used by commercial and research direct-to-consumer platforms to predict individual outcomes. We recognize that returning individual genomic "results" can be a fun way to engage people in research and other projects and has at least the theoretical potential to stoke their interest in, and educate them about, genomics and how genes and environments interact. But it is important that participants/users understand that, at present, most of these individual results, including all social and behavioral outcomes, are not *meaningful* predictions (in the sense that they generally have very little predictive power at the individual level). Failure to make this point clear risks sowing confusion and undermining trust in genetics research.

##### 3.4. Can the polygenic indexes accurately be used for research studies in non-European-ancestry populations?

No. We constructed polygenic indexes only for individuals classified as "European ancestry." (The precise definition of "European ancestry" differs in different datasets, but it usually means that a person's pattern of genetic variation across the genome is statistically close to the average pattern from a "reference sample" for some European country. The reference samples used by geneticists are based on samples of people who live in the European country today and whose recent ancestors also lived in that country.) Therefore, the Polygenic Index Repository only includes polygenic indexes for these individuals.

The main reason we only constructed polygenic indexes for these individuals is that the polygenic indexes are likely to be much less predictive—and hence much less useful—in a sample of people of non-European ancestries. That is because our original GWAS data was obtained from samples of people with European-ancestry, and GWAS results have been found to have only limited portability across ancestries (Belsky et al., 2013; Domingue et al., 2015, 2017; Martin et al., 2017; Vassos et al., 2017). There are a number of reasons for the limited portability. For one thing, the set of SNPs that are associated with an outcome in people of European ancestries is unlikely to overlap closely with the set of SNPs associated with the outcome in people of non-European ancestries. And even if a given SNP is associated in both ancestry groups, the effect size—in other words, the strength of the association—will almost surely differ. This is primarily because linkage disequilibrium (LD) patterns (i.e., the correlation structure of the genome) vary by ancestry. This means that some SNP may be associated with the outcome because the SNP is in LD (i.e., correlated) with a SNP elsewhere in the genome that causally affects education (see FAQ 1.5). If the strength of the correlation is greater in one ancestry group than in another, then the size of the association will be larger in that ancestry group. Moreover, even if LD patterns were similar in each ancestry group, the association may differ in different groups because environmental conditions differ (see FAQ 1.6). The fact that there are differences across ancestry groups in the set of associated SNPs and their effect sizes means that the weights for constructing polygenic indexes in European-ancestry individuals (FAQ 1.3) would be the "wrong" weights for non-European-ancestry individuals. For a more extensive, excellent discussion of these and related issues, see Graham Coop's blog post, "Polygenic scores and tea drinking."

Unfortunately, this attenuation of predictive power means that for non-European-ancestry populations, many of the benefits of having a polygenic index available will have to wait until large GWAS studies are conducted using samples from these populations. (Currently, most large genotyped samples are of European ancestries.) We intend that future versions of the Polygenic Index Repository will include polygenic indexes for non-European-ancestry populations, once it becomes possible to produce polygenic indexes with adequate predictive power. We believe that the relative scarcity of polygenic indexes that can be used for research that focuses on non-European ancestry groups is a disparity that should be rapidly eliminated by prioritizing GWAS studies that focus on non-European populations.

#### 3.5. Would it be appropriate to use the Repository social and behavioral polygenic indexes in policy or practice?

No. We reiterate that polygenic indexes are poor predictors of social and behavioral outcomes (see FAQs 2.3 and 3.3). Their *incremental* predictive power over and above other, non-genetic predictors that are already used is even smaller than a polygenic index's predictive power on its own. Moreover, the predictive power of the polygenic indexes for social and behavioral outcomes depends on the environment in which the GWAS participants live (FAQ 3.3). Thus, enshrining polygenic indexes in policy risks basing policy (which can be difficult to change) on weak predictions that could become even weaker or nonexistent as the environment changes. Furthermore, the polygenic indexes can operate through environmental channels (FAQ 3.2). Allocating resources based on polygenic indexes could therefore exacerbate inequalities that were originally due to environmental disparities (a similar risk to that of other biased algorithms that bake in pre-existing discrimination). Using polygenic indexes in order to prioritize giving resources to individuals who are already advantaged would further limit the opportunities of individuals who are disadvantaged, which would be ethically inappropriate. Finally, even if polygenic indexes were used to offer additional resources to disadvantaged individuals, any small potential benefits of using such weak individual predictors would almost certainly be offset by the risk of stigmatization and by the fact that this technology is currently only accessible to people of European ancestries (FAQ 3.4). For all these reasons, we are deeply skeptical that the Repository social and behavioral polygenic indexes have any appropriate role to play in policy now or in the foreseeable future.

##### 3.6. Could research on polygenic indexes lead to discrimination against, or stigmatization of, people with higher or lower polygenic indexes for certain outcomes? If so, why facilitate the spread of polygenic indexes?

Unfortunately, like a great deal of research—including, for instance, research identifying genomic variation associated with increased cancer risk—the results can be misunderstood and misapplied. This includes being used to discriminate against those with higher or lower polygenic indexes for certain outcomes (e.g., in insurance markets). Nevertheless, for a variety of reasons, in this instance, we do not think that the best response to the possibility that useful knowledge could be misused is to refrain from producing the knowledge. Moreover, many researchers already have access to and use polygenic indexes; against this background, the Repository helps ensure that a much wider array of researchers have the same opportunity to access and probe these research tools, and also that the polygenic indexes themselves will be more accurate. Here, we briefly discuss some of the broad potential benefits of this research. We then describe what we see as our ethical duty as researchers conducting this work.

First, one benefit of conducting social-science genetics research in ever larger samples is that doing so allows us to correct the scientific record. An important theme in our earlier work has been to point out that most existing studies in social-science genetics that report genetic associations with behavioral outcomes have serious methodological limitations, fail to replicate, and are likely to be false-positive findings (Benjamin et al., 2012; Chabris et al., 2012, 2015). This same point was made in an editorial in *Behavior Genetics* (the leading journal for the genetics of behavioral outcomes), which stated that "it now seems likely that many of the published [behavior genetics] findings of the last decade are wrong or misleading and have not contributed to real advances in knowledge" (Hewitt, 2012). One of the most important reasons why earlier work has generated unreliable results is that the sample sizes were far too small, given that the true effects of individual SNPs on behavioral outcomes are tiny. Pre-existing claims of genetic associations with complex social-science outcomes have reported widely varying effect sizes, many of them purporting to "predict" as much of the variation across individuals as do the polygenic indexes we construct in this paper that aggregate the effects of millions of SNPs.

Second, behavioral genetics research also has the potential to correct the *social* record and thereby to help *combat* discrimination and stigmatization. For instance, overestimating the role of genetics can be damaging, and the present work can help debunk the myth of genetic determinism. By quantifying how various outcomes are predicted by genetic data, we show that for all of the outcomes we study, the genetic data can explain a very small fraction of the variation across individuals (see FAQ 2.3). By clarifying the *limits* of deterministic views of complex outcomes, recent behavioral genetics research—

if communicated responsibly—could make appeals to genetic justifications for discrimination and stigmatization *less* persuasive to the public in the future.

Third, behavioral genetics research has the potential to yield many other benefits, especially as sample sizes continue to increase—as briefly summarized in FAQ 1.9. Foregoing this research necessarily entails foregoing these and any other possible benefits, some of which will likely be the result of serendipity. Indeed, very few of the uses of polygenic indexes were anticipated when they were first proposed (Wray, Goddard and Visscher, 2007).

In sum, we agree with the U.K. Nuffield Council on Bioethics, which concluded in a report (Nuffield Council on Bioethics, 2002, p114) that "research in behavioural genetics has the potential to advance our understanding of human behaviour and that the research can therefore be justified," but that "researchers and those who report research have a duty to communicate findings in a responsible manner" (see FAQ 3.7).

#### 3.7. What have you done to mitigate the risks of research using Repository polygenic indexes?

In our view, the responsible behavioral genetics research called for by the Nuffield Council on Bioethics (see FAQ 3.6) includes sound methodology and analysis of data (e.g., only conducting analyses that are adequately powered and, when feasible, preregistering power calculations and planned analyses); a commitment to publish all results, including any negative results; and transparent, complete reporting of methodology and findings in publications, presentations, and communications with the media and the public. A critical aspect of the latter is particular vigilance regarding what research results do—and do not—show, and how polygenic indexes can—and cannot—be appropriately used. In an effort to reduce the risk that its results might be misinterpreted by readers, misreported by the media, or misused, the SSGAC has developed and publicly posted FAQs like this document with every major paper it has published since its first paper in 2013.

In addition, the SSGAC will require researchers who download the SNP weights for constructing polygenic indexes to agree to Terms of Service. Among the many terms that we require researchers to agree to, we highlight two here:

> I agree to conduct research that strictly adheres to the principles articulated by the American Society of Human Genetics (ASHG) position statement: "ASHG Denounces Attempts to Link Genetics and Racial Supremacy." (See also International Genetic Epidemiological Society Statement on Racism and Genetic Epidemiology.) **In particular, I will not use these data to make comparisons across ancestral groups.** Such comparisons could animate biological conceptualizations of racial superiority. In addition, such comparisons are usually scientifically confounded due to the effects of linkage disequilibrium, gene-environment correlation, gene-environment interactions, and other methodological problems.
>
> I have read the principles articulated by the ASHG with respect to "Advancing Diverse Participation in Research with Special Consideration for Vulnerable Populations". I agree to adhere to the principles articulated in the final two sections of this statement, "In the Conduct

of Research with Vulnerable Populations, Researchers Must Address Concerns that Participation May Lead to Group Harm" and "The Benefits of Research Participation Are Profound, Yet the Potential Danger that Unethical Application of Genetics Might Stigmatize, Discriminate against, or Persecute Vulnerable Populations Persists."

These Terms of Service stem from the observation that SNP associations are not necessarily causal (see FAQ 1.5) and depend on the environment of the individuals included in the GWAS (see FAQ 1.6). Different ancestry groups arise in the population because they became partially separated from each other many generations ago, for example, due to geographic factors or social forces. When two groups are geographically or socially separated, they also face different environments, which not only may have direct effects on certain outcomes (such as disease risk) but may also change the strength of the association between the outcomes and certain SNPs. Therefore, when individuals from two ancestry groups have different average outcomes, it is extremely difficult to identify whether the difference is due to average genetic differences between the groups or to the different environments faced by the groups. For this reason, it is scientifically invalid to make general statements about ancestry group differences based on SNP associations identified in a GWAS. (Also see FAQ 3.2.) The Terms of Service also require users to securely store the data and to immediately report any breach of the Terms.

Finally, we have developed and provided to participating data providers a User Guide to be distributed to researchers who use Repository polygenic indexes (see FAQ 2.5). We will also provide the User Guide to researchers who download the SNP weights. One section of the User Guide discusses six "interpretational considerations" that are likely to arise when conducting research with polygenic indexes and which we urge researchers to seriously consider as a critical part of responsibly conducting and communicating their research. One recurring ethical concern about genetic research is the tendency for its predictive power to become exaggerated in the media and in the public's minds, at the expense of a more nuanced understanding of how genes and environment interact, the importance of environmental influences, and the ability of interventions to improve outcomes. Many of the interpretational considerations we discuss in the User Guide involve how to anticipate and address potential confounds and how to navigate complex questions about causality and ensure responsible communication of causality.

For instance, the User Guide cautions researchers to appreciate and communicate that associations between a polygenic index and an outcome may operate through *environmental* (rather than biological) mechanisms (see FAQs 3.2 and 3.3).
